# OPTIMAL: An OPTimised Imaging Mass cytometry AnaLysis framework for benchmarking segmentation and data exploration

**DOI:** 10.1101/2023.02.21.526083

**Authors:** Bethany Hunter, Ioana Nicorescu, Emma Foster, David McDonald, Gillian Hulme, Andrew Fuller, Amanda Thomson, Thibaut Goldsborough, Catharien M.U. Hilkens, Joaquim Majo, Luke Milross, Andrew Fisher, Peter Bankhead, John Wills, Paul Rees, Andrew Filby, George Merces

## Abstract

Analysis of Imaging Mass Cytometry (IMC) data and other low-resolution multiplexed tissue imaging technologies is often confounded by poor single cell segmentation and sub-optimal approaches for data visualisation and exploration. This can lead to inaccurate identification of cell phenotypes, states or spatial relationships compared to reference data from single cell suspension technologies. To this end we have developed the “OPTIMAL” framework to benchmark any approaches for cell segmentation, parameter transformation, batch effect correction, data visualisation/clustering and spatial neighbourhood analysis. Using a panel of 27 metal-tagged antibodies recognising well characterised phenotypic and functional markers to stain the same FFPE human tonsil sample Tissue Microarray (TMA) over 12 temporally distinct batches we tested several cell segmentation models, a range of different *arcsinh* cofactor parameter transformation values, five different dimensionality reduction algorithms and two clustering methods. Finally we assessed the optimal approach for performing neighbourhood analysis. We found that single cell segmentation was improved by the use of an Ilastik-derived probability map but that issues with poor segmentation were only really evident after clustering and cell type/state identification and not always evident when using “classical” bi-variate data display techniques. The optimal *arcsinh* cofactor for parameter transformation was 1 as it maximised the statistical separation between negative and positive signal distributions and a simple Z-score normalisation step after *arcsinh* transformation eliminated batch effects. Of the five different dimensionality reduction approaches tested, PacMap gave the best data structure with FLOWSOM clustering out-performing Phenograph in terms of cell type identification. We also found that neighbourhood analysis was influenced by the method used for finding neighbouring cells with a “disc” pixel expansion outperforming a “bounding box” approach combined with the need for filtering objects based on size and image-edge location. Importantly OPTIMAL can be used to assess and integrate with any existing approach to IMC data analysis and, as it creates .FCS files from the segmentation output, allows for single cell exploration to be conducted using a wide variety of accessible software and algorithms familiar to conventional flow cytometrists.

## Introduction

Single cell suspension technologies have now advanced to the point where we can measure thousands of parameters on millions of individual cells at truly “multi-omic” scale. However the digestion and destruction of tissues to liberate single cells can affect the native cellular states as well as obliterating all spatial context. As such, “space” very much remains the “final frontier” with multiplexed single cell tissue imaging traditionally lagging behind suspension technologies due to previously insurmountable technical issues around how many signals can be measured on the same sample/slide/section. Over the past few years these issues have been overcome by the use of cyclical approaches to staining and imaging with fluorescent probes (1,2), or by moving away from fluorescence detection entirely with technologies such as “Multiplexed Ion Beam Imaging” MIBI (3) and Imaging Mass Cytometry (IMC). IMC uses a powerful 1µm laser to raster scan the metal-conjugated antibody stained slide liberating small pieces of tissue for analysis by “Cytometry by time of flight” (CyTOF) technology (4). IMC has several advantages over cyclical fluorescence detection such as no auto fluorescence and no increase in measurement time with an increasing number of signals. It does, however, lack the same image resolution as optical systems (fixed at 10x magnification) due to the 1 µm beam size of the ablating laser. While this is still sufficient to detect individual cells for phenotyping and spatial analysis the low image resolution can present challenges with subsequent data analysis. Unlike suspension technologies, IMC, along with all tissue imaging approaches with single cell resolution, usually requires an additional pre-processing step whereby single cells or objects are identified using an image analysis technique called “segmentation”. Segmentation algorithms are generally based on assessing variance at the pixel level and then using commonalities and differences to group individual pixels together as “super pixels” or “single cell objects” via machine learning approaches (5). It is then possible to derive single object/cell features based on metal intensity (antibody/DNA intercalator), morphometrics (area, circularity etc.) as well as the x and y centroid co-ordinates for every cell within each image. These features can then be used to explore the data using classical single cell analysis approaches such as dimensionality reduction and clustering (6,7). There is generally a need to validate any cell types/states identified within the tissue against reference data derived from tissue digestion and suspension technologies, with usual caveats concerning the effects on cells/markers caused by enzymatic and/or mechanical disaggregation. As such, poor single cell segmentation can have dramatic and confounding effects on accurate cell type/state identification, akin to measuring doublets/aggregates of debris by conventional flow cytometry or scRNAseq. There are a number of published “end to end” pipelines for IMC data analysis (8–12) that utilise open source software for segmentation such as Ilastik (13) and CellProfiler (14,15), as well as StarDist (16) and IMC-specific approaches that utilise deep learning (17). There have also been attempts to use matched fluorescent images of the nuclei using DAPI co-staining to improve segmentation accuracy (18) as well as removing image noise (19,20). Nonetheless, it has been shown that, due to the nature of tissue imaging, simple approaches to single cell segmentation are often highly effective (21). Once single cells have been identified and exploratory features created and assigned, analysis follows an analogous route to suspension technologies with various corrections being applied. This includes a form of isotopic signal spillover correction (22), as well as parameter transformation and batch effect normalisation prior to the use of dimensionality reduction techniques to visualise high parameter data and clustering to identify resident cell types/states. There are a number of existing approaches for visualising and analysing IMC data such as HistoCat (9) and ImaCytE (23), both provide the ability to perform spatial neighbourhood analyses on the cell types and states identified via clustering approaches. However they lack the flexibility to be able to optimise key steps and parameters of the pipeline in an easy and accessible manner. Here we present a novel framework we call “OPTIMAL” that provides metrics and benchmarks for each major step of IMC data analysis including segmentation, parameter correction, normalisation and batch effect removal, as well as dimensionality reduction, clustering and spatial analysis. This is not a new analysis pipeline per se, rather an exploration and optimisation of existing approaches that allows for democratised analysis of cellular phenotypes from multiplexed tissue imaging technologies such as IMC; especially as we convert all data to .FCS file format allowing it to be explored in an easy to use, accessible software. To test OPTIMAL we stained, acquired and analysed Tissue Microarrays (TMAs) from the same human tonsil sample over 12 temporally distinct batches using a panel of 27 metal-tagged antibodies and IMC. We then investigated several different cell segmentation approaches based on the previously described Bodenmiller method (10), open source software (Ilastik and CellProfiler), as well as deep learning (CellPose) (24) using cell type cluster “fidelity” as our measure of success using the human tonsil “ground truth” populations known to be identified by our 27 marker panel. Prior to clustering however, we used OPTIMAL to identify the optimum *arcsinh* transformation cofactor to maximise signal resolution and to identify the use of a subsequent Z-score normalisation factor as the best method of batch effect removal. We also identified the most effective dimensionality reduction and visualisation method for IMC data to be PacMap, and FLOWSOM to be the best performing clustering algorithm for finding the expected cell types and states. Finally we developed an approach to optimise spatial neighbourhood analysis that used a more accurate method of finding neighbouring cells than existing approaches and benchmarked this against well-defined cell types and structures in human tonsil. The OPTIMAL framework can be applied to any existing and future IMC data analysis as it provides a set of methods and metrics to empirically assess each stage in any pipeline, moreover, by producing .FCS files from our segmentation output we make exploration of the single cell data highly democratised and not reliant on further expert coding skills.

## Materials and methods

### Tonsil tissue preparation and antigen retrieval

Formalin-fixed paraffin-embedded 2mm human tonsil tissue cores were obtained from the Novopath Tissue Biobank (Royal Victoria Infirmary, Newcastle upon Tyne) and embedded into a 3 core Tissue Miccroarray (TMA). TMA blocks were constructed manually using Medical biopsy punches (PFM Medical, UK). Cores were selected using haematoxylin and eosin-stained slides to guide suitable areas in the donor blocks. Cores were placed in a paraffin embedding mould, heated to 65°C and embedded in molten wax before cooling to set. 8µm serial sections were cut using HM 325 Rotary Microtome (Fisher Scientific, USA) and mounted onto SuperFrost Plus™ Adhesion slides (Epredia, CAT#10149870).

### Antibody panel design, conjugation and validation by Immuno-Fluorescence

A 27-plex antibody panel was designed to identify the immune, signalling and stromal components in the surrounding microenvironment. All antibodies used in this study were first validated for performance using the chosen single antigen retrieval methods outlined previously (Tris EDTA pH9 “Heat-Induced Epitope Retrieval”, HIER) for IMC using simple two colour immuno-fluorescence (IF). All relevant antibody details are shown in Table S1 including the choice of metal tag based on the relative staining intensity of each marker by IF using the rules of “best practice” for CyTOF panel design (25). Unless stated otherwise, following verification of staining pattern and performance quality, approved antibodies were subject to lanthanide metal conjugation using a Maxpar X8 metal conjugation kit following manufacturer’s protocol (Standard Biotools, CAT#201300). Antibodies conjugated to platinum isotopes 194 Pt and 198 Pt were conjugated as described in Mei *et al*. 2015 (26). Conjugated antibodies were validated by firstly checking the recovery of antibody post-conjugation. Secondly, we checked for successful metal conjugation by binding the antibody to iridium labelled antibody capture beads AbC™ Total Antibody Compensation Beads (Thermo Fisher, USA, CAT#A10513) and acquiring on a Helios system (Standard Bio-tools, USA) in suspension sample-delivery mode. Finally, we checked that the antibody had refolded and retained the ability to recognise antigen by using the post-conjugation antibody in either a two layer IF with a fluorescently labelled secondary antibody recognising the primary antibody species or directly by IMC using the Hyperion imaging module (Standard Bio-Tool) connected to the Helios. A gallery of IMC-derived grey scale images for each stain (Ab and DNA) is shown in Figure S1B. Test tissue sections were then stained with the 27 marker antibody cocktail as outlined in Table S1.

### Hyperion (IMC) set up, quality control (QC) and sample acquisition

Prior to each sample acquisition, the Hyperion Tissue Imager was calibrated and rigorously quality controlled to achieve reproducible sensitivity based on the detection of 175Lutetium. Briefly, a stable plasma was allowed to develop prior to ablation of a single multi-element-coated “tuning slide” (Standard Biotools). During this ablation, performance was standardised to an acceptable range by optimising system parameters using the manufacturer’s “auto tune” application or by manual optimisation of XY settings whilst monitoring 175Lutetium dual counts. After system tuning, tonsil sections were loaded onto the Hyperion system in order to create Epi-fluorescence panorama images of the entire tissue surface to guide region of interest (ROI) selection. Two ROIs of approx. 500µm^2^ encompassing lymphoid follicles and surrounding structural cells were selected for ablation per batch run. Small regions of tonsil tissue were first targeted to ensure complete ablation of tissue during the laser shot with ablation energies adjusted to achieve this where required. Finally, ablations were performed at 200Hz laser frequency to create a resultant MCD file containing all data from ROIs. Correction of ‘spillover’ between isotopes was performed as per the protocol described at Spill over correction | Analysis workflow for IMC data (bodenmillergroup.github.io) without deviation (22).

### Image QC and export

MCD files from the Hyperion system were opened using MCD™ Viewer v1.0.560.6 software (standard bio-tools) in order to perform a qualitative, visual QC of the staining intensity and pattern with the initial IF images as a benchmark. Pixel display values (max/min and gamma) were set to optimise the display of the 16 bit pixel range from the Hyperion detector (0 – 65,535) to the 8-bit display (0 - 255). Multi-pseudo coloured, overlaid images were built for figures with a scale bar included and the option to export as an 8-bit TIFF with “burn in” was used. The digital magnification was also set to “1x” so that each signal was carefully balanced for display purposes to aid qualitative visual interpretation. All images were exported as 16-bit single multi-level TIFFs using the “export” function from the “file” menu. For ease of use, all open collection channels from the experimental acquisition template (in this case, 60, including several “Blank” channels for QC purposes) from all ROIs were left ticked and any image/channel removal was dealt with later in the analysis. This avoided having to repeatedly deselect image channels for each ROI in the MCD file. These multi-level 16-bit TIFF images were then input in to our pipeline as shown in Figure 1A.

**Figure 1:**
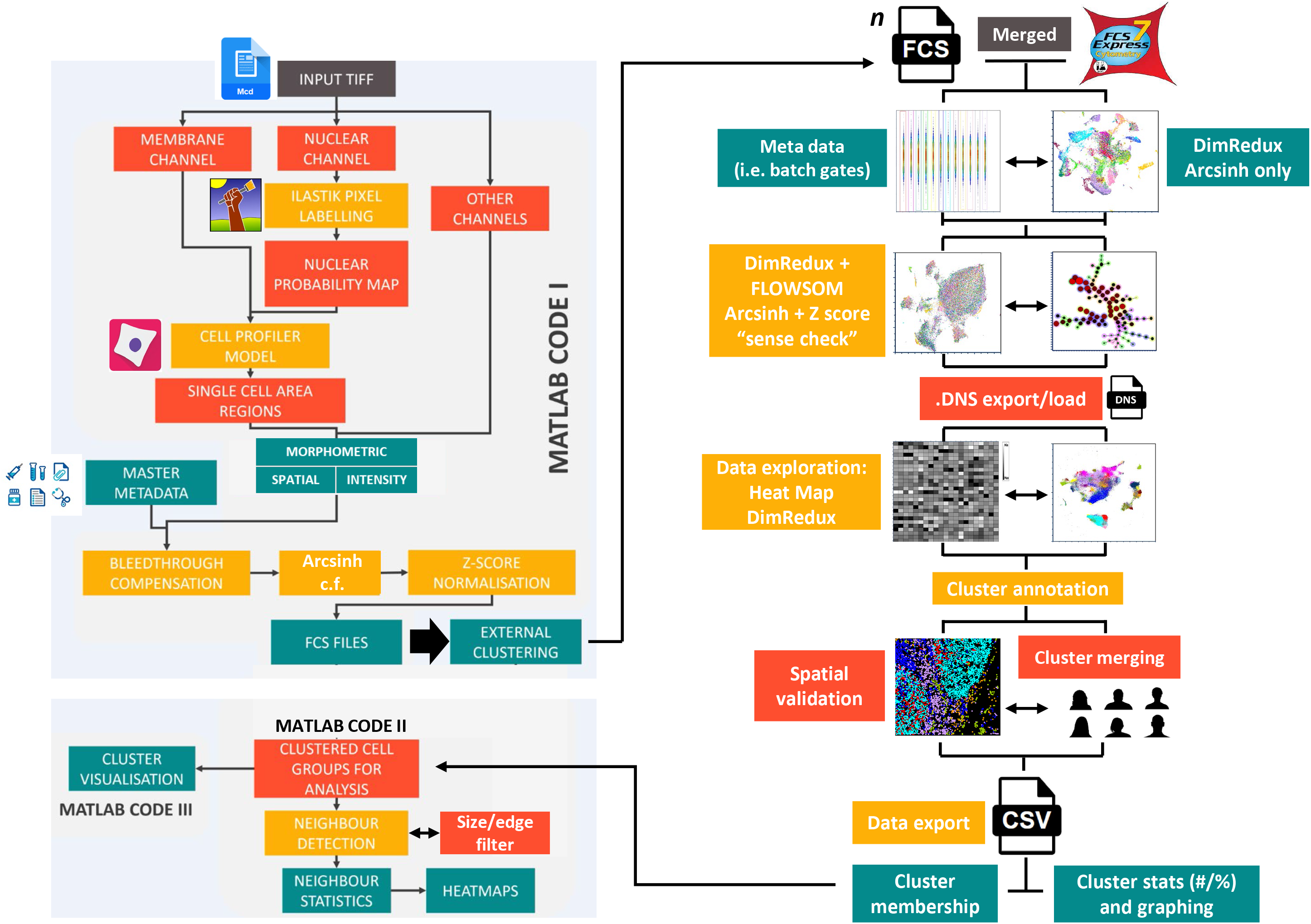
Summary workflow diagram for the OPTIMAL analysis pipeline. Briefly, input multi-level TIFF images created from MCD file export were segmented using a combination of the nuclear channel via an Ilastik random forest pixel classification to generate a probability map (p-map). Then single or multiple membrane channel images were used in conjunction with the p map to create single cell objects via CellProfiler and to create features sets based on intensity, morphometry and spatial location. Additional metadata was also incorporated at this stage and included batch number. Metal intensity values were corrected for bleed through and two sets of subsequent parameters were created i) *arcsinh* c.f. 1 transformed and (ii) a subsequent Z-score normalised set. This data matrix was converted in to .FCS file format (collectively MATLAB code I) and analysed using FCS Express to explore things like batch effect normalisation, dimensionality reduction (DimRedux) for visualisation and clustering via FLOWSOM and heat map creation/interpretation. Cluster annotation was performed using a combination of hierarchical consensus merging and expert *a priori* knowledge combined with a basic spatial validation using x/y centroid plots. Once all cells in all images were assigned a cluster membership, a master .CSV file was created with the minimal necessary metadata to perform a neighbourhood interaction analysis using MATLAB code II with the results visualised using MATLAB code III.

### Cell segmentation, feature extraction, parameter correction/normalisation and FCS file creation

Cell segmentation was based on the previously described method of Zantonelli *et al.* (10), that uses a combination of random forest pixel classification using Ilastik (Version 1.3.2 or later) (13) and helps to inform single cell segmentation and feature extraction using CellProfiler (version 4 or later) (15). Ilastik models were created to distinguish nuclear vs non-nuclear pixels based on partial labelling of multiplexed images of tonsil tissue or “Vero” monolayer cell culture. An additional run using unprocessed input Iridium 193 (DNA channel 51) was also trialled for comparison to Ilastik processing. Output nuclear probability maps were input into CellProfiler, enabling instance segmentation of cell nuclei, which were subsequently used as seeds for cell segmentation. Cell boundaries were determined using a seeded watershed algorithm either to EPCAM (channel 29) signal, or a maximum intensity projection of multiple membrane markers (see supplemental notes, section S2.5.3). Following cell segmentation, individual cells were measured for mean intensity in each of the labelled channels. Intensity measurements were compensated for spillover according to a previously described approach (22). *Arcsinh* transformation was trialled using values from 0.1 to 120 using the Fisher Discrimination Ratio (Rd) to determine the optimum value for positive vs negative signal distribution (see supplemental notes). Following optimisation, *arcsinh* transformation was applied to all experimental datasets with a value of 1. Additionally, a second set of metal intensity parameters were derived whereby an additional subsequent Z-score normalisation step was applied to the previously *arcsinh* cofactor (c.f.) 1 transformed values. This additional Z-score normalisation was used to remove batch effect as well as to normalise marker intensities relative to one another for subsequent optimised heat map display. At this stage, any additional metadata was included in the files such as batch number in order to be an accessible and plot-able parameter for subsequent analysis. Final matrix data was converted to .FCS file format within the MATLAB pipeline for preparing for clustering. More details on our method and tests can be found in the supplemental method section (S2.5 and also visual guide at the end of the supplemental notes).

### Visualisation, clustering and exploration of single cell IMC data

For this study we used the commercially available “FCSExpress” software for all single cell data analysis (Version 7.14.0020 or later, Denovo software by Dotmatics, USA). More extensive information can be found in supplemental notes. Briefly, the FCS files created from the segmentation pipeline shown in Figure 1A for all 24 tonsil images across 12 batches were loaded as a single merged file. We then created a set of batch gates using a simple density plot of batch number (x axis) versus Iridium signal (Z normalised parameter version) and selected contrasting colours for each and used the “pipelines” function within the “tools” menu to create UMAP parameters derived from the *arcsinh* c.f. 1 transformed and Z-score normalised antibody channels. In addition we created UMAP parameters from the *arcsinh* only versions in order to verify for the presence of batch effect and subsequent correction by Z score normalisation (see Figure 1B). Next we used the same fully transformed and corrected parameters for FLOWSOM clustering using the default settings (see supplemental notes S2.6.5) with a merging of the 100 SOMs to 30 consensus clusters (cSOMs) based on hierarchical clustering and created a set of uniquely coloured cSOM gates using the “plate heat map” and “well gates” function. We also created a PacMap dimensionality reduction plot using the same parameters using the Python interface within the FCS Express pipeline module (see supplemental notes S2.6.4). At this stage, after validation of results, we exported the data as both a single merged and a set of individual “Data stream” (.DNS) files. These contained the new clustering and visualisation parameters (SOMS and PacMap x/y co-ordinates) as well as all the original .FCS file metadata but in a smaller, compressed and easier to work with file format. Next we loaded the merged .DNS file in to a new incidence of FCS Express and conducted a much more extensive analysis of the data by (re)- constructing all the necessary meta-data and SOM gates as well as a heat map of transformed and normalised antibody-derived signals (rows) versus the 30 cSOMs (columns). The median values were normalised by column (cluster) to aid interpretation of the heat map on a per cluster per marker basis. Using the information on the panel in table S1, we assigned broad cluster identities to these SOMs. We then used simple x/y centroid plots as well as further *a priori* legacy knowledge to manually merge any highly similar clusters with basic spatial verification. Finally we exported the clustered data in two formats. Firstly as a .CSV file only containing the minimum information needed for neighbourhood analysis; namely the ROI/Sample/Image ID, the cell ID within the ROI and the final cluster assignment for each cell. The second export file contained the ROI/Sample/Image name, the total cell count in the ROI and the percentage and/or cell number in each of the final cSOMs. The latter step can easily be performed using the individual .DNS files for each ROI/Sample/Image and using the FCS Express “batch export” function (see supplemental notes section S2.6).

### Neighbourhood analysis

Neighbourhood analysis was performed with slight adaptations to the method outlined by HISTOCAT (9) and ImaCytE (23). Cell identities were determined by cluster analysis and saved, along with all other cell data, into a single large .CSV file. A separate excel file was used to store the cell type information as a biologically relevant name. Cell masks, stored from the cell segmentation stage, were input and cell identity transposed onto this data. Each cell was assessed for the number of unique cell identities within a pixel-defined threshold distance from the cell edge. The original HISTOCAT code was used, in addition to a modification using a “disc” element to determine the nearest-cell neighbours to each start point cell to investigate. We also tested the use of and automated edge cell removal as well as cells of extreme areas (<20 µm^2^ and >200 µm^2^) to account for any possible segmentation errors. The cell identities for analysis were then mapped at random onto the cell masks, according to the number of each cell type identified by clustering for each image. This was repeated to create 100 iterations of randomly organised cell types on the underlying tissue. The interaction between cell types (i.e. the neighbour breakdown by cell type) was compared between these iterations and the original data, to determine if a difference can be identified between the original data and the randomly organised iterations. If differences are detected in the original data compared to a 90% threshold of the random iterations, then a significant difference is listed for that cell type for that image. These positive, neutral, and negative interactions were then collated to create the overall proportion heatmap for the condition (i.e. pathology, region, etc.) ranging from 1 (100% of images showed positive interaction) to -1 (100% of images showed negative interaction). A cluster “occupancy” cut-off percentage value of 0.01 was used for all analyses, however this was unimportant as all final consensus clusters were present in all 24 ROIs.

## Results

### Ilastik-derived nuclear probability maps and a single “pan” membrane signal provides the optimal segmentation of single cells in human tonsil

We began by generating an IMC data set using TMA tissue from serially sectioned FFPE human tonsil tissue that had been stained with a panel of 27 metal tagged antibodies targeting well-characterised cell types/states as detailed in Table S1 over 12 temporally distinct batches (staining and acquisition). In each case ROI selection was designed to capture as much structure as possible, including lymphoid follicles, germinal centres and epithelium in order to provide high likelihood of positive staining for all 27 markers in all ROIs selected (see Figure S1 A and B). Tonsil tissue was also selected for its dense cellularity in order to present a genuine challenge to segmentation but with clear *a priori* knowledge concerning what cell types our panel should find and where. It should be noted that while EPCAM is not biologically expressed across all cells in the Tonsil, due to some degree of non-specific staining, it was judged to show the most uniform and comprehensive ability to identify cell membranes across the entire tonsil tissue, far superior to using combination of tonsil specific markers (see figure S1C) Figure 2A shows a representative ROI from batch 3 with sequential composite images to mark distinct cell types such as T cells (CD3+), B cells (CD79a+) and macrophages (CD68+). Ki67 was included to help denote follicles/germinal centres by virtue of a proliferative signature. The selection of these markers was deliberate as CD3, CD79a and CD68 should all be mutually exclusive and not co-expressed by any single cell. Moreover the spatial location/segregation of several populations that our panel was designed to identify should follow a well-established pattern. As such this provided us a qualitative way to assess the potential signal overlap in each ROI that would likely be due to the dense cellular nature of the human tonsil combined with the lack of Z plane information afforded by IMC (see Figure S2 for CD3/CD79a/DNA composite images for all 24 ROIs). We next sought to test our different segmentation approaches on these images to determine which was optimal. To do this we constructed probability maps (p masks) using the Random Forest pixel classifier within Ilastik using only the DNA (Iridium 193) channel image from either our tonsil TMA tissue or from an embedded “irrelevant” suspension cell line (Vero cells) using two pixel classes; “nuclear” and “background”. Figure 2B (upper panel) shows the Vero and Tonsil-derived probability maps (p-maps). The final step of our segmentation approach was to use the nuclear objects derived from the p-maps as “seeds” to anchor a marker-controlled watershed approach to expand out and delineate the boundaries for each single cell. In this case we compared the use of a single membrane signal (EPCAM) for both models versus using the sum of several membrane signals (tonsil p-map model only) and the cell segmentation boundaries for the same representative ROI are shown in Figure 2B (lower panel). As an example of sub-optimal segmentation we also used an approach that was not based on an Ilastik machine learning model, but instead directly attempted to segment objects within CellProfiler using the DNA (Iridium 193) channel. The segmentation outputs for all 24 tonsil ROIs derived from the Tonsil Ilastik – EPCAM membrane approach are shown in Figure S3 and the same for the Nucleus only model in Figure S4. To provide some quantifiable metric to assess each approach we plotted the intensity of CD3 versus CD79a and looked for double positive (DP) “nonsense cells” and included the total number of objects identified (Figure 2C). For the representative ROI shown in Figure 2B, we noted that the Vero cell p-map identified fewer objects than the Tonsil derived p-map (5039 versus 5839) with the nucleus only approach identifying far fewer (3551). Moreover the frequency of CD3/CD79a DP cells was also similar regardless of the p-map model (tonsil versus Vero) or the approach used to delineate cell boundaries (EPCAM alone versus a multi-marker signal approach) with ∼21% of events within the gates. There was however an increase in the frequency of DP cells in the nucleus only plot (∼29%) but it was surprisingly modest considering the gross under segmentation using this method. Bivariate plots of CD3 versus CD79a on all segmented objects are shown for the Tonsil Ilastik – EPCAM membrane approach in Figure S5 and for the Nucleus only approach in Figure S6 for comparison and again show that there was minimal impact on the percentage of DP cells as a result of clearly sub-optimal segmentation. To provide some basic spatial context we also plotted the x and y centroid values for each segmented object coloured by membership of each gate (B cells, T cells and DP cells, see figure 2D). These data showed very little differences in the arrangement of CD3+, CD79a+ and DP cells between segmentation approaches but did highlight the fact that without the use of a p-map approach, the cells were grossly under segmented. Collectively these data suggested that an effective, yet straightforward approach to cell segmentation is the use of random forest pixel classifier trained on the same or similar sample/tissue type with a single widely expressed membrane marker to delineate cell boundaries. We hypothesised that more profound differences between segmentation methods may be revealed by clustering, however before moving to this stage, we needed to optimise other elements of the data set.

**Figure 2:**
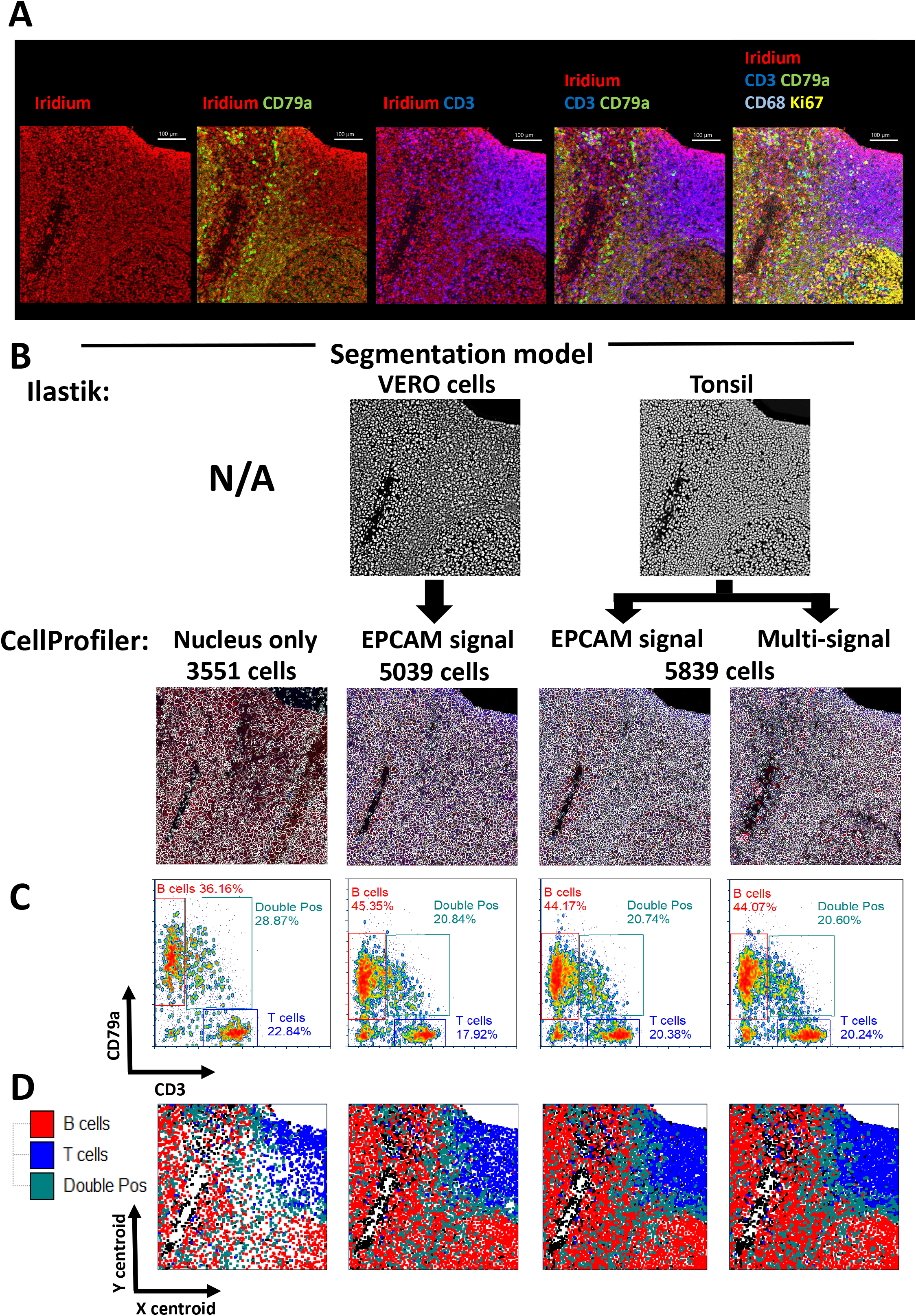
Assessment of segmentation approaches for accurate single cell identification in complex tissues using a 28 parameter (27 antibody) IMC panel on human FFPE tonsil. (A) Multi-parameter pseudo-coloured images from a representative human tonsil ROI with well-separated B and T cell areas. The first image column (left to right) shows DNA staining with Iridium (red pseudo colour), the next column images include CD79a as an overlay (green) with iridium (red), the next shows CD3 (blue) overlaid with iridium (red). The next set of images combine the CD3 (blue), CD79a (green) and iridium (red) as a triple overlay. The final image panel shows the addition of two further parameters, CD68 (teal) and ki67 (yellow). (B) Segmentation maps for the 4 different segmentation models tested in this study showing the same ROI as in A. The upper panels show (where used) the probability map outputs from the indicated Ilastik model (derived from either the same tonsil data set or from Vero cells). The lower panels show the segmentation boundaries generated using CellProfiler alone (far left image) or from the indicated Ilastik p-map. Various approaches to delineate the cell boundary are indicated and include using a single membrane signal (EPCAM) or a combination of multiple markers (multi-signal). (C) Bi-variate single cell level intensity plots for each of the 4 segmentation approaches shown in A-B with CD3 intensity displayed on the x-axis and CD79a intensity displayed on the y-axis. In both cases the *arcsinh* c.f.1 transformed, Z-score normalised values have been used. Gates have been set to quantify the percentage of cells that express CD3 or CD79a alone as well as biologically impossible double positive (DP) cells that may indicate a failure in accurate segmentation. The total number of single cell events are also shown on each plot. (D) x/y cell centroid maps of the same tonsil ROI in A/B/C coloured by the gated population shown in C for each of the 4 individual segmentation approaches coloured as indicated in the legend.

### IMC data structure and batch effect removal benefits from optimal parameter transformation, Z score normalisation and optimal dimensionality reduction approaches

Having determined that the optimal cell segmentation approach we tested used the Tonsil Ilastik p-map combined with watershed detection of the EPCAM membrane boundary, we next wanted to determine the optimal parameters for transformation, batch effect normalisation and multi-parameter data visualisation. We began by evaluating the most suitable cofactor for *arcsinh* transformation of the metal signal parameters. Single cell data variance increases with parameter value meaning that distances at higher (positive) values are less significant than distance from lower (negative) values. This is not suitable for dimensional reduction or clustering algorithms as most assume distances are of equal importance/weight. It is therefore essential to use special scaling formulas to stabilise variance. One of the most effective and widely utilised approaches is the hyperbolic arcsine (*arcsinh*) transformation (27). It is widely used in fluorescence-based flow cytometry and suspension-based mass cytometry (28). The choice of cofactor has a profound influence on the post-transformation data structure and values of between 100 – 150 have been recommended and are widely used for fluorescence-based detection whereas a lower value of 5 is routinely used for mass cytometry in suspension. To our knowledge however, there have been minimal attempts to empirically prove why these values have been used in either technology (29) or, importantly, any attempts to determine what co factor is optimal for IMC data. Values of 5 have been used to simply mirror suspension based mass cytometry (9) or values of 5-15 have been proposed (30). To this end we performed a titration of *arcsinh* c.f. values spanning a range from 150 – 0.1 and used the “fisher discrimination ratio” (Rd), also known as the “Linear Discriminate Analysis” (LDA) (31) to determine the statistical separation between a gated low and high signal distribution (figure 3A) and then created UMAP plots from each c.f. values parameter set. By plotting the *arcsinh* c.f. versus the Rd value we were able to empirically determine that a value of “1” was optimal for achieving the maximal resolution of IMC-derived metal signal parameters (see figure 3B). This was the same for all 28 metal isotope parameters in the panel.

**Figure 3:**
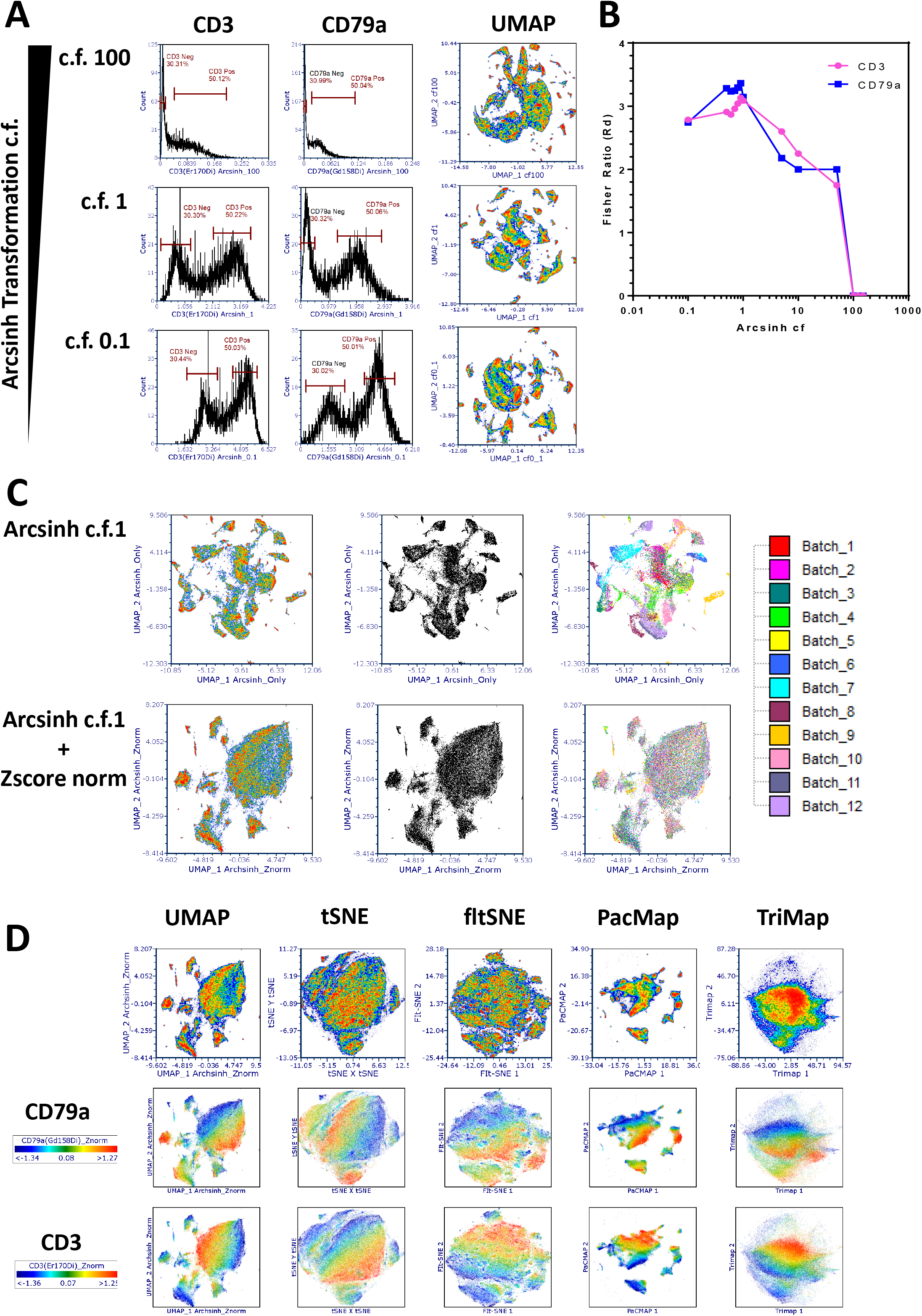
Optimisation of data scaling co-factors, batch effect correction and dimensionality reduction for IMC data analysis. (A) The impact of *arcsinh* cofactor (c.f.) values on parameter/channel resolution. Left panel shows histograms of CD3 expression intensity derived from segmented single cells within the human tonsil tissue with decreasing *arcsinh* c.f. values down the rows (100, 1 and 0.1). The right panels show the same analysis for CD79a expression intensity levels. “Negative” and “positive” gates are set on each plot to derive the population statistics (median and rSD) required to calculate the “Fisher ratio” (Rd) resolution metric (see methods). (B) The graph shows the relationship between Rd (y axis) as a function of *arcsinh* c.f. (x axis) for CD3 and CD79a. (C) Batch effect in data can be eliminated by correct normalisation approaches. UMAP plots of 27 antibody-based parameters of *arcsinh* c.f. 1 transformed data only (upper panels) and *arcsinh* c.f. 1 transformed, Z-score normalised data (lower panels) showing all 109,535 single cells. From left to right, the first UMAP plots are coloured by cell density. The middle UMAP plots are standard black and white dot plots and the third UMAP plots are coloured by batch (see key). (D) The choice of dimensional reduction algorithm impacts on the representation and interpretation of underlying IMC data structures. The indicated dimensionality reduction algorithms were run on the same 27 *arcsinh* c.f.1 Z score normalised parameters as in B. The upper row shows general cell density, the middle row is density weighted by CD79a expression and the lower panel by CD3 expression.

We next sought to address the issues of batch effect normalisation. While every attempt was made to eliminate and control batch effect by using the same donor tonsil tissue across all batches, the same lot of conjugated antibodies, the same person carrying out the staining protocols and a well maintained/QCe’d Hyperion instrument, the nature of working with FFPE tissues often generates significant variation. We began by firstly assessing whether there were batch effects in our data set that could influence the data structure and thus any biological interpretation. By plotting all 109,535 cells derived from the Tonsil-EPCAM segmentation model as a UMAP we could see that our 28 *arcsinh* c.f.1 transformed metal signal parameters (27 antibodies plus iridium) gave us very well structured data. However when we introduced colouration to these events based on batch membership, we could see that the majority of the data structure came from batch effects rather than true underlying biology (see figure 3C upper panels). To attempt to correct for batch effect we tried a number of approaches such as “Batchelor” (32), “Harmony” (33) and “Seurat” (34) however we found that they were not easily compatible with our data files. As such, we found that Z-score normalisation of the *arcsinh* c.f.1 transformed metal signal parameters to be most effective (Z-score normalisation is available in the FCS Express pipelines feature). When we created a UMAP plot using the Z-score normalised version for all 28 of the metal signal parameters, while the global data structure did collapse somewhat, colouration of each event by batch membership revealed an almost complete removal of batch effect (Figure 3C lower panels). We also tested the method of “0-1 scaling” as described by Ashhurst *et al.* (30) but this did not eliminate the batch effect in our data (see Figure S7).

Having established the optimal transformation c.f. and normalised for batch effects, we next wanted to determine if UMAP was indeed the optimal algorithm for presenting the underlying structure of IMC data. To this end we assessed five different dimensionality reduction methods in all cases uses the recommended hyper-parameter settings (see figure 3D upper panel); UMAP (as described previously), fltSNE (35), tSNE (6), PacMap (36) and Tri-Map (37). Typically tSNE is widely to visualise IMC data however it is often the case that it projects very little data structure. There is some argument that UMAP performs better for data with parameters that are poorly resolved and does a better job of projecting both local and global data structures (38). Our data supports this concept as tSNE representation of our IMC data lacked any discernible structure and moreover, density-based overlay of fiducial phenotyping markers such as CD3 (Figure3D middle panel) and CD68 (Figure 3D lower panel) showed very little focus of events expressing these markers in defined areas of the map. The fltSNE algorithm performed as poorly as tSNE with triMAP giving by far the most sub-optimal results. Interestingly though, PacMap performed very well and gave better data structure than UMAP in our hands, with very clear islands with mapping of the fiducial markers to defined areas. As such we decided to use PacMap to visualise our IMC data using the *arcsinh* c.f. 1 and subsequent Z-score normalised metal parameter feature set.

### Suboptimal segmentation has a detrimental impact on the ability to confidently identify all expected tonsil-resident phenotypes using clustering approaches

Although the segmentation approach did not seem to create overtly inferior or superior single cell level data outputs as judged by our simplistic CD3 and CD79a DP “nonsense” cell frequency analysis (Figure 2B and figures S5 and S6), we wanted to assess whether clustering and cell type identification would be more affected. To this end we used the FLOWSOM clustering algorithm (39) to partition the single cells into initially 100 SOMs (clusters) based on “similarity” over the 27 antibody-derived metal signal parameters. We used the *arcsinh* c.f.1 and Z-score normalised versions as previously reasoned. Figure 4A shows the 100 SOMs for the output of the 109,535 single cell objects generated by the “Tonsil Ilastik –EPCAM membrane” segmentation approach in the form of a radial spanning tree with the mean expression of the fiducial markers CD79a and CD3 used as the radial statistic for each of the three plots respectively. In Figure 4B we present the same visualisations but this time using the 84,268 single cell objects derived from the “Nucleus only” segmentation approach. A qualitative comparison between the two sets of data suggest that the sub-optimal segmentation output (Nucleus only model) leads to more SOMs (clusters) that seem to have higher expression of CD3 and CD79a but also less radial spanning structure compared to the “Tonsil Ilastik –EPCAM membrane” model SOMS. To further investigate these potential differences we compressed the 100 SOMs to 30 consensus SOMs (cSOMs) using the standard hierarchal approach (39). We then sought to annotate the clusters based on the heat map outputs and marker expression pattern on a per-SOM, per-marker basis using heat maps. Figure 4C shows that the majority of “Tonsil Ilastik –EPCAM membrane” model consensus SOMs could be assigned a biological identity (27 out of 30) using expert a priori knowledge whereas for the 30 cSOMs derived from the “Nucleus only” segmentation model we were only able to confidently assign identities to 23 (figure 4D). Interestingly we also noted a reduction in T cell and macrophage cluster heterogeneity in the data derived from the “Nucleus only” segmentation model with no evidence of naïve CD8 T cells or mature macrophages as well as an over-clustering of B cells. Overall these data collectively suggested that sub-optimal segmentation did have a negative impact on phenotypic identification based on clustering approaches where all “*n* dimensions” are considered. While FLOWSOM has been widely used for suspension cytometry data analysis, we are not aware of a study using it for IMC data analysis. IMC data clustering tends to be done using the Phenograph algorithm (9,23,40,41). As such we wanted to also test this approach on our “Tonsil Ilastik –EPCAM membrane” data set. Again we used the optimal *arcsinh* c.f.1 transformed, Z-score normalised metal signal parameter feature set and selected a “k nearest neighbour” value of 17 to generate a similar number of Louvain communities (clusters) to our FLOWSOM consensus approach (30 clusters). Figure 4E shows the Phenograph output as a heat map with attempts to assign cell identities to the clusters. In this case we could only confidentially annotate 18 out of the 30 clusters and as a result several populations were totally absent compared to the equivalent FLOWSOM approach (Figure 4C). It was also of note that Phenograph found several non-classified (NC) clusters that were of very low frequency also suggesting a suboptimal performance compared to FLOWSOM as our panel was not designed to find any rare cell types in tonsil.

**Figure 4:**
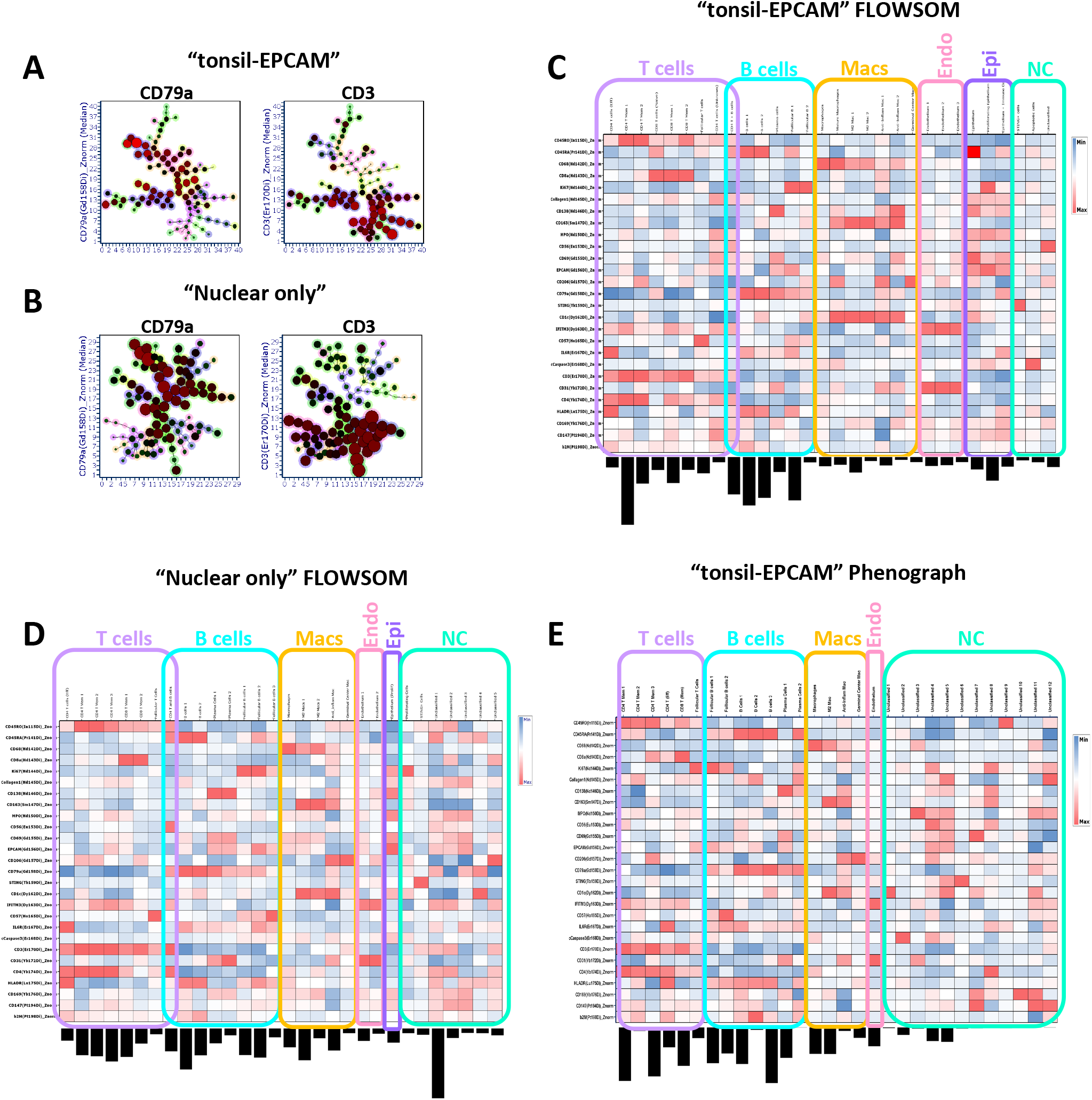
FLOWSOM clustering performs better on IMC data than Phenograph but is affected by suboptimal segmentation. (A) Radial spanning trees of the original 100 SOMs generated by the FLOWSOM clustering algorithm from single cell outputs generated by the “Tonsil EPCAM” segmentation model with the mean expression of the “fiducial” markers CD79a and CD3 used as the radial statistics as indicated. (B) The same plots as in A but for the output of FLOWSOM clustering on the single cell data from the “Nucleus only” segmentation model. (C) Heat map of the 30 consensus cluster (SOMs) derived from the original 100 SOMs for the “EPCAM” model. The frequency of each cluster is indicated by the bar chart below each column (cluster). Specific as well as broad cluster annotations are provided for T cells, B cells, Macrophages (Macs), Endothelial cells (Endo) and Epithelial cells (Epi). Where a cluster could not be confidently identified, they were labelled “NC” (not classified). The heat map is showing the median of the *arcsinh* c.f.1 transformed, Z-score normalised 27 antibody marker signals as indicated on the y axis (rows) and have been further normalised by column value (by cluster). (D) An analogous heat map as shown in C but for the 30 consensus cluster (SOMs) derived from the original 100 SOMs for the “nucleus only” segmentation model. (E) A heat map as shown in C but for the 30 Louvain communities (clusters) derived from analysing the single cell outputs generated by the “Tonsil EPCAM” segmentation model using the Phenograph clustering algorithm with a “K-nearest neighbour” value of 17 (see methods)

### FLOWSOM clustering combined with expert cluster merging is able to identify cell types/states with high spatial accuracy

Having established the optimal clustering approach for correctly transformed and batch normalised IMC data we wanted to further refine our clusters in terms of biological meaning. Several of the annotated cSOMs from the FLOWSOM approach were still phenotypically identical to one another and thus were unlikely to represent truly unique cell types or states. We also wanted to combine out final annotated cSOMS with the use of PacMap dimensionality reduction. To this end, we manually merged any of the 30 cSOMs from the heat map shown in Figure 4C based on highly similar marker expression patterns. This left us with 21 unique clusters, all of which could be biologically annotated with a high degree of confidence apart from one cluster of cells with high CD56 expression present as a majority in a single image (see Fig S2). Figure 5A shows the heat map of the 21 manually merged cSOMS with biological annotations and the relative frequencies of each. The clusters were also mapped back on to the same PacMap plot constructed from all 109,535 cells as shown in Figure 2B. The follicular B and T cells formed a distinct structure as did the non-follicular immune cells and the macrophages/ structural cells (endothelium and epithelium). The real power of IMC and other high parameter tissue imaging approaches is that spatial context of all cell phenotypes/clusters can be mapped back in to the tissue space. We chose human tonsil, the antibody panel and the specific ROIs precisely as they should identify well known cell types that also possess well defined spatial co-ordinates with respect to anatomical structures but also in relation to one another. To validate our final 21 manually merged cSOMs we used the fact that each of the 109,535 cell objects identified by Tonsil-EPCAM segmentation within the 24 ROIs retained their x and y centroid co-ordinates as part of the FCS file creation (see methods and supplemental methods). This meant we could simply plot the X and Y centroid features for any ROI as bi-variate dot plot and colour by the selected cSOMs. Figure 5B shows the spatial mapping of six annotated cSOMs from the heat map in 5A for two representative ROIs. Reassuringly the spatial locations of each cluster followed the expected biological patterns with the follicular T and B cells mapping to follicular structures and the endothelial cells mapping to the inner walls of the vessels/tonsillar crypt. These observations were further verified by the original staining patterns in IMC images (figure 5C) with cells in the follicles Ki67 positive as they are undergoing aggressive proliferation. As a final level of validation, we mapped all 21 clusters using unique colours on to the cell object maps derived from the segmentation pipeline (Figure 5D). These were also in agreement with the expected spatial patterns of locations. The coloured cluster maps for all 24 ROIs are shown in Figure S8. Overall the combination of manual merging of FLOWSOM derived cSOMs, PacMap visualisation and validation by spatial mapping confirmed our analysis approach to be optimal and accurate with respect to our panel and tonsil tissue.

**Figure 5:**
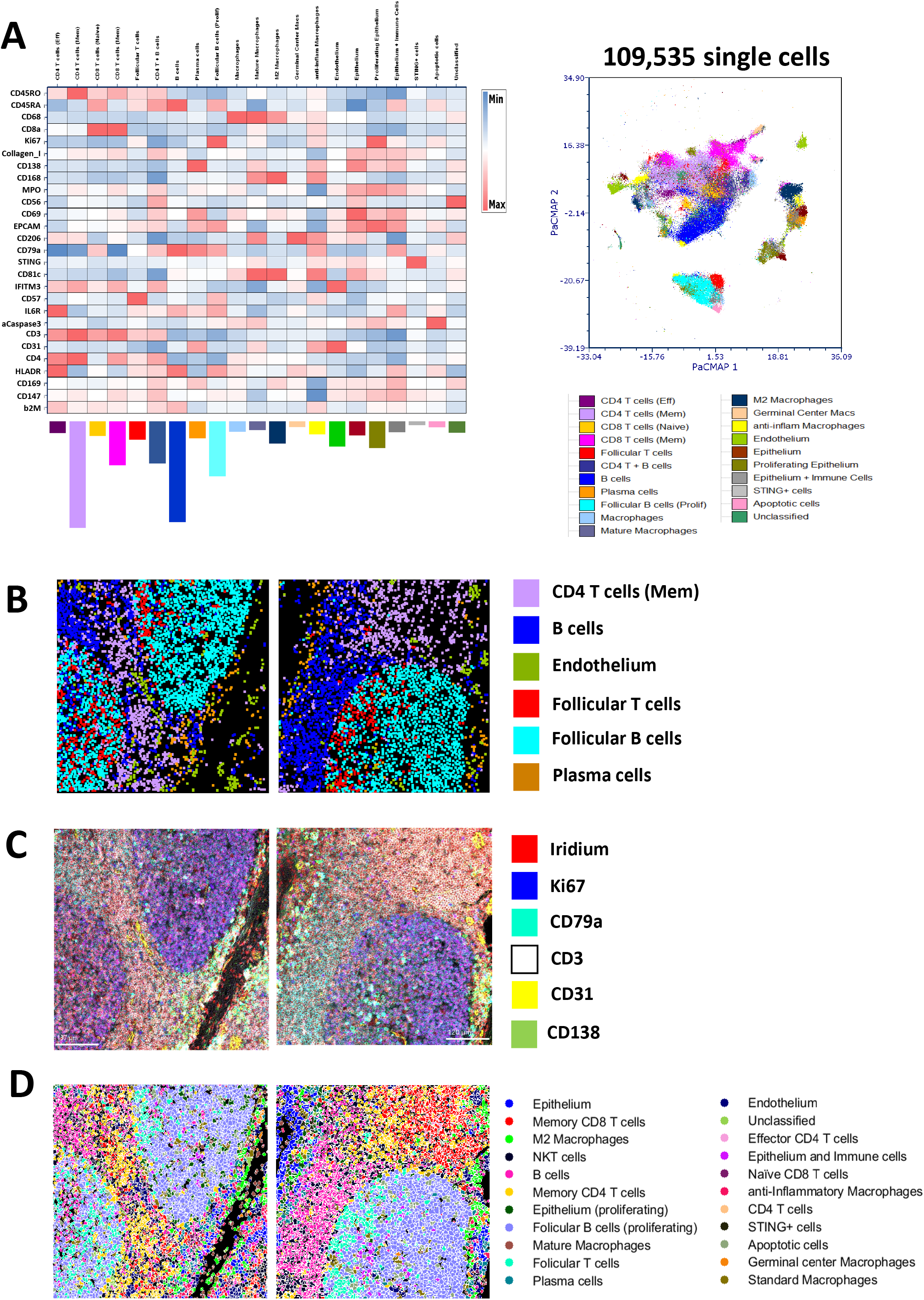
Cell type/state verification using the original IMC images and spatial mapping visualisation tools. (A) A heat map (left panel) of the 21 final manually merged populations created from the 30 consensus clusters for the “EPCAM” segmentation model shown in 3C. Cluster frequencies are shown by the coloured bar charts below each cluster column and the map intensity has been derived from the *arcsinh* c.f.1 transformed, Z score normalised markers as indicated in each row with further normalisation down each column (by cluster). A PacMap dimensionality reduction plot (right panel) of the 109,535 segmented single cells from all 24 tonsil ROIs across the 12 staining batches as shown in figure 2C but now coloured by the final 21 FLOWSOM clusters as per legend. (B) x/y centroid maps for two representative tonsil ROIs with 6 different unique cell (see legend) clusters displayed. (C) Pseudo-coloured IMC images of the same representative tonsil ROIs as in B showing 6 fiducial stains as indicated in the legend (nuclear plus 5 antibodies) that support classification of the cell types in A-B. (D) Cluster maps of the same two representative tonsil ROIs as shown in B-C with all 21 final consensus clusters shown (see legend).

### The choice of pixel expansion approach combined with removal of edge cells has a negative impact on neighbourhood mapping

Having established that our analysis approach could reliably identify cell types and states in tonsil tissue with high accuracy both phenotypically and spatially, we wanted to use these data to benchmark our neighbourhood analysis. Our method was based on the previously described approach from HistoCat (9) and is based on a defined pixel outgrowth that creates a bounding box in which significance of interaction or avoidance is tested using a permutations-based approach with a significance cut off (typically 100 permutations and a 10% cut off). There is also a threshold parameter that can filter out clusters that only appear in a certain percentage of the images/ROIs (see Figure S9 A). A threshold of 0.1% means that clusters have to be present in over 10% of all ROIs to be considered in the neighbourhood analysis. This was not a function relevant to our data set however as all final 21 cSOMs were present in all 24 ROIs (see Figure S9B). We would also caution against using this feature as it could lead to removal of a key, biologically defining cluster from one sample group in a large batch analysis. A further important consideration is the removal of any partial and fragmented cells around the edge of the image as well as size-based filter for removing under and over-segmented objects. We set a gate on our data that would ignore cells/objects less than 20 µm^2^ (5 µm^2^ diameter) and more than 200 µm^2^ (16 µm^2^ diameter) as shown in Figure S9C. We could then compare the outcomes and results of using this filtering method with edge cell removal versus analysing all objects. First we wanted to compare the results of conducting a spatial analysis on a defined, well characterised ROI (ROI 23 in this example) using the “bounding box” approach versus a “disc”-based method of pixel outgrowth (see Figure 6A). Our first metric of assessment was the median nearest neighbour number (Median NN) versus the pixel outgrowth value (see Figure 6B). As expected for all approaches, increasing the pixel distances led to an increase in the median NN with the largest values coming from the original bounding box approach. However based on the physical geometry of the cells in the tonsil tissue we reasoned that a median NN value between 6 and 8 would be indicative of an optimal area for “true” neighbourhood analysis. The usual recommended pixel outgrowth for this approach is 5 (9,23), however the data in Figure 6B showed that the original bounding box method gave a median NN of ∼11 cells at this distance. This suggests that the bounding box created was sampling an area greater than the area occupied by the immediate nearest neighbouring cells. Both the “disc”-based approaches at 5 pixels gave NN values of 8 regardless of any object filtration suggesting that it was more accurately finding “true” immediate neighbour cells compared to the original “bounding box” approach. Finally we wanted to generate heat maps for each of the 12 combinations and focus on the accuracy of the interactions and avoidances at the clustering level. We first considered the general “qualitative” appearance of the heat maps with regard to how much red (significant interaction), blue (Significant avoidance) and white (indifference) we observed. Figure 6C shows that in qualitative terms, only the disc method with object filtration (first column) produced heat maps over the entire pixel range tested (3 – 15) that were not dominated by red (significant interactions), rather the majority of cluster relationships were “indifferent” or random (white). Based on the “ground truth” image of ROI 23 in Figure 6A, the majority of clusters were not interacting with one another suggesting that the disc with object filtering method was more accurate in reflecting the actual spatial arrangement of cells in the tissue. Finally we focused on the four cell types (clusters) highlighted in Figure 6A; namely follicular B cells, memory CD4 T cells, memory CD8 T cells and B cells and looked at the heat maps for each up to a 10 pixel expansion. The only condition where we noted significant avoidance (blue) between follicular B cells as the central cluster (row) with respect to memory CD4/CD8 T cells and epithelium as well as an acceptable median NN value was the disc approach with filtering and a 5 pixel outgrowth. Overall these data support the idea that this was the optimal approach for finding “true” neighbours.

**Figure 6:**
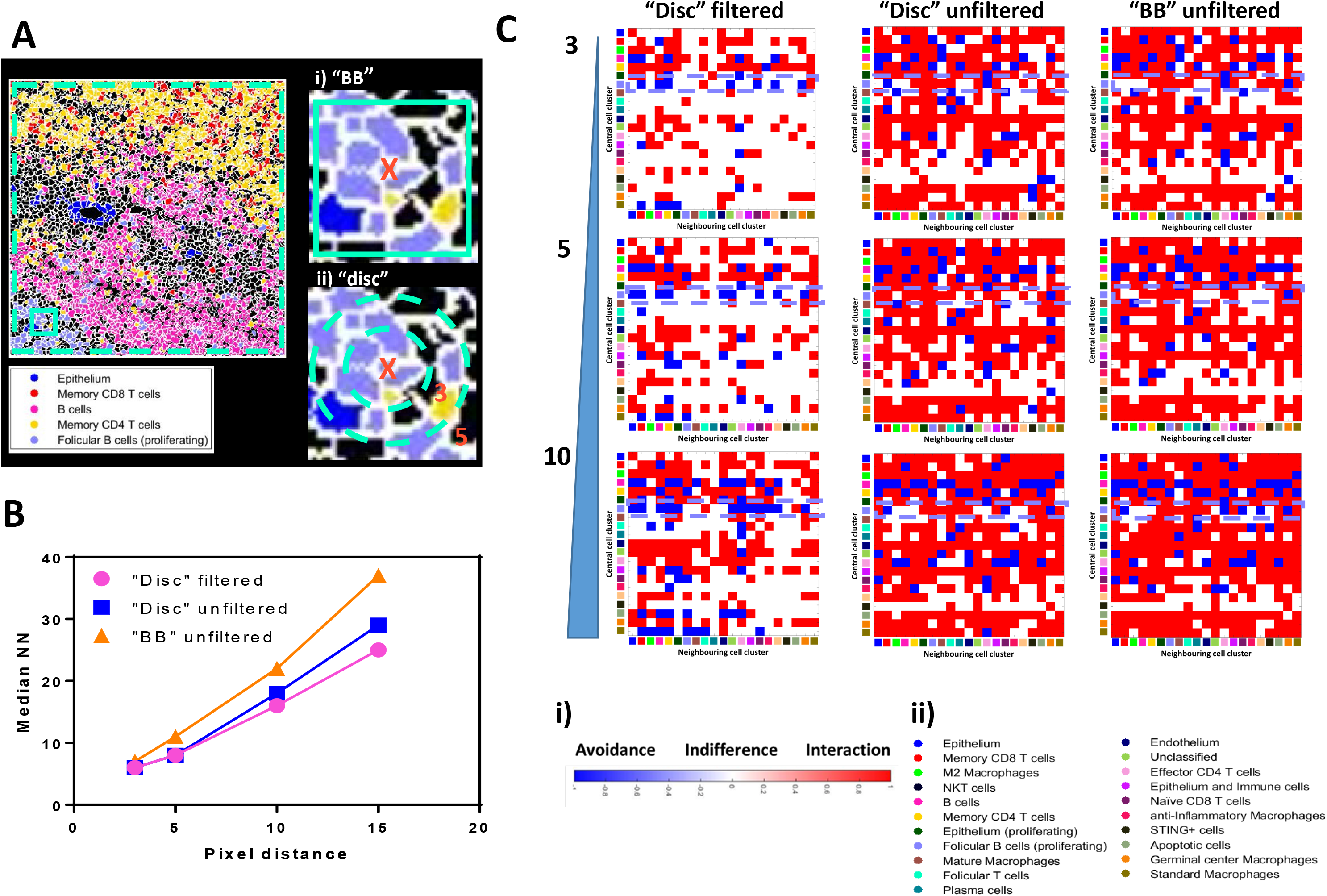
Optimisation of spatial “neighbourhood” analysis of human tonsil tissue reveals the importance of edge cell removal and the method of pixel expansion. (A) A cluster map for ROI 23 (see Figure S8) only showing 5 of the final consensus clusters (see legend) with clear and expected spatial relationships. The teal dotted line denotes the optional removal of edge cells prior to neighbourhood analysis. The area of tissue within the solid teal square has been magnified to show the two methods of pixel expansions for finding neighbouring cells to the central cell (X); a “bounding box” (BB) approach (i) or a disc approach (ii). (B) A graph showing the relationship between the selected pixel expansion/distance value (x-axis) and the median number of nearest neighbours (NN) for each of the three conditions tested (see legend). (C) Interaction heat maps for the different input options shown in B (columns) versus pixel expansion distance (rows, 3, 5 and 10 pixels). The rows for each heat map denote the central cell cluster (marked as X in A) and the columns denote the potential neighbouring cell types (clusters). The colour of each square in the grid relates to the nature of the spatial relationship with red denoting a significant interaction, blue a significant avoidance and white indifference as per legend (i). The clusters in x and y are denoted by colour as per legend (ii). The violet dashed boxes highlights the interactions and avoidances of the follicular B cell population with the memory CD4 and memory CD8 T cell populations. All maps were created using significance cut off of 10% and 100 permutations.

## Discussion

The analysis of IMC data has historically been challenging with limited attempts made to develop accurate, scalable, and accessible solutions. Moreover, approaches tend not to be very accessible and require expert knowledge of programming languages such as R, Python or MATLAB (12,42). This has resulted in significant frustration in the community, contributing to inaccurate single cell data and unconvincing biological conclusions. There are several issues with existing analysis pipelines, namely the approaches used to segment single cells accurately from low resolution, often “noisy” data. They often lack any real appraisal of how successful the segmentation is in terms of how well they find known/expected cell types/states in a given tissue. To this end, the “OPTIMAL” framework for analysing IMC-derived multiplexed image data provides several recommendations for testing, optimising and benchmarking key steps in any pipeline. Firstly we show using the currently most established approach for IMC data segmentation (10) that object identification is improved by the use of an Ilastik-generated probability map constructed using only nuclear and background signal but that the p-map does not necessarily need to be derived from the same cell or tissue type as an embedded cell line (Vero cells) performed comparably. Of note, we only utilised two classes of pixel classification for Ilastik learning (nucleus and background) and not membrane as this reduced the computational burden and showed no benefit over using three pixel-class Ilastick learning to define cell boundaries (Figure S10). When we also looked at the frequency of so called “nonsense” CD3/CD79a DP cells we found a minimal increase as a result of under segmentation in the absence of a p-map input. While this seemed surprising, it likely reflected the fact that T cells and B cells are often in quite distinct anatomical locations in human tonsil (43). Perhaps unsurprisingly however, when we took the data to the clustering stage, we did see dramatic effects on the fidelity and identity of the cell types and states derived from poorly segmented images. As such we would recommend always assessing any segmentation approaches using a clustering-based metric against a known “ground truth”. We saw no measurable benefit to using an approach for cell boundary detection that used the sum of multiple membrane signals over a single membrane marker such as EPCAM; even considering that EPCAM was labelling membranes in a non-specific fashion. Moreover IMC is not an optical imaging technology and has a relatively low resolution (1 µm per pixel, equivalent to around 10x magnification on an optical imaging system) with an ablation laser that “drills” in to a depth of tissue in order to liberate sufficient material to achieve suitable metal ion detection and signal resolution (1) it is therefore highly likely that information from cells from different z planes are mixed. As such, one could argue that segmentation will always be flawed to an extent. Certainly there are numerous approaches that do not attempt to segment cells but instead work with the pixel level data in the image (44). To this end, several groups are working on 3D imaging of cleared tissues (45) or by modifying the IMC approach to achieve Z stack information (46), however at present these techniques either lack the parameter space or throughput. We also tested the deep learning-based segmentation approach CellPose (17,24) as well as an approach that only segmented nuclei and saw no measurable benefits (Figure S10). None-the-less, we propose that OPTIMAL provides a framework for benchmarking any segmentation approach.

Post-segmentation but prior to any further single cell analysis using dimensionality reduction and clustering techniques it is essential to apply various transformations and corrections to the data in order to remove noise, background, maximise signal resolution and remove any batch effects. Any form of semi-quantitative tissue imaging is by nature composed of quite poorly resolved signals due to the fact that we are never measuring a whole cell, unlike flow cytometry or suspension mass cytometry. Moreover, IMC is around 5-fold less sensitive than fluorescence-based detection (our unpublished observation). As such it is imperative to ensure that the resolution of signal is optimised in order to provide the very best overall data structure prior to going in to both dimensionality reduction (DimRedux) and clustering. As described previously (22), we applied spillover correction to all of the mean pixel values for all metal ion channels. This has been shown to improve data interpretation. We also removed any “hot” pixels by capping at the top and bottom 5% for analogous reasons. However probably the most important step is the use of data transformations such as the hyperbolic arcsine (*arcsinh*). Without applying such a transformation, comparatively high parameter values with greater variance will have a lower weighting in any subsequent dimensionality reduction or clustering compared to lower values with greatly reduced variance. Using a very simple approach based on the Fisher discriminatory ratio (also known as the Linear Discriminate Analysis (LDA)) we used our OPTIMAL approach to determine the best *arcsinh* cofactor value for IMC data to be “1”, not between 5 and 15 as reported by others (30). We show that values greater than or less than 1 do not project the IMC data structure in an optimal fashion. While previous attempts have been made to try and develop frameworks for optimising parameter transformations for fluorescence-based flow cytometry data (47), to our knowledge OPTIMAL is the first for IMC data.

After parameter transformation, the next important step of data pre-processing is to look for, and if necessary, correct for batch effects. While we purposefully attempted to minimise and where possible eliminate all sources of batch effect by the same person staining TMAs from the same FFPE human tonsil section with the same panel of 27 metal tagged antibodies on 12 separate occasions and acquiring data on a well maintained and consistently QCe’d Hyperion IMC system, the nature of FFPE tissue analysis remains highly variable. As such it was no surprise that a UMAP-based analysis of our 27 *arcsinh* c.f.-transformed antibody-metal parameters revealed measurable batch effect in the data structure and presented us with the perfect opportunity to develop an OPTIMAL solution for correction. As with parameter transformation, there has also been a lack of exploration as to what the best approach is for batch effect normalisation, and while several approaches exist for cytometry and single cell data often these are not tested using actual dedicated, empirically generated batch controls. We did test a number of these approaches on our data set, including the “0-1” scale compression proposed by Ashhurst *et al.* (30) but found that a simple Z-score normalisation after *arcsinh* transformation was sufficient to remove all measurable batch effect from our data without eliminating biological relevance.

Having formulated the OPTIMAL approach for empirically determining the necessary transformations and corrections to achieve the very best resolution from our IMC data we next assessed the suitability of five different dimensionality reduction algorithms to determine which provided the best representation of our data structure. As DimRedux approaches are used to present multi-dimensional single cell data in a way that can aid interpretation it is essential that the right approach is used. The use of widely available software such as FCS Express of FlowJo that requires little to no knowledge of coding means that using our OPTIMAL approach, researchers can easily explore what DimRedux method is best for their data. To our knowledge, existing analysis methods such as HistoCat and ImaCyte do not offer the same flexibility of choice and ability to also alter the hyper parameters for these algorithms (iteration, seed, neighbours, perplexity etc.). In this case we found that PacMap performed the best with UMAP a close second. PacMap generated more discrete structures as well as showing improved mapping of fiducial markers back on to these whereas other approaches lacked any discernible structures and fiducial markers were more diffuse in mapping. Of note, PacMap also clearly identified the follicular structures in the tonsil driven by Ki67, CD57, CD3 and CD79a expression and has been proposed to handle weakly resolved signals better than other DimRedux approaches (36), making it highly suitable for IMC data.

The use of clustering approaches to identify cell types and states based on marker expression levels/patterns is well established in single cell analysis. There are several different approaches and one of the best performing is the FLOWSOM algorithm (7). To date, few if any IMC analysis approaches have reported the use of FLOWSOM for cell type and state identification, but rather have used Phenograph as part of HistoCat or IMaCYte (9,23). Our data was conclusive in that FLOWSOM, in conjugation with the Tonsil-EPCAM segmentation model, *arcsinh* c.f. 1 parameter transformation and subsequent Z score normalisation could identify the majority of expected cell types and states within the human tonsil. Phenograph performed poorly, missing several expected cell types as well as generating a large number of low frequency, unidentifiable clusters. While it may be possible to try and optimise the Phenograph algorithm to improve the outputs, in all cases we deliberately used the default hyper-parameters for both clustering algorithms we tested to mainly reflect that we want the OPTIMAL framework to be accessible to non-specialists in data analysis.

Finally, having arrived at a set of cell clusters that we could annotate with phenotypic, functional and spatial confidence, we wanted to assess whether we could benchmark and optimise the commonly used neighbourhood analysis method used in HistoCat (9). We purposely ensured that we selected quite varying areas within the tonsil tissue sections over the 12 batches, to introduce variance at the spatial level (but not at the cell type or marker level). We selected example ROIs where there was clear spatial definition of different cell types so that we could always compare any interaction-based neighbourhood analysis with what we could observe to be true in the image. We also restricted our analysis to only a few cell types such as follicular B cells and memory CD4 and CD8 T cells. We tested a number of different tune-able parameters for detecting immediate neighbouring cells from a central cell phenotype and used two metrics to determine which was best; the median number of nearest neighbour (NN) and the significance of either interaction or avoidance as a heatmap. The median NN values as a function of pixel outgrowth was very interesting as it showed the original script’s “bounding box” approach to be including cells that were not true neighbours. We found that a better approach was to use a radial “disc”-based pixel outgrowth and that filtering of edge cells as well as small and large cells from the images was essential to generate the expected interactions and avoidances. Moreover this was optimal at 5 pixels, as recommended by HistoCat and ImaCytE by default but only when using the “disc”-based outgrowth method. While this approach was simple and informative, we did notice that the structural heterogeneity we purposefully collected in our data set meant that if we created interaction heatmaps from the average of all 24 ROIs the data was almost un-interpretable (data not shown). As such analysis of structurally heterogeneous tissue may benefit from other spatial analysis methods that the one we tested.

In conclusion, we show using the OPTIMAL approach that methods for segmenting single cells in IMC data can be assessed using well characterised tissues and antibody panels followed by cluster analysis to verify that the expected cell types/states are identified. However prior to any clustering analysis, IMC data structure can be optimised by transforming all metal parameters with an arscinh c.f. of 1, and this can be empirically tested using the Rd approach, and also corrected for batch effect using a an additional Z score normalisation. We also found that PacMap was the best dimensionality reduction approach for visualising IMC data and that FLOWSOM was the best performing closeting algorithm. Finally, we show that the OPTIMAL approach for conducting neighbourhood analysis of the resident cell types/states is to use a “disc” based radial pixel outgrowth rather than a “bounding box approach”. We have further validated and utilised the OPTIMAL approach to analyse several other tissues using similar and distinct panels of antibodies to the ones used in this study. These include COVID-19 post mortem lung tissue (manuscript submitted), gut tissue from various inflammatory conditions (manuscript in preparation) and inflamed synovial tissue from rheumatoid arthritis patients (manuscript in preparation). Furthermore the OPTIMAL framework has been used to analyse data from other multiplexed tissue imaging technologies that are fluorescence-based such as the Miltenyi MACSima with a high degree of success. As stated previously, we do not describe OPTIMAL as a new pipeline per se for analysing IMC and other multiplexed imaging technology data sets, but we do argue that it offers a framework for assessing, optimising and benchmarking existing and future approaches.

## Acknowledgements

This work was funded by UK Research and Innovations / NIHR UK Coronavirus Immunology Consortium (UK-CIC; MR/V028448) and the European Union’s Horizon 2020 research and innovation programme under grant agreement No 860003, and the JGW Patterson Foundation. This work was also supported by the United Kingdom Research and Innovation (grant EP/S02431X/1), UKRI Centre for Doctoral Training in Biomedical AI at the University of Edinburgh, School of Informatics. For the purpose of open access, the author has applied a creative commons attribution (CC BY) licence to any author accepted manuscript version arising We would like to thank Jennifer Doyle and Saskia Bos for critical review of the methods and fellow members of the Newcastle University Flow Cytometry Core Facility and Bio-Imaging Unit for advice and support as well as colleagues from Novopath for creating the TMAs used in this study. . Finally we would like to acknowledge the support and advice from Andrea Valle at Denovo Software/Dotmatics for guidance with FCS Express software and pipelines.

