## Supplemental Material for "OPTIMAL: An OPTimised Imaging Mass cytometry AnaLysis framework for benchmarking segmentation and data exploration"

**Hunter et al, Supplemental Notes**

#### S2: Expanded Methods

##### S2.1 Tissue Section Preparation and Regions of Interest Selection

Formalin-fixed paraffin-embedded 2mm human tonsil tissue cores were obtained from the Novopath Tissue Biobank (Royal Victoria Infirmary, Newcastle upon Tyne) and embedded into a 3 core Tissue Microarray (TMA). TMA blocks were constructed manually using PFM Medical biopsy punches. Cores were selected using haematoxylin and eosin-stained slides to guide suitable areas in the donor blocks. Cores were places in a paraffin embedding mould, heated to 65°C and embedded in molten wax before cooling to set. 8µm serial sections were using HM 325 Rotary Microtome (Fisher Scientific, USA) and mounted onto SuperFrost Plus™ Adhesion slides (Epredia, CAT#10149870).

##### S2.2 Panel Design and Antibody Validation

A 27-plex antibody panel was designed to identify the immune, signalling and stromal components in the surrounding microenvironment of the human tonsil (See Figure S1 and Table S1). CD3 was purchased pre-conjugated to the assigned metal (Table S1). For all other targets of interest, carrier free antibody clones were sourced, and validated using standard single marker immunofluorescence on appropriate positive control tissues. Following verification of staining pattern and performance quality, approved antibodies were subject to lanthanide metal conjugation using Maxpar X8 metal conjugation kit following manufacturer's protocol (Standard Biotools, CAT#201300). Antibodies conjugated to platinum isotopes 194 Pt and 198 Pt were conjugated as described in Mei et al. 2015 (1). (AbC™ Total Antibody Compensation Beads (Thermo Fisher, USA, CAT#A10513) were stained with conjugated antibodies and analysed in suspension using the Helios™ mass cytometer (Standard Biotools, USA) and successful conjugation was determined via detection of appropriate signal in the correct channel. Positive control tissues were then stained with conjugated antibodies. Suitable antibody titres were evaluated using Imaging Mass Cytometry™ via the Hyperion™ Imaging System (Standard Biotools, USA).

**S2.3 IMC Staining**

Tonsil sections were baked for 1 hour at 60°C in a dry oven followed by dewaxing in two changes of fresh xylene (Sigma-Aldrich, USA,CAT# 534056-4L), 5 minutes each. Slides were rehydrated in decreasing grades of ethanol (100% 90% 70% 50%), for 5 mins each, and then washed twice for 5 mins in Milli-Q® Type 1 ultrapure water. Antigen retrieval was carried out using Tris-EDTA (pH9) buffer 0.5% Tween in the microwave at maximum power for 20mins. Slides were cooled to 70 °C, washed in two changes of Milli-Q® water 5 mins each and followed by two washes in PBS (Fisher Scientific, USA, CAT# 10728775). Tissue sections were blocked with 3% BSA (Sigma-Aldrich, CAT# A3059) in PBS for 45 mins at room temperature and then incubated overnight at 4°C with a cocktail of the 28 metal-conjugated antibodies (Table S1). Slides were washed with 0.2% Triton X-100 (Thermofisher, USA, CAT#85111) for 8 mins followed by two 8 min washes in PBS. Cell nuclei were stained with Cell-ID™ Intercalator-Ir (Standard Biotools, USA, CAT#201192A) at a 1 in 400 dilution, for 30 mins at room temperature. The slides were then washed for 5mins in Milli-Q® water and air dried prior to ablation.

**S2.4 Hyperion set up, quality control (QC) and data acquisition**

Prior to each sample acquisition, the Hyperion Tissue Imager was calibrated and rigorously quality controlled to achieve reproducible sensitivity based on the detection of 175Lutetium. Briefly, a stable plasma was allowed to develop prior to ablation of a single multi-element-coated "tuning slide" (Standard Biotools). During this ablation, performance was standardised to an acceptable range by optimising system parameters using the manufacture's "auto tune" application or by manual optimisation of XY settings whilst monitoring 175Lutetium dual counts. After system tuning, Tonsil sections were loaded onto the Hyperion system in order to create Epi-fluorescence panorama images of the entire tissue surface to guide region of interest selection (ROI). Two ROIs of approx. 500µm$^2$ encompassing lymphoid follicles and surrounding structural cells were selected for ablation per batch run. Small regions of tonsil tissue were first targeted to ensure complete ablation of tissue during the laser shot with ablation energies adjusted to achieve this where required. Finally, ablations were performed at 200Hz laser frequency to create a resultant MCD file containing all data from ROIs. Correction of signal 'spillover' between isotopes was performed as per the protocol described in here https://bodenmillergroup.github.io/IMCDataAnalysis/spillover-correction.html without deviation (2).

##### S2.5 Basic image QC and segmentation

###### S2.5.1 Ilastik Model Training

Ilastik (3) is a "random forest" machine learning tool used for classification of pixels or objects from images. It relies on providing training images, partially labelled by the user, in order for the machine learning model to determine what features within the image are being identified by the human user. For the two Ilastik models trialled in this manuscript, pixels were classified into either nuclear or background. This simplifies the model learning and application processes. The models were both trained on nuclear Iridium 193 datasets, however one was trained on images acquired of tonsil tissue data, while the other was trained on images of "Vero" cells. This comparison served to determine if Ilastik model training material matching the input material for analysis was important for correct macro functioning. Partial labelling was applied to the training images followed by training of the Ilastik models. Following training, the models were applied to the training images and checked visually for regions where the model did not perform as expected. In these regions, additional labelling is then placed, and the model is retrained. Once output probability map images match the expected results sufficiently, the model is saved and applied to any further datasets of unseen data. In the experimental dataset whereby no Ilastik model was applied to the nuclear channel Iridium 193, the raw unprocessed channel was used for the following CellProfiler steps instead.

##### S2.5.2 CellProfiler Cell Segmentation

CellProfiler (4,5) is an image processing and analysis pipeline tool. In this research, we have developed a CellProfiler pipeline to take input individual channels from the MATLAB processing pipeline, along with Ilastik processed nuclear channels, to output single-cell quantitative data. The CellProfiler pipeline requires the input data folders to be specified within the program, along with any folder naming conventions to be explicitly listed. The intensities of the membrane channel, the nuclear input channel, and the nuclear probability channel are first rescaled to allow for comparable intensity levels between images, regardless of any batch effect that might be present. The nuclear probability maps are then segmented into binary objects. This is achieved through a global "otsu thresholding", splitting the pixels into three classes (combining the middle intensity pixels with the background pixels), leaving just the nuclear pixels as white on a black background. A threshold smoothing scale of 1 and correction factor of 1 is also applied, with shape then used to distinguish between clumped objects. The pipeline was set to automatically calculate the size of the smoothing filter and the minimum distance between maxima for splitting of clumped objects. To facilitate the faster processing of the large datasets provided by multiplexed images, the pipeline was allowed to use lower resolution images to identify the local maxima. The membrane channel is then smoothed (keeping edges) to facilitate the following step of identifying the cell boundaries. A propagation method was used, using the identified nuclear objects as initial seeds, along with global otsu thresholding (classifying middle intensity pixels to foreground). A regularisation factor of 0.05 is applied, meaning the boundary is determined by the intensity gradient in the membrane channel. Holes are filled within each object. The cell boundaries are then expanded by 2 pixels. Visualisation images are generated and saved for checking segmentation accuracy in quick sense-checks. The cell objects are then used to measure the intensity of each of the channels in their area, outputting intensity information along with some morphological data regarding the cells, all on a single-cell basis in .npy file format.

##### S2.5.3 ImageJ Macro Channel Combination

A Fiji macro was created to potentially aid with cell boundary segmentation by creating a single combined channel to potentially better represent the whole cell membrane for use in CellProfiler. The raw Hyperion images were difficult to interpret visually as a whole due to the quantity of channels (28 metal signals/channels). Channels containing membrane or cytosol markers could be looked at individually to determine overall cell outline, however in isolation many channels had relatively low signal-to-noise giving a poorly defined, speckled appearance, while the specificity of other markers gave a clear indication of cell boundaries of some cell types but not others, meaning the single channel option was not optimal as a visual reference.

The images were processed with a Fiji macro which combined multiple non-nuclear channels into a single reference channel with greater signal-to-noise than its individual parts for a clearer illustration of cell morphology. Following visual interrogation, 14 non-nuclear channels with the best signal-to-noise ratio were selected. These were: channel 14 (CD45RA In115), channel 15 (CD68 Nd142), channel 17 (Ki67 Nd144), channel 19 (CD138 Nd146), channel 20 (CD163 Sm147), channel 23 (MPO Nd150), channel 30 (CD206 Gd157), channel 31 (CD79a Gd158), channel 36 (IFITM3 Dy163), channel 38 (CD57 Ho165), channel 43 (CD3 Er170), channel 45 (CD31 Yb172), channel 47 (CD4 Yb174) and channel 48 (HLADR Lu175). To process the images, the contrast of each channel was increased individually using "Enhance Contrast" with a default saturation of 0.35%. The look up table (LUT) was then applied. "Despeckle" was run on all of the channels to smooth noise. The channels were then stacked and projected into separate maximum intensity projections (MIP). The following outputs from each Hyperion image stack were produced: a composite image showing nuclear (green) and non-nuclear (red) regions for visual validation of segmentation, and a single channel non-nuclear MIP for use in CellProfiler for segmentation.

**S2.5.4 Spillover corrections, transformation, normalisation, metadata addition and .FCS file creation.**

Output npy files are subsequently processed within the MATLAB script. This portion of the script has three main functions: 1) Correct the data for spillover between the channels; 2) scale the data according to an *arcsinh* transformation; and 3) normalise the data to correct for batch variation (Z correction). Following these steps, the files are converted into FCS format to facilitate use with FCS Express for cluster analysis.

Due to the nature of multiplexed imaging, a degree of spillover between similarly labelled compounds is to be expected. As the Hyperion system, used for this analysis, relies on the distinction between very similar weight metal ions (even different isotopes of the same metal), the spillover could negatively impact interpretations, especially in a variation-rich field such as biology. To remove the contribution of metal isotopes in non-target channels we followed the protocol as outlined by Chevalier et al (2) whereby a slide was made that had each of the 28 metals used in this study spotted individually. The Hyperion system was then set to ablate each spot one at a time and measure across the entire detector range the proportion (%) of the target channel that appeared in non-target channels in order to create a spillover correction matrix that was used to subsequently correct the mean intensity values for each channel derived from our multi-stained tonsil tissue.

A Hyperbolic arcsine (*arcsinh*) scaling factor (6) is applied to the data with the intention of ensuring the high variance of a potentially larger value (positive signal) is given the same significance and weighting as a negative (low) signal that has low variance. This scale normalisation is essential for dimensionality reduction and clustering algorithms to work well.

Z-score normalisation was then applied to the spillover-corrected and *arcsinh*-processed data. The goal of this step is to align the median around 0 and compresses the scale to minimise influence of comparative brightness from batch effects and also inter/intra-marker variance due to biology and batch effects respectively.

##### S2.6 Data analysis and exploration using FSC Express software

###### S2.6.1 Introduction

The “OPTIMAL” analysis approach is designed to work with any software, script or package that can read FCS file format data. We have decided to use FCS Express from DeNovo by Dotmatics as it is a widely available software solution for cytometry data analysis, however the “OPTIMAL” approach could be executed using other commercial software as well as open source solutions. Successful implementation of this approach using FCS Express requires some basic and advanced knowledge of the software. We would direct anyone to look at the extensive YouTube tutorials that can be found here https://denovosoftware.com/videos/. However as much as is practical, we will outline the key steps in this supplemental methods section. Note that this is not designed to serve as a user guide for the software, which already exists https://denovosoftware.com/full-access/manual-tutorials/. The result of our segmentation approach is the generation of .FCS files that are composed of a combination of morphometric features (area, eccentricity), spatial features (x and y centroids) sample metadata (batch number, patient number) and metal “intensity” features (mean pixel intensity) that have been “compensated” (corrected) for signal spillover as described previously. Each metal intensity parameter exist in two formats, one set of metal intensity features have been *arcsinh* transformed using the pre-determined optimal cofactor for IMC data of 1, and the second set have been Z-scored transformed in addition to the *arcsinh* transformation in order to minimise the influence of technical variation (batch effect) and also natural variation in the intensity of various markers/antibodies that would dominate the data structure (DimRedux visualisation, clustering or heat maps).

###### S2.6.2 Determining the optimal transformation cofactor for IMC data

In order to determine the optimal *arcsinh* transformation co-factor value for IMC data we performed a “titration” of possible values from 120 to 0.1 using the original (pixel capped and compensated) metal signal parameters and using FCS Express "mathematical transformations” option in FCS Express “Pipelines”. We then used the Fisher Discrimination Ratio (Rd for short), also known as “Linear

Discriminate Analysis" (LDA) to enumerate the separation and resolution between a gated low (or negative) and high (or positive) signal using simple gating in FSC Express. The Rd is calculated as follows (7):

$$Rd = \frac{(Median\ hi - Median\ lo)}{(rSD\ hi + rSD\ low)}$$

It is then possible to plot for any parameter the *arcsinh* transformation value (x axis) versus the Rd value (y axis) to determine the cofactor that provides the best resolution of low and high signals and thus maximise the data resolution and structure prior to any dimensionality and clustering. However we strongly suggest that the use of these transformations be empirically tested using the OPTIMAL approach.

**S2.6.3 Dimensionality reduction, batch effect determination and correction**

The fastest and simplest way to visualise multi-dimensional single cell data is usually via some kind of dimensionality reduction approach whereby *n* dimensions are compressed and represented by 2 or sometimes 3 parameters (8). There are numerous options to achieve this, all with various benefits and drawbacks as well as tuneable hyper-parameters that will influence the output. In all cases the choice and implementation of the dimensionality reduction method must remain true to the original underlying data structure. Via the use of FCS Express and the "pipelines" tool we were able to assess multiple dimensionality reduction approaches using both embedded options within the software (tSNE and UMAP) as well as a number of external algorithms (TriMAP, PacMAP and fltSNE) via the python script interface in the FCS Express pipelines module. References for each of these algorithms can be found in the main manuscript. Information and instructions on how to run python scripts as part of the FCS Express pipelines module as well as the necessary software links and scripts that need to be installed with the relevant publications can be found here https://denovosoftware.com/full-access/knowledge-base/python-transformation/.

To begin the analysis open a new incidence of FCS Express and navigate to the "data" menu on the top options ribbon and select the "data list" option. At this point it is possible to select the "+" icon and choose which files to load. For this stage it is important to select the "merge FCS files" option as this will bring all samples in as a single merged entity and allow any analysis to be carried out simultaneously while still preserving individual file identity (a new column will be created called "file identifier" that can be used to de-convolute to individual sample level as necessary). At this stage, it is good practice to open up a plot by dragging the merged file on to the workspace. We tend to select a density plot of Area on the X axis versus eccentricity on the Y axis. By then "right clicking" on this plot and selecting "gate stats", this will show the total number of events loaded from all the merged files and can be cross checked against the expected total number of events. If these numbers are in agreement, one can proceed to set up and run a set of dimensionality reduction and visualisation methods to rapidly assess a range of key issues such as the presence of any batch effects, attempts to correct these as well as selecting the best overall method for presenting the data structure. Navigate to the "tools" option on the top options ribbon and select "transformations". This will open up a new window and clicking on the "+" icon allows you to then select the "pipelines" option in the drop down menu. Details on how to set up and run pipelines in FCS Express can be found here https://denovosoftware.com/flow-cytometry-pipelines/. Briefly selecting the pipeline option will open up a further window where it should be possible to input which parameters you want to have available for the further analysis. We recommend selecting all available parameters in the data set at this stage. There is also an option to "automatically run pipeline" that is ticked by default. This should be unticked so as to avoid the software attempting to run a pipeline before it has been finalised. Creating a new pipeline will activate a second "+" icon to the right of the first one. Selecting this will provide a new drop down menu where the required pre-set analysis options will be present as well as the Python terminal for running scripts that are external to the software (see later section). For the analysis of possible batch effect and to determine whether our chosen method of Z-score normalisation was able to remove it, we used UMAP. The Z-score normalisation equation is shown below:

$$z\ score = \frac{x - \mu}{\sigma}$$

This was done by selecting “dimensionality reduction” then “UMAP” from the drop down menu. This creates a new step in the pipeline that can be named (for example) “UMAP_*arcsinh*_only” and select in the option window to only use the *arcsinh* c.f. 1 transformed parameters that we know have successfully stained our tonsil tissue from our basic image QC step (2.3 and table S1). We used the default hyper-parameter settings for UMAP (number of neighbours = 15, min. Low Dim Distance = 0.1 and iterations = 500) we did however add a parameter suffix of “archsinh_only” to be able to identify each set of UMAP parameters in the same pipeline. We created a duplicate of this UMAP step in the same pipeline and renamed it “UMAP_*arcsinh*_Zscore” and selected the *arcsinh* c.f.1 and Z-score corrected versions of the same parameters as before. We kept the hyper-parameters of the UMAP identical but changed the parameter suffix to “*arcsinh*_Zscore” to provide distinction between these UMAP parameters and the other set of UMAP parameters. Once all steps were cross checked, the pipeline was executed by re-ticking the “automatically run pipeline” option and clicking on the icon to “apply to all plots” located to the right of the “+” icon. Successful launch of the pipeline can be determined by the progress bar at the bottom of the FCS Express workspace. Run time is dependent on numerous factors such as computer specifications, number of events/samples as well as the complexity of the pipeline, both in terms of steps and the algorithm itself. To give some context, generating two sets of UMAP parameters from the ~100,000 single cells across 24 samples took approximately 10 minutes on a Microsoft Surface Pro Series 4 with an 4 Intel(R) Core(TM) i7-7660U CPU @ 2.50GHz 2.50 GHz, 8 GB RAM and 64-bit operating system with x64-based processor. We were then able to create a colour dot plot for the *arcsinh* only UMAP parameters 1 and 2, as well as one for the *arcsinh* and Z-score normalised set. It was then possible to plot batch "batch number" versus Iridium signal (Z-score version) as a density plot and set individual gates for each of the 12 data batches with suitable colours. Displaying the batch gates on each UMAP plot allowed us to quickly assess if there was any batch effect (*arcsinh* only UMAP parameters) and if it had been corrected/minimised (*arcsinh* plus Z-score normalised UMAP parameters and *arcsinh* plus 0-1 normalised parameters).

**S2.6.4 PacMap visualisation via Python script interface**

Having shown empirically that Z-score normalisation of the *arcsinh* transformed parameters eliminated batch effects in the data, we used these for subsequent clustering and optimal visualisations. UMAP visualisation provided a good two-dimensional representation of the 27 antibody parameter data set, however we wanted to assess if there was an option that would improve upon this further. Our criteria for assessing these methods was based on the overall data structure projection; namely the presence or absence of defined "islands" as well as how well key fiducial markers mapped to these structures. For example, structures with concentrated maximal expression of CD3 or CD79a were preferred over apparent diffuse staining patterns. At time of writing, FCS Express currently has tSNE and UMAP directly available in the pipelines options. However it is possible to access and implement additional dimensionality reduction algorithms via the "miscellaneous" option in the "pipelines" sub menu and then selecting "Python Transformation". More details on what supporting software is required as well as the necessary scripts can be found here: https://denovosoftware.com/full-access/knowledge-base/python-transformation/. Briefly, once C++, Anaconda (Python) and the FCS Express Environment (for Anaconda) have been installed it is possible to select the desired script and install the package using Anaconda by selecting the FCS Express environment and opening up a new terminal. To install (for example) PacMap type **pip install pacmap** and the required dependencies will be installed automatically. It is then possible to copy the required script text from the FCS Express web link and paste into the python script window in the pipeline (clear the existing script first). If successful, on saving the relevant hyper-parameters should be available as well as a list of input features to select. As mentioned the *arcsinh* plus Z-score normalised parameters were used. For PacMap, the “n neighbours” value was calculated based on the recommended equation (9) and for 109,535 single cells this was “26”. For all other hyper-parameters and dimensionality reduction algorithms default settings were employed (Flt-SNE, TriMap, tSNE, UMAP).

**S2.6.5 FLOWSOM and Phenograph Clustering**

In order to determine the resident phenotypes in our segmented tonsil data set of 109,535 single cells we compared two different clustering algorithms namely FLOWSOM (10) and Phenograph (11); both of which are available in FCS Express directly in the “pipelines” menu. The panel of metal-tagged antibodies selected was designed to find at least 15 known cell phenotypes within the human tonsil (see table S1 and Figure S1) and as such allowed us to benchmark the different clustering approaches as well as the segmentation method. To create a FLOWSOM pipeline with embedded PacMap visualisation we created a new “pipeline” and then selected the “Pre-Defined Algorithms” option in the drop down. This allows access to a complete “FLOWSOM” script with all the required modules. The first step “new scaling” is not required as the data has already been transformed within the FCS files and can be deleted or unticked. The last step “New parameter removal” can also be removed. In the “New Batch Self-Organizing Map” module, we selected the *arcsinh* transformed Z-score normalised parameters representing the 27 phenotypic markers in the panel. All other parameters were left as default and as such a 10 x 10 SOM grid generated 100 clusters (SOMs). The only other module we altered was the “new consensus clustering” where we asked for 30 consensus clusters to be derived from the original 100 SOMs (twice the expected 15 clusters, so still over clustered). We then created a new pipeline module to run Phenograph clustering. Again this was done by selecting the Pre-Defined Algorithms” option in the drop down menu, but this time selecting “Phenograph”. As before, the first part of the module “scaling” can be removed as the data has already been suitably scaled/normalised. Within the “New K-nearest neighbour” module we selected a value for “number of neighbours” that generated 30 clusters. The value used for the 109,535 single cells was 17-18. Cluster number was assessed by creating a plate heat map from the “insert” tab “heat map” option and selecting “Louvain communities” to display on the x-axis. Once all required cluster parameters and PacMap coordinates were created and basic outputs checked, we exported the data as a single merged .DNS file (Data Stream) using the “export” function in the “data” menu. We also created individual .DNS files by repeating the export step but selecting the “export split on classification” drop down menu and selecting “classification identifier”. The reason to create .DNS files is that they are smaller than the source files and will contain all the previous features as well as the new cluster info and PacMap coordinates. All further analysis and exploration were done using the .DNS files.

**S2.6.6 Data visualisation, exploration and analysis**

A new incidence of FCS Express was launched and the merged .DNS file containing all original metadata and features plus the newly created dimensionality reduction coordinates for UMAP and PacMap as well as all FLOWSOM cluster information (100 SOMs and 30 cSOMs) was loaded into the data view. Any meta-data gates such as those set previously for batches were recreated using a bivariate density plot of batch (x) versus Ir 193 signal (*arcsinh* and Z-score version on y). The “plate heat map” option was selected from the “heat map” menu in the “insert” tab and either the “batch SOM cluster assignment” option was selected for x-axis display on the plot. This created a radial spanning tree of the original 100 SOMs. It was then possible to right click on this plot and select the “format” option then “overlays”. This gives access to the ability to select a different radial statistic for a given marker and emphasise any SOMs characterised by high expression of that marker on the plot. We then created a second plate heat map and this time selected “consensus clustering assignments” as the x-axis parameter from the plots drop down menu. This displayed the 30 cSOMs in a grid of 10 x 3. We then used the “well gate” function from the “gating” tab and selected all “wells” and asked to create individual gates for each with the prefix “cSOM” and to select individual colours for each (however this function often works poorly so manual colour selection is required). It was then possible to create a new PacMap colour dot plot, and select the "gates to display" option in the plot format menu (again accessible via a right mouse click on the plot).

In order to explore the outputs from the clustering (FLOWSOM or Phenograph), "parameter heat maps" were selected from the "insert" then "heat maps" sub menu. It is then possible to introduce multiple overlays of the same merged .DNS file on a "per gate" or in this case "per cluster" basis. To do this, right click on the newly created heat map and select "Add Overlay using Advanced Open Data Dialog". This is the quickest way to select the current open file for duplication. Once selected a new window will open and select the "duplicate for gates" option. Select all the necessary cSOM gates and click ok. The heat map requires further editing however to generate an output that can be displayed and interpreted. To modify the heat map further, right click and select "format". Select "specific options" and then choose (as a preference) to display in portrait mode. This now places the markers (parameters) on the y-axis (rows) and the cSOMs (clusters) on the x-axis (columns). At this stage, also select the option to "apply colours based on the values in each column" as this will normalise the heat map values by marker within each cluster (only possible because we have used the Z-score normalised versions). Next select "parameters to display" and ensure that the appropriate *arcsinh* and Z-score normalised metal channel features are selected by ticking the "only the items checked below" box and then selecting the correct parameters. Next go to the "overlays" option and select all overlays in the list. Change the "statistic to show" setting to the "median". It is also good practice to check that all the clusters are shown. It is also possible in this menu to move clusters (columns) left or right in the heat map order. This is useful after annotation to group all T-cell subsets (for example) together. It is also often useful to modify the axis. This can be done in the "axes" sub menu and allows one to change font sizes, colours and also rotate labels 90 degrees; this is useful for the cluster names. Cluster annotation was done using expert analysis and input. In this case it was guided by the panel information shown in table S1. Briefly, gate (cluster) names were edited to reflect broad or specific annotations. Where two clusters were clearly very similar, they were given a label such as "Memory T cells 1, 2 etc." Where the cluster was deemed to be unique it was given a more definitive annotation. To create merged populations, we used the "well gate" approach on a plate heat map showing all consensus clusters (so 30 in this case from both FLOWSOM and Phenograph). Merging was achieved by multi-selecting the appropriate clusters and the new merged cluster was given a definitive annotation (for example Memory CD4 T cells). A new parameter heat map was then created to only show the final merged consensus clusters.

As a second stage of validation, we also plotted simple colour dot plots with the centroid x (x-axis) versus the centroid y (y-axis) coordinates and coloured by either cSOMs or the final merged clusters. We did this to ensure that the cell types/states we had annotated mapped to the expected structure and anatomical locations in the tonsil (see figure S1). In order to be able to show one ROI/FCS file at a time, we also used the plate heat map function to display file identifier on the x-axis and create well gates for each of the 24 merged tonsil FCS files. We could then show/display each gated file/sample/image on the x/y centroid plot. We found that the centroid plots were visually more pleasing when we set the fill background to black and also removed all axes ticks, labels and titles. This can be done via the "format" menu.

Once all clusters were annotated we created a simple excel file that contained the name and number of the final clusters (so 1-22 for example) and info regarding which cSOMs made up these final clusters. So for example cluster 1 could be named as "memory CD4 T cells" and could be made up from merging cSOM3 and cSOM9 from the original 30 cSOMs. We also created a .CSV using the "export" option from the "data" menu. We only selected the minimal information needed for the advanced spatial mapping and neighbourhood analysis. These were "file identifier" "centroid x", centroid y" and "consensus clustering assignment". This created a matrix of all 109,535 cells across all 24 images, each assigned a cluster membership (1-30).

Finally, we also created a batch statistics export for further analysis of the final clusters. To do this, we opened up a new incidence of FCS Express and loaded the individual .DNS files for all 24 tonsil ROIs. As mentioned previously, these contained all the necessary consensus clustering info as well as the original meta-data from the original FCS files. We had to recreate the cluster (cSOM) gates using the plate heat map option and individual well gates with the "cSOM" prefix. We also created a single bi-variate plot and displayed the first file in the data list. Use any two parameters, however we often use centroid x versus centroid y. This is important as the "data source" guide for running the stats report as a batch process (see later). Once these gates were created (could be using any of the .DNS files), we navigated to the "batch" option on the menu ribbon. We then selected "batch actions". In this new window, under the "add report" window, we select the "export to EXCEL (column mode)". Leave the option to "start with an empty file" in the "append option" area and in the "output file options" direct where to save the file and what to call it. Select "ok" and it should now be possible to add in the statistics required in the output. To add these, right click on the "EXCEL (column mode)" in the "Batch Process Action" window and select "add item" and "statistics token". Select in the sub window "data source" and select the single bi-variate plot created previously. Next move to the "Statistic" menu. In the "gate" window select "no gate" and in the "statistic" window select "filename". This means the first column on the export will be the FCS file/ROI name. Press "Ok" to add to the export actions and it should now appear under the "EXCEL (column mode)" in the "Batch Process Action" window. Highlight the bi-variate plot so the borders turn red and drag it on to the "EXCEL (column mode) icon. This should open a new window that says "choose the format to paste". Select "statistics token". This will open up a new window where it is possible to select all possible gates (ungated to cSOM X) by holding down the shift key and left clicking on the last gate at the bottom of the list. In the "statistics" window, again hold down control and multi-select "filename", "# of events" and (optional) "% of all cells". Once the EXCEL stats export routine has been finalised, ensure that the bivariate plot is showing the data from the first file in the data list then select "Run" in the "Batch" menu. FCS Express will then visibly iterate through each sample and create an EXCEL file from the results. These data can then be further analysed either in EXCEL or PRISM (Graph Pad) and should be arranged as: First column: filename (ROI); Second column: Total #cells (ungated); Third column: #cells in cSOM 1 and so on. The export of frequencies is optional as they can be derived using the total cell counts and the counts per cSOM.

**S2.7 Spatial analysis**

The MATLAB code developed for spatial analysis works by examining each cell identified within the cell mask file for a single image and is based on the code previously described in HistoCat (12) . It uses that cell mask and expands the cell region by the number of pixels specified by the user within the code. The function will then count the number of unique cells found within the expanded cell area, excluding the original cell, keeping a list of each cell identifier within the label image. The cluster identities of each of these neighbour cells is then assessed. This process is repeated for each cell within a ROI. In order to compare the spatial arrangement of the cells for higher-than-expected or lower-than-expected interactions (where no positive or negative interactions are the nul hypothesis), the cell mask image is then used as a framework with the cell cluster identities randomly mapped onto the cell locations. The number of random images generated is dictated by the "Iteration" value specified by the user within the MATLAB code. If the original image shows a higher level of clustering than the randomly distributed images in more than 1-(significance threshold) proportion of the random images, then the interaction is deemed significantly positive for that image. If the reverse is true, showing a lower level of neighbour correspondence than the random images, then a significantly negative interaction is identified. In all other cases, no significant interaction is identified. The number of ROIs belonging to a group (i.e. pathology) that show a positive, negative, or neutral interaction is then averaged over the group condition, outputting a single heatmap with values ranging from a possible 1 (all images showed significant interaction between the cell types) to -1 (all images showed a significant avoidance between the cell types), with intermediate values indicating that some images showed interactions while others have not. The heatmap generated is asymmetric, allowing assessment of interactions that occur uni-directionally (i.e. cell type X interacts positively with cell type Y, but cell type Y does not interact in either direction with cell type X).

Tested within this research were three methods of spatial analysis, based on the above description. The first method (Ungated Original) used the pixel expansion method used within the HistoCat code. Another method (Ungated Disk) performs the process above the same, with the exception of using an alternative radial approach to cell area expansion (Disk erosion/dilation), which appears to better preserve the shape of the cells examined and thus more likely to find "true" neighbours. Finally, an additional gating step was incorporated (Gate Disk), which excluded cells based on two factors: area and edge. Only cells with physiologically reasonable areas (20-200 $\mu m^2$, equating to 5 – 15 µm dimeter) were included, and cells in contact with the edge of the image (and thus showing inaccurate neighbour reporting on their blank edged) were also excluded.

Visualisation of the cluster spatial information was also performed by applying a high-contrast colour labelling to the original cell masks based on their reported cluster identity, with the possibility to map only selected cell types for ease of visualisation.

**S2.8 Code availability**

All code and data files used in this manuscript is available here:

https://www.ebi.ac.uk/biostudies/studies/S-BSST1047

**Supplemental Figure Legends**

Table S1: Key information concerning the panel of 27 antibodies used in this study to stain human tonsil tissue over 12 individual, temporally distinct batches. The final two columns denote the "ground truth" cell type that each marker in isolation or in combination was selected to identify in human tonsil as well as the expected spatial location in order to benchmark the clustering approaches used in this study.

Figure S1: (A) A single IMC image showing Iridium (DNA, red pseudo colour) overlaid with Ki67 (white pseudo colour) and CD31 (green pseudo colour) as per legend key. The major anatomical and structural features are labelled. (B) A gallery of grey scaled IMC images showing staining patterns for each of the 27 antibodies in table S1 plus the two Iridium channels (191 and 193). (C) Images showing the comparative staining coverage of EPCAM (left panel) and a combination of immune cell membrane signals (right panel).

Figure S2: IMC images from all 24 Tonsil ROIs across the 14 temporally distinct staining batches. Images shown are at 1 pixel per micron and equivalent to a 10x optical image and are between 0.25 and 0.5 $mm^2$ of total image area. The pseudo-coloured overlays show DNA by virtue of Iridium intercalator (red), CD3 expression (Blue) and CD79a expression (Green) as per the legend. In all cases ROIs were selected to try and capture as much of the diverse tonsil structure as possible such as follicles/germinal centres and epithelium.

Figure S3: Cell segmentation maps for all 24 Tonsil ROIs across the 12 temporally distinct staining batches derived from the "Tonsil EPCAM model" of cell segmentation whereby the tonsil nuclear image was used to train an Ilastik model based on classification of nuclear pixels and background, then single cell boundaries were defined by the EPCAM membrane signal using CellProfiler to generate single cell information from the Ilastik model-derived probability map. The markers are denoted by the pseudo colours as shown in the legend.

Figure S4: Cell segmentation maps for all 24 Tonsil ROIs across the 12 temporally distinct staining batches derived from the “Nucleus only” of cell segmentation whereby the tonsil nuclear image was used directly by CellProfiler to generate single cell information without any Ilastik-derived probability map. In all cases the object boundaries are shown on a merge of the nuclear (iridium) and EPCAM images.

Figure S5: Bi-variate single cell level scatter plots for all 24 Tonsil ROIs across the 12 temporally distinct staining batches derived from the “Tonsil EPCAM model” of cell segmentation showing CD3 intensity levels on the x axis versus CD79a intensity levels on the y axis. Both parameters are the *arcsinh* c.f.1 transformed, Z-score normalised versions. Gates have been set to determine the frequency of CD3 and CD79a single positive events, as well the double positive (DP) “nonsense” cells.

Figure S6: Bi-variate single cell level scatter plots for all 24 Tonsil ROIs across the 12 temporally distinct staining batches derived from the “Nucleus only” of cell segmentation model showing CD3 intensity levels on the x axis versus CD79a intensity levels on the y axis. Both parameters are the *arcsinh* c.f.1 transformed, Z-score normalised versions. Gates have been set to determine the frequency of CD3 and CD79a single positive events, as well the double positive (DP) “nonsense” cells.

Figure S7: The top row shows UMAP plots coloured by batch (as indicated in the legend) created using the *arcsinh* transformed parameters only (first column), the *arcsinh* parameters transformed by 0-1 scaling (middle column) and the *arcsinh* parameters Z-score transformed. The middle row of UMAP plots are density weighted and the bottom row are density weighted by CD3 expression.

Figure S8: Cell cluster maps showing for all 24 Tonsil ROIs across the 12 temporally distinct staining batches derived from the “Tonsil EPCAM model” of cell segmentation as in S2. All 21 final cell type clusters are shown as per the legend.

Figure S9: (A) A graphical representation of how the cluster threshold function works when considering across all images what clusters to measure in the neighbourhood analysis. In the ten mock images shown, the brown population only appears in one image so setting a 10% threshold means that it will not be considered however at a 1% it would be. The purple population appears in nine of ten images so at a 90% cut off, this population and the brown population will not be included in any neighbourhood analysis. (B) A graph showing the frequency of all 21 final cSOMs in all of the 24 tonsil images. Note that all 24 images contained all of the 21 final cSOMs making the threshold function unnecessary in this study.

Figure S10: (A) Segmentation checker imagines for an expanded set of cell segmentation approaches as indicated. (B) Bi-variate plots for the single cell segmented outputs from the ROIs shown in A with CD3 (x axis) and CD79a (y axis) expressions levels plotted. Gates are set to capture the frequency (%) of CD3 single positive, CD79a single positive and double positive events. (C) PacMap plots for each of the single cell outputs for all ROIs from each of the respective segmentation approaches with expression density set on CD3 levels. (D) The same as in C but with expression density set on CD79a levels. (E) Heatmaps of the FLOWSOM clustering outputs from either the "nuclear only II" (left panel) or CellPose (right panel) segmentation approaches.

**This user guide will assist users in using the macro and scripts present within this repository. For further information or queries, please contact George Merces at**

#### Folder Organisation

1. Arrange your folders according to the naming convention listed below

| Name | Date modified | Type | Size |
|---|---|---|---|
| 1_ImageData | 22/05/2023 21:51 | File folder | |
| 2_image_extraction | 11/07/2023 15:10 | File folder | |
| 3_ilastik_model | 22/05/2023 16:27 | File folder | |
| 4_structured_data_for_CP | 24/05/2023 11:38 | File folder | |
| 5_CP_pipeline | 23/05/2023 10:07 | File folder | |
| fcs_files | 24/05/2023 22:23 | File folder | |
| 5_CP_pipeline_vero.zip | 24/05/2023 23:03 | Compressed (zipp... | 757,010 KB |
| fcs_files.zip | 24/05/2023 22:51 | Compressed (zipp... | 92,763 KB |
| Notes.txt | 22/05/2023 16:29 | Text Document | 1 KB |

2. Put all raw .tiff images into the folder "1_ImageData"
3. Put the Matlab macros, along with any excel files needed for the macros, in folder "2_image_extraction" and organise according to the image below

| Name | Date modified | Type | Size |
|---|---|---|---|
| Functions | 22/05/2023 16:07 | File folder | |
| Outputs | 22/05/2023 15:49 | File folder | |
| Gut_Data_Analysis_Master.xlsx | 02/05/2023 14:06 | Microsoft Excel W... | 39 KB |
| Gut_Data_Analysis_Master_1stRun.xlsx | 20/04/2023 11:35 | Microsoft Excel W... | 37 KB |
| Number_Name_Conversion.xlsx | 22/05/2023 15:45 | Microsoft Excel W... | 10 KB |
| OPTIMAL_Pipeline_Part_1.m | 24/05/2023 22:03 | MATLAB Code | 22 KB |
| OPTIMAL_Pipeline_Part_2_Heatmaps.m | 14/03/2023 16:08 | MATLAB Code | 14 KB |
| OPTIMAL_Pipeline_Part_3_Visualisaton.m | 14/03/2023 16:45 | MATLAB Code | 13 KB |
| Saskia_MAY23.xlsx | 23/05/2023 15:00 | Microsoft Excel W... | 21 KB |
| spillover2.1.xlsx | 24/05/2023 11:05 | Microsoft Excel W... | 16 KB |

4. Structure folder "3_ilastik_model" to have your ilastik model in the folder, with a folder for your output Pmaps (this must match with what is in the Matlab macro later, so ensure the code matches the information)

| Name | Date modified | Type | Size |
|---|---|---|---|
| OUTPUT_PMAPS | 24/05/2023 12:18 | File folder | |
| 20210906_Tonsil_pixel_classification_v1.0.... | 22/05/2023 16:27 | ilastik project | 35,035 KB |
| 20230220_JD_Nuclear_Model.ilp | 22/02/2023 09:51 | ilastik project | 5,694 KB |
| 20230328_JD_Nuclear_Model.ilp | 22/05/2023 16:36 | ilastik project | 7,542 KB |

5. The folder "4_structured_data_for_CP" should be empty when you initially run the Matlab macro for the first time, the script will populate the folder with other folders for the individual channels in your image and folders for the nuclear, membrane, and probability

maps

| Name | Date modified | Type | Size |
|---|---|---|---|
| Channel_001 | 24/05/2023 12:18 | File folder | |
| Channel_002 | 24/05/2023 12:18 | File folder | |
| Channel_003 | 24/05/2023 12:18 | File folder | |
| Channel_004 | 24/05/2023 12:18 | File folder | |
| Channel_005 | 24/05/2023 12:18 | File folder | |
| Channel_006 | 24/05/2023 12:18 | File folder | |
| Channel_007 | 24/05/2023 12:18 | File folder | |
| Channel_008 | 24/05/2023 12:18 | File folder | |
| Channel_009 | 24/05/2023 12:18 | File folder | |
| Channel_010 | 24/05/2023 12:18 | File folder | |
| Channel_011 | 24/05/2023 12:18 | File folder | |
| Channel_012 | 24/05/2023 12:18 | File folder | |
| Channel_013 | 24/05/2023 12:18 | File folder | |
| Channel_014 | 24/05/2023 12:18 | File folder | |
| Channel_015 | 24/05/2023 12:18 | File folder | |
| Channel_016 | 24/05/2023 12:18 | File folder | |
| Channel_017 | 24/05/2023 12:18 | File folder | |
| Channel_018 | 24/05/2023 12:18 | File folder | |
| Channel_019 | 24/05/2023 12:18 | File folder | |
| Channel_020 | 24/05/2023 12:18 | File folder | |
| Channel_021 | 24/05/2023 12:18 | File folder | |
| Channel_022 | 24/05/2023 12:18 | File folder | |
| Channel_023 | 24/05/2023 12:18 | File folder | |
| Channel_024 | 24/05/2023 12:18 | File folder | |
| Channel_025 | 24/05/2023 12:18 | File folder | |
| Channel_026 | 24/05/2023 12:18 | File folder | |
| Channel_027 | 24/05/2023 12:18 | File folder | |
| Channel_028 | 24/05/2023 12:18 | File folder | |

6. Structure the folder “5_CP_pipeline” with two folders, “OUTPUTS” and “PIPELINE”. The “PIPELINE” folder should only contain the CellProfiler pipeline file to be used with the Matlab macro. The “OUTPUTS” folder should be organised according to the image below. Note, the excel files and the contents of these subfolders will not exist until after the CellProfiler pipeline has completed running

| Name | Date modified | Type | Size |
|---|---|---|---|
| CellMasks | 24/05/2023 19:00 | File folder | |
| Segmentation_Checker | 24/05/2023 19:00 | File folder | |
| Tissue | 24/05/2023 19:00 | File folder | |
| MyExpt_Cells.csv | 24/05/2023 19:17 | Microsoft Excel C... | 4,144,522 KB |
| MyExpt_Experiment.csv | 24/05/2023 19:17 | Microsoft Excel C... | 222 KB |
| MyExpt_Image.csv | 24/05/2023 19:17 | Microsoft Excel C... | 1 KB |
| MyExpt_Nuclei.csv | 24/05/2023 19:24 | Microsoft Excel C... | 1,654,134 KB |
| MyExpt_secondary_objects.csv | 24/05/2023 19:30 | Microsoft Excel C... | 1,654,134 KB |

**Metadata Incorporation**

1. Format your metadata excel file according to the example below. Any data that is planned on being incorporated into the fcs files at the end of the macro must be in numerical format

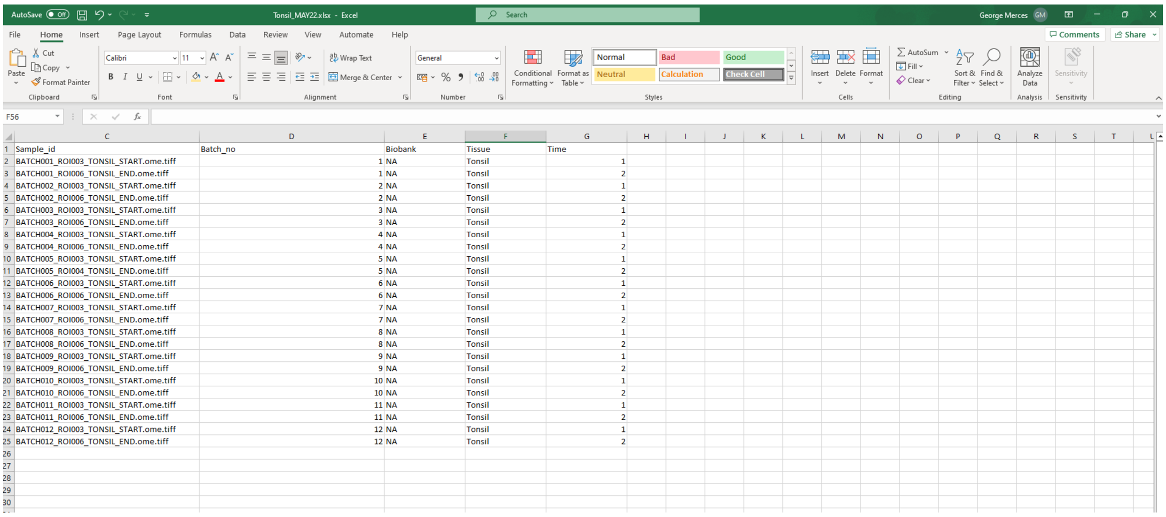

| Sample_id | Batch_no | Biobank | Tissue | Time |
|---|---|---|---|---|
| BATCH001_ROI003_TONSIL_START.ome.tiff | 1 | NA | Tonsil | 1 |
| BATCH001_ROI006_TONSIL_END.ome.tiff | 1 | NA | Tonsil | 2 |
| BATCH002_ROI003_TONSIL_START.ome.tiff | 2 | NA | Tonsil | 1 |
| BATCH002_ROI006_TONSIL_END.ome.tiff | 2 | NA | Tonsil | 2 |
| BATCH003_ROI003_TONSIL_START.ome.tiff | 3 | NA | Tonsil | 1 |
| BATCH003_ROI006_TONSIL_END.ome.tiff | 3 | NA | Tonsil | 2 |
| BATCH004_ROI003_TONSIL_START.ome.tiff | 4 | NA | Tonsil | 1 |
| BATCH004_ROI006_TONSIL_END.ome.tiff | 4 | NA | Tonsil | 2 |
| BATCH005_ROI003_TONSIL_START.ome.tiff | 5 | NA | Tonsil | 1 |
| BATCH005_ROI004_TONSIL_END.ome.tiff | 5 | NA | Tonsil | 2 |
| BATCH006_ROI003_TONSIL_START.ome.tiff | 6 | NA | Tonsil | 1 |
| BATCH006_ROI006_TONSIL_END.ome.tiff | 6 | NA | Tonsil | 2 |
| BATCH007_ROI003_TONSIL_START.ome.tiff | 7 | NA | Tonsil | 1 |
| BATCH007_ROI006_TONSIL_END.ome.tiff | 7 | NA | Tonsil | 2 |
| BATCH008_ROI003_TONSIL_START.ome.tiff | 8 | NA | Tonsil | 1 |
| BATCH008_ROI006_TONSIL_END.ome.tiff | 8 | NA | Tonsil | 2 |
| BATCH009_ROI003_TONSIL_START.ome.tiff | 9 | NA | Tonsil | 1 |
| BATCH009_ROI006_TONSIL_END.ome.tiff | 9 | NA | Tonsil | 2 |
| BATCH010_ROI003_TONSIL_START.ome.tiff | 10 | NA | Tonsil | 1 |
| BATCH010_ROI006_TONSIL_END.ome.tiff | 10 | NA | Tonsil | 2 |
| BATCH011_ROI003_TONSIL_START.ome.tiff | 11 | NA | Tonsil | 1 |
| BATCH011_ROI006_TONSIL_END.ome.tiff | 11 | NA | Tonsil | 2 |
| BATCH012_ROI003_TONSIL_START.ome.tiff | 12 | NA | Tonsil | 1 |
| BATCH012_ROI006_TONSIL_END.ome.tiff | 12 | NA | Tonsil | 2 |

#### Training An Ilastik Model

Note, if you are using the Ilastik model provided, this step may be skipped, and you can just follow the instructions for "OPTIMAL_Pipeline_Part_1.m".

1. Open Ilastik and create a Pixel Classification Model

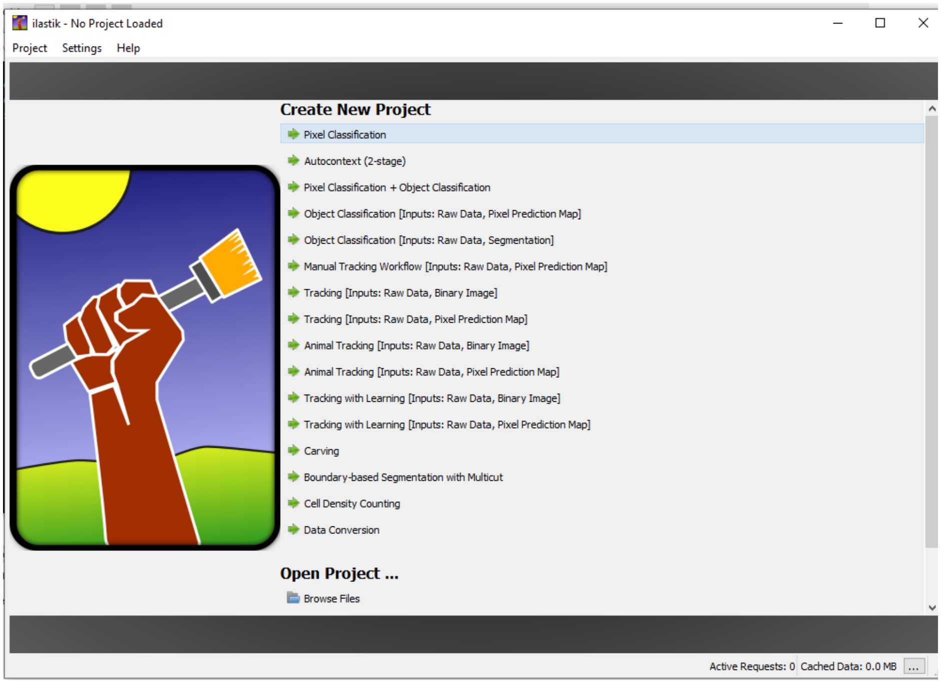


2. Drag a selection of your images (in our case, we use rescaled nuclear channel images) into the images window. Note, for ease of distribution of your model, we recommend setting the image "Location" to "Internal", which increases the file size for your model but ensures it

should be executable by anyone who downloads it.

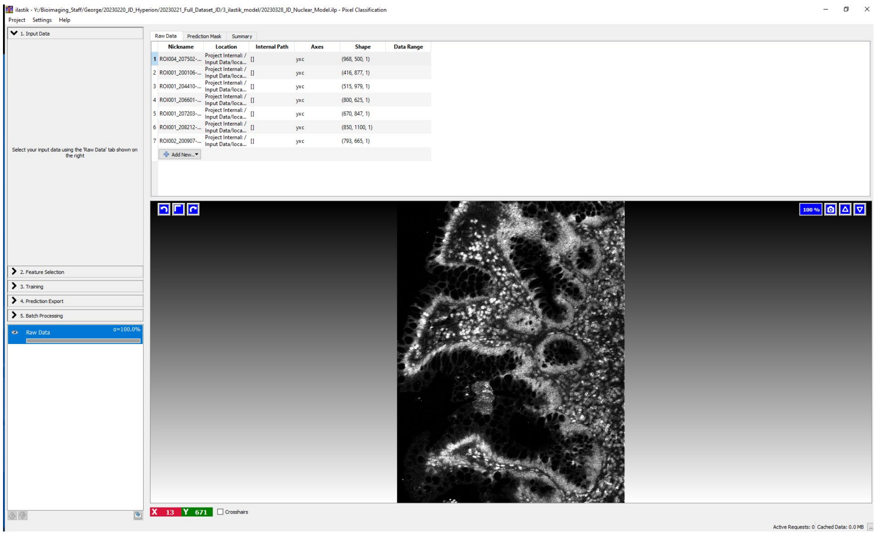

3. Select you Features for use. We recommend selecting a wide range of options

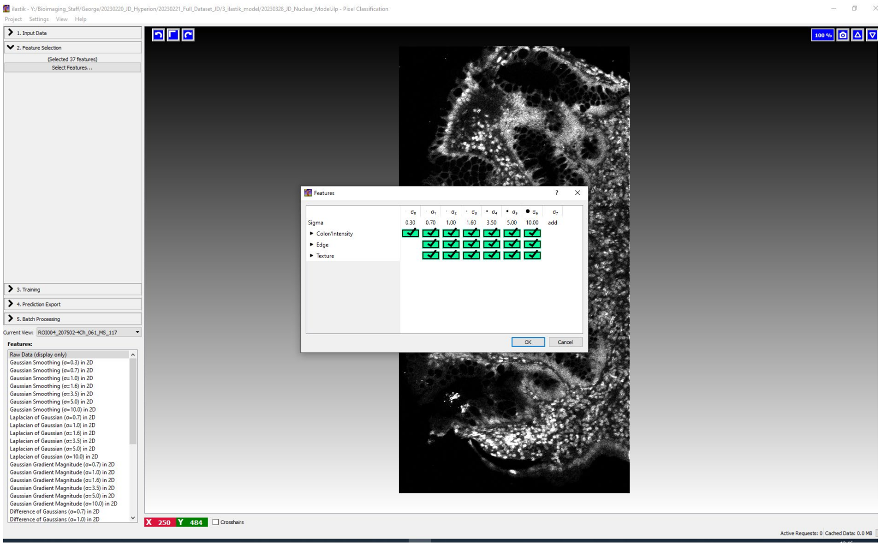

4. Train your model: Rename the label options to be relevant for your sample (e.g. nuclear vs non-nuclear) and label pixels sporadically around your images. Do not label too much, as this can lead to a lack of general ability of the model, and can train it too heavily to the training

dataset.

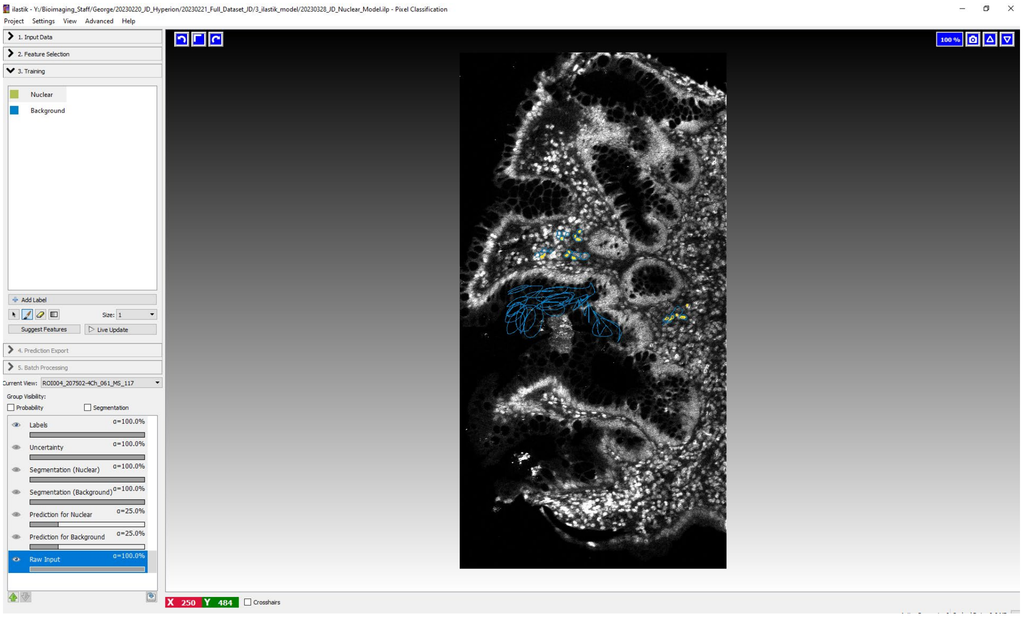


5. Select “Live Update” to train your model. If the output probability is not good enough, try adding additional labelling information to your images, focussing on regions where the probability mapping has made errors

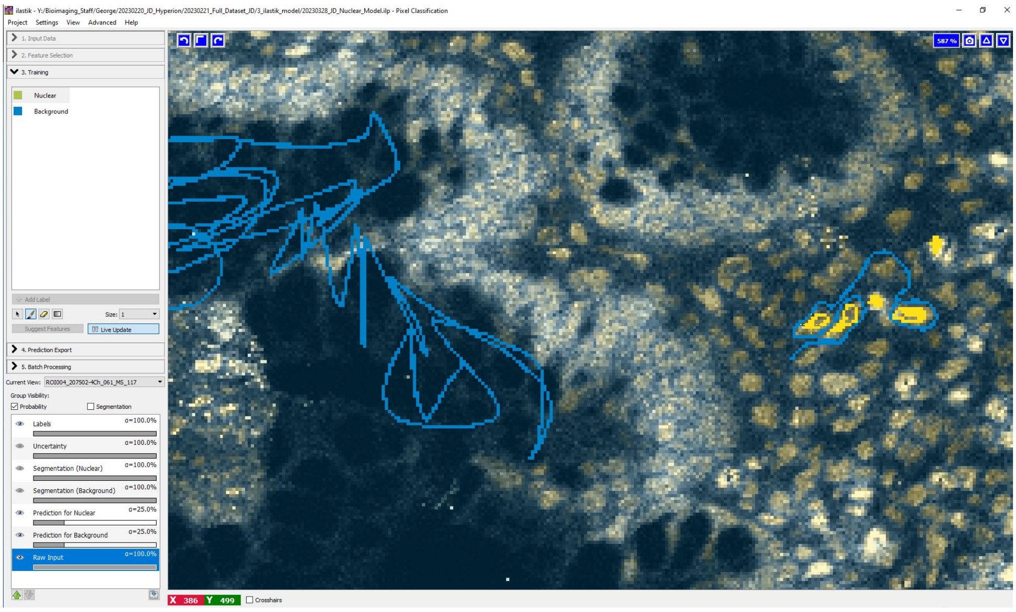


6. Once you’re happy with your probability mapping model, choose the export settings. In the Matlab script given as part of this manuscript, we only export the nuclear option, and we export this to a location close to the Ilastik model file. The exports are then detected and resaved in the appropriate location for CellProfiler with a different name within the Matlab script. Note the name used to save the image under, as this needs to match with the Matlab

script for proper function.

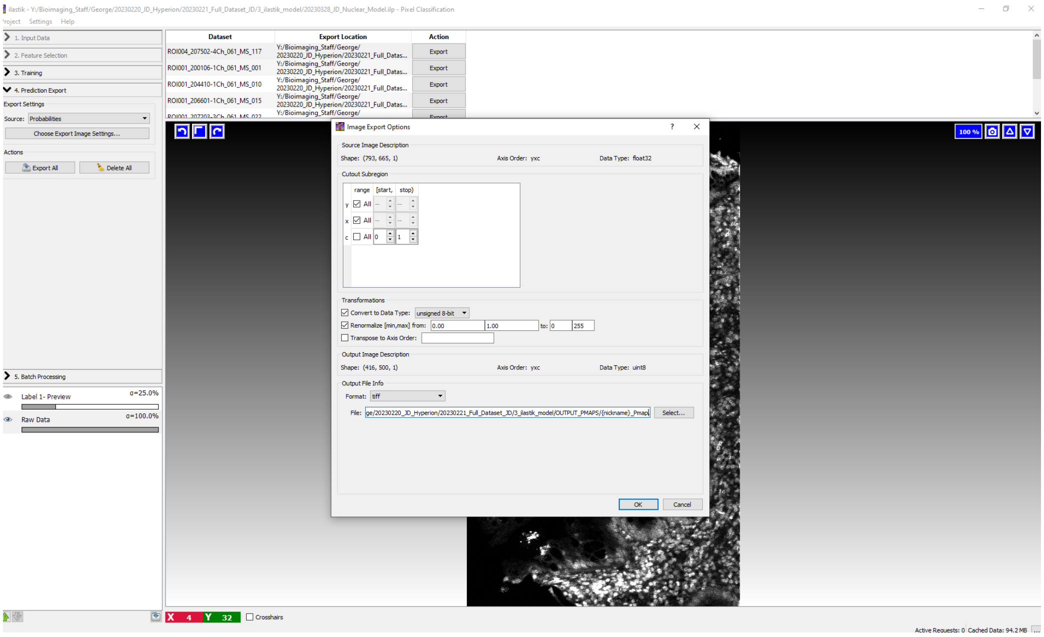


7. Save your finished model to an appropriate location, and make sure this is the location called within the Matlab script to ensure proper macro function

**Matlab Code I**

1. Open Ilastik on your device, select an appropriate folder to save your output probability maps to. Make sure the settings are as listed in the image to ensure correct use of the

images in macro running

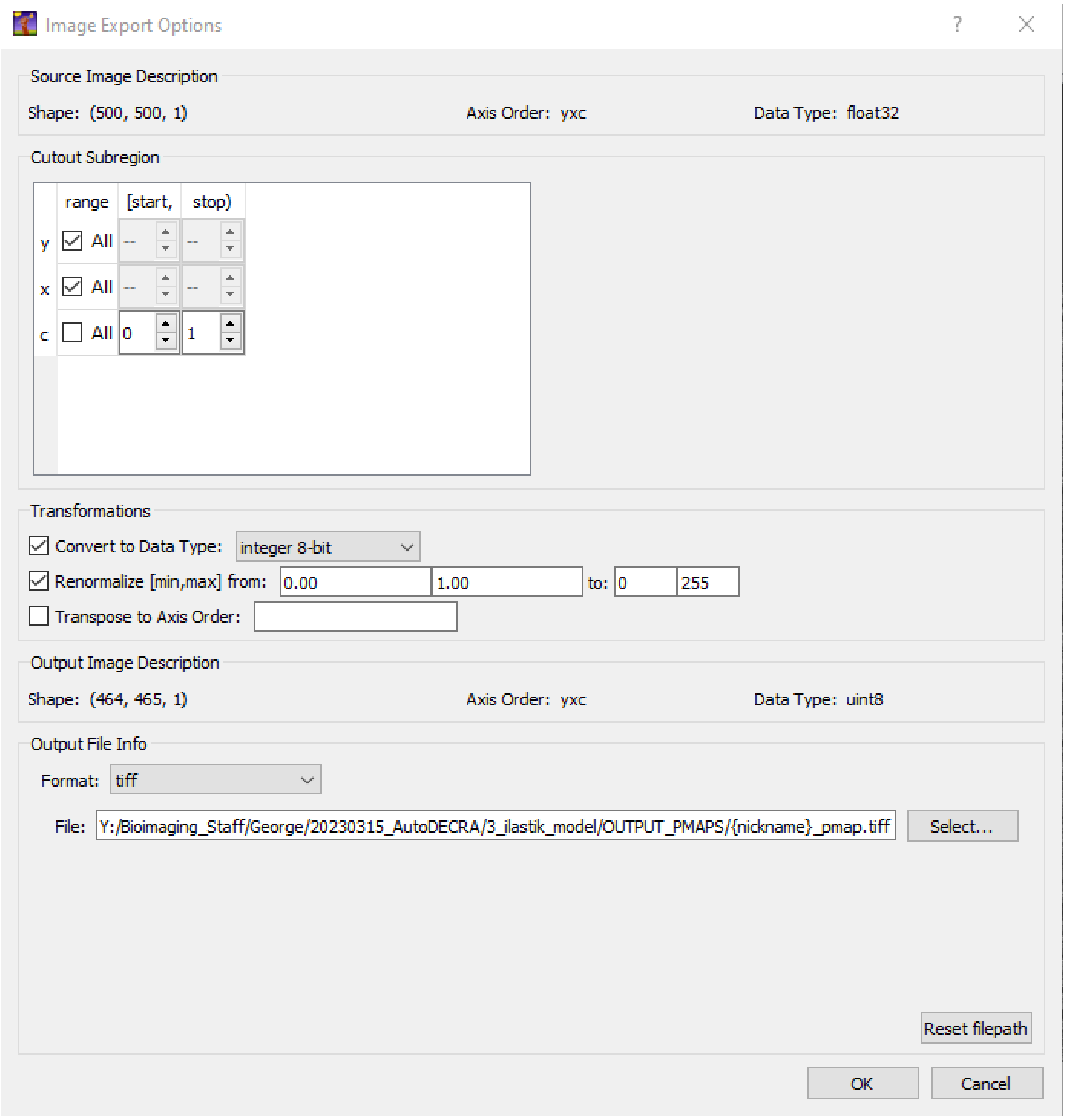

4. Rename the locations for Ilastik and Cell Profiler, using the **exact** format shown in the template code. The spacing and additional features are essential for proper function

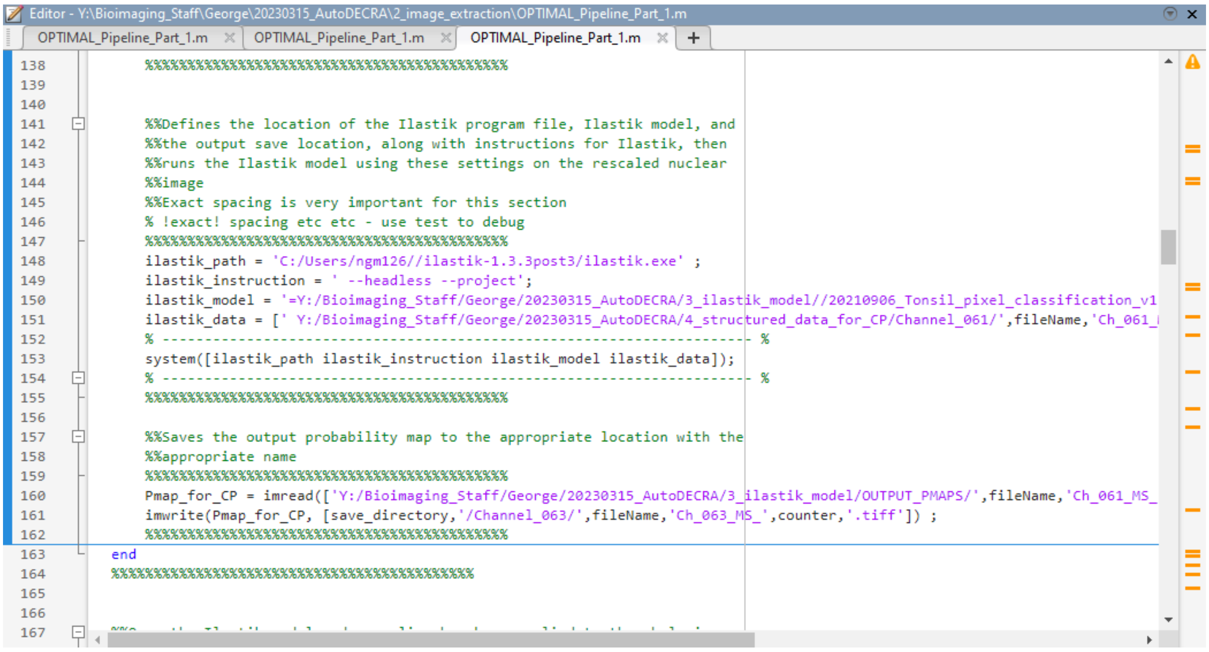


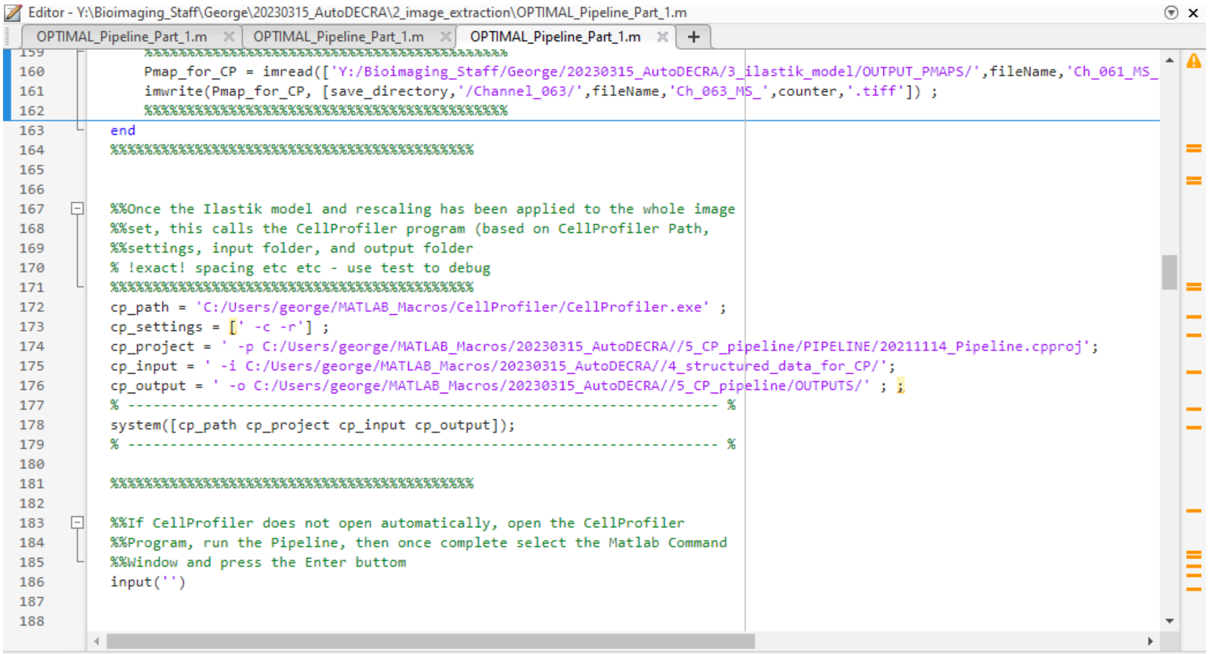


5. Incorporate the names of the excel files to be used for metadata incorporation within the fcs files at the end of the code, including the spillover matrix for compensation of signal bleedthrough.

Editor - Y:\Bioimaging_Staff\George\20230315_AutoDECRA\2_image_extraction\OPTIMAL_Pipeline_Part_1.m

OPTIMAL_Pipeline_Part_1.m | OPTIMAL_Pipeline_Part_1.m | OPTIMAL_Pipeline_Part_1.m

```
%% Simple error check based on number of ROI
if Number_of_masks~=Number_of_ROI
    error('Number of masks and ROI dont match');
end

%% Load compensation matrix and extract important information
%%%%%%%%%%%%%%%%%%%%%%%%%%%%%%%%%%%%%%%%%%%%%%%%%%%%%%%%%%
comp_matrix=readtable('spillover2.1.xlsx');
comp_channels=comp_matrix.Properties.VariableNames;
comp_channels=comp_channels(2:end);
compensation_matrix=table2array(comp_matrix(:,2:end));
%%%%%%%%%%%%%%%%%%%%%%%%%%%%%%%%%%%%%%%%%%%%%%%%%%%%%%%%%%


%% Load the ROI spreadsheet to get the PATIENT metadata
%%%%%%%%%%%%%%%%%%%%%%%%%%%%%%%%%%%%%%%%%%%%%%%%%%%%%%%%%%
ROI_data = readcell('Tonsil_Data_Analysis_Master.xlsx','sheet',4);
Patient_data = readcell('Tonsil_Data_Analysis_Master.xlsx','sheet',5);
Patient_identifier=Patient_data(2:end,2);
ROI_list=ROI_data(2:end,1);
Patient_name_list=ROI_data(2:end,3);
for loop=1:numel(Patient_name_list)
    for loop2=1:numel(Patient_identifier)
        if contains(Patient_name_list(loop),Patient_identifier(loop2)) ==1
            Patient_number_list(loop)=loop2;
        end
    end
end
%%%%%%%%%%%%%%%%%%%%%%%%%%%%%%%%%%%%%%%%%%%%%%%%%%%%%%%%%%
```

Editor - Y:\Bioimaging_Staff\George\20230315_AutoDECRA\2_image_extraction\OPTIMAL_Pipeline_Part_1.m

OPTIMAL_Pipeline_Part_1.m | OPTIMAL_Pipeline_Part_1.m | OPTIMAL_Pipeline_Part_1.m

```
%%Load the spreadsheet containing essential metadata (in this example,
%%Patient, Batch_no, Time, and Tissue)
%%%%%%%%%%%%%%%%%%%%%%%%%%%%%%%%%%%%%%%%%%%%%%%%%%%%%%%%%%
Spreadsheet=readtable('Tonsil_MAY22.xlsx');
Patient_name_list=Spreadsheet.Patient;
Batch_number_list=Spreadsheet.Batch_no;
Patient_identifier=unique(Patient_name_list);
for loop=1:numel(Patient_name_list)
    for loop2=1:numel(Patient_identifier)
        if contains(Patient_name_list(loop),Patient_identifier(loop2)) ==1
            Patient_number_list(loop)=loop2;
        end
    end
end
condition_list=Spreadsheet.Tissue;
%%%%%%%%%%%%%%%%%%%%%%%%%%%%%%%%%%%%%%%%%%%%%%%%%%%%%%%%%%

%%Incorporates the tissue type into the stored data (in this case Tonsil
%%should represent all of the images, with no Syn images
%%%%%%%%%%%%%%%%%%%%%%%%%%%%%%%%%%%%%%%%%%%%%%%%%%%%%%%%%%
Tonsil_ROI_Index = find(contains(condition_list,'Tonsil'));
Syn_ROI_Index = find(contains(condition_list,'Syn'));
% check number should be total number of ROI
check_number=numel(Tonsil_ROI_Index)+numel(Syn_ROI_Index);
Tissue_index=zeros(numel(condition_list),1);
Tissue_index(Tonsil_ROI_Index)=1;
Tissue_index(Syn_ROI_Index)=2;
test=find(Tissue_index==0);   %should be 0 i.e. all ROI allocated a
if test==0
```

6. Incorporate the specific metadata necessary for inclusion within the fcs files later. Make sure this information is purely numeric within your files, otherwise issues with fcs file generation

may occur

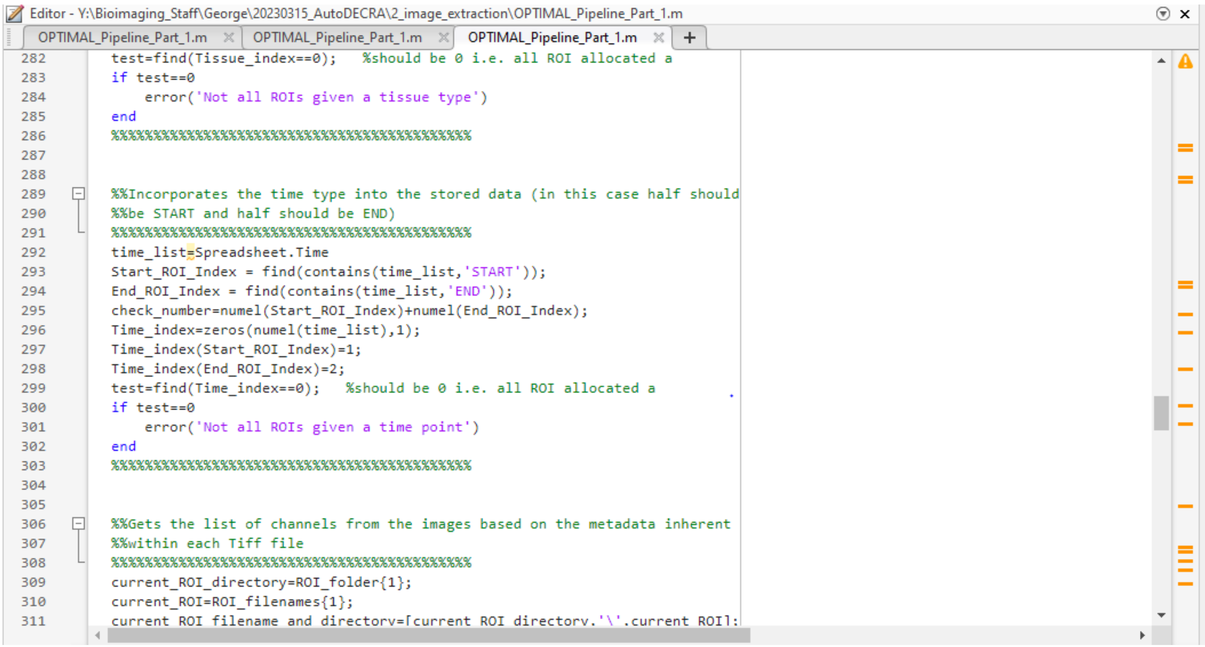


7. Select the channels of interest, i.e. the channels that have been specifically stained and imaged during your experiment (note here that again the first channel is channel 1 etc.)

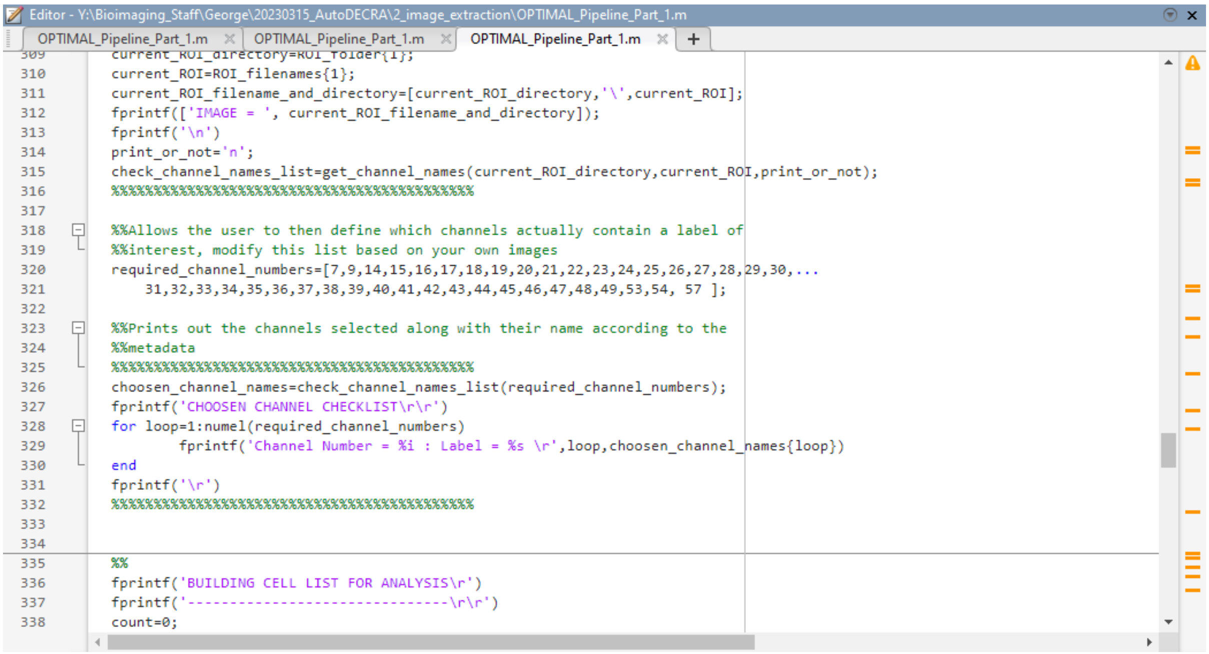

8. Adapt the fcs conversion portion of the code to include the metadata specified earlier, making sure to adjust the headers as well

Editor - Y:\Bioimaging_Staff\George\20230315_AutoDECRA\2_image_extraction\OPTIMAL_Pipeline_Part_1.m

OPTIMAL_Pipeline_Part_1.m | OPTIMAL_Pipeline_Part_1.m | OPTIMAL_Pipeline_Part_1.m

```
        %%Add patient/ROI metadata to single cell file list
        %%%%%%%%%%%%%%%%%%%%%%%%%%%%%%%%%%%%%%%%%%%%
        number_of_cells=size(ONE_ROI_DATA,1);
        all_ROI_list(all_ROI_loop).number_of_cells=number_of_cells;
        Single_cell_ROI_number=repelem(all_ROI_loop,number_of_cells)';
        Single_cell_Patient_number=repelem(Patient_number_list(all_ROI_loop),number_of_cells)';
        Single_cell_File=repelem(Patient_name_list(all_ROI_loop),number_of_cells)';
        Single_cell_Pathology_index=repelem(Tissue_index(all_ROI_loop),number_of_cells)';
        Single_cell_batch_number=repelem(batch_number,number_of_cells)';
        FCS_ROI_DATA=[AS_DATA, AS_NORM_DATA, EXTRA_DATA, Single_cell_ROI_number, Single_cell_Patient_number, Single_cell_Patholo
        if count==1
           COMPLETE_COMP_DATA_SET=[AS_DATA, AS_NORM_DATA, EXTRA_DATA, Single_cell_ROI_number, Single_cell_Patient_number, Single_
        else
           COMPLETE_COMP_DATA_SET=[COMPLETE_COMP_DATA_SET; [AS_DATA, AS_NORM_DATA, EXTRA_DATA, Single_cell_ROI_number, Single_ce
        end
        %%%%%%%%%%%%%%%%%%%%%%%%%%%%%%%%%%%%%%%%%%%%


        %%Creates the fcs file data, and saves as fcs to the appropriate folder
        %%using the appropriate file name
        %%%%%%%%%%%%%%%%%%%%%%%%%%%%%%%%%%%%%%%%%%%%
        fcs_name_all=all_ROI_list(all_ROI_loop).channels_filename;
        [filepath,filename,ext] = fileparts(fcs_name_all);
        fcs_name=[fcs_directory,filename,'.fcs'];
        fcs_channel_list=[marker_channels,norm_marker_channels,extra_markers , {'ROI_number'},{'Patient_number'},{'Pathology_ind
        fprintf('\r')
        fprintf('Writing %s \r',filename)
        fca_writefcs(fcs_name,FCS_ROI_DATA,fcs_channel_list, fcs_channel_list)
        %%%%%%%%%%%%%%%%%%%%%%%%%%%%%%%%%%%%%%%%%%%%

```

9. Repeat this process for the "collated fcs" file at the very end of the macro

Editor - Y:\Bioimaging_Staff\George\20230315_AutoDECRA\2_image_extraction\OPTIMAL_Pipeline_Part_1.m

OPTIMAL_Pipeline_Part_1.m | OPTIMAL_Pipeline_Part_1.m | OPTIMAL_Pipeline_Part_1.m

```
        %%%%%%%%%%%%%%%%%%%%%%%%%%%%%%%%%%%%%%%%%%%%
        fcs_name_all=all_ROI_list(all_ROI_loop).channels_filename;
        [filepath,filename,ext] = fileparts(fcs_name_all);
        fcs_name=[fcs_directory,filename,'.fcs'];
        fcs_channel_list=[marker_channels,norm_marker_channels,extra_markers , {'ROI_number'},{'Patient_number'},{'Pathology_ind
        fprintf('\r')
        fprintf('Writing %s \r',filename)
        fca_writefcs(fcs_name,FCS_ROI_DATA,fcs_channel_list, fcs_channel_list)
        %%%%%%%%%%%%%%%%%%%%%%%%%%%%%%%%%%%%%%%%%%%%

    end
    %%%%%%%%%%%%%%%%%%%%%%%%%%%%%%%%%%%%%%%%%%%%

    %%Creates one final fcs file containing all the data processed within this
    %%pipeline
    %%%%%%%%%%%%%%%%%%%%%%%%%%%%%%%%%%%%%%%%%%%%
    fprintf('\r')
    filename='Combined_dataset';
    fcs_name=[fcs_directory,filename,'.fcs'];
    fprintf('Writing %s \r',filename)
    fcs_channel_list=[marker_channels,norm_marker_channels,extra_markers , {'ROI_number'},{'Patient_number'},{'Pathology_index'}
    fca_writefcs(fcs_name,COMPLETE_COMP_DATA_SET,fcs_channel_list, fcs_channel_list)
    %%%%%%%%%%%%%%%%%%%%%%%%%%%%%%%%%%%%%%%%%%%%

    fprintf('\r\r')

    %%End of Macro
```

10. Run the script. There is a manual input stage following CellProfiler calling to check that the script has been effective before attempting to continue. If CellProfiler has failed to run, it is possible to run CellProfiler manually according to the instructions in the following section and to then press "Enter" on the keypad back in Matlab to continue the script.

#### CellProfiler Manual Use

If CellProfiler does not run, you may manually open the CellProfiler software from outside Matlab and run according to the instructions below

1. Drag all image folders from the folder “4_structured_data_for_CP” into the images window. Note, this system is set up for 60 channel images, and modifications will need to be made if a different number of channels are actually present.

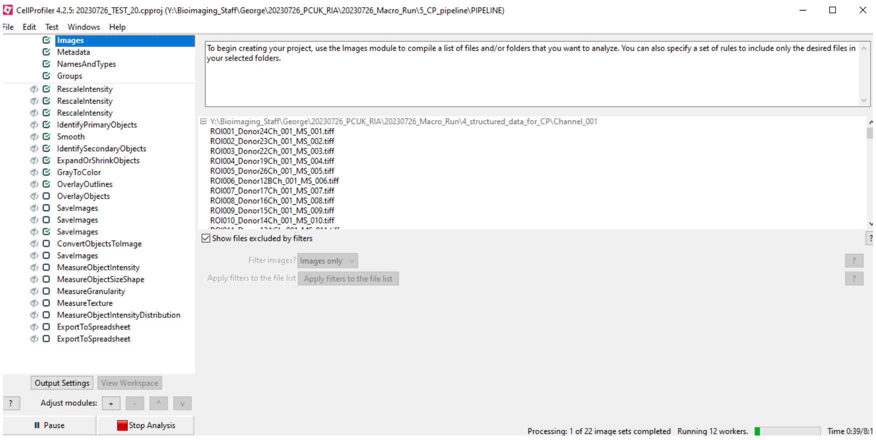


2. Modify the save locations for all “SaveImages” modules, along with the “ExportToSpreadsheet” module if you plan on using that data.

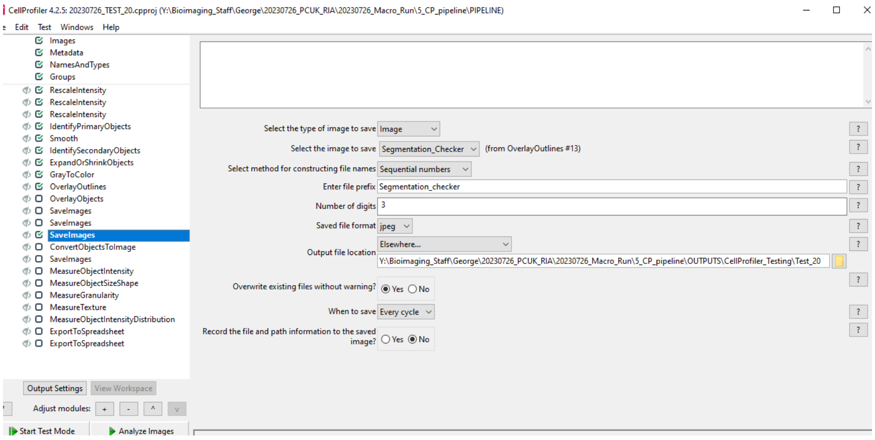


3. When ready to run the pipeline, simply press “Analyse Images” and wait for the pipeline to finish before returning to Matlab

#### Clustering Data Excel

1. Structure your clustering data according to the image, with the original cluster assignment in column "cSOM_s" and the new cluster assignment in column "Tier2ClusterAssisgnment" or "Tier1ClusterAssignment" depending on the clustering tier. Put the name of the cluster in column "Name".

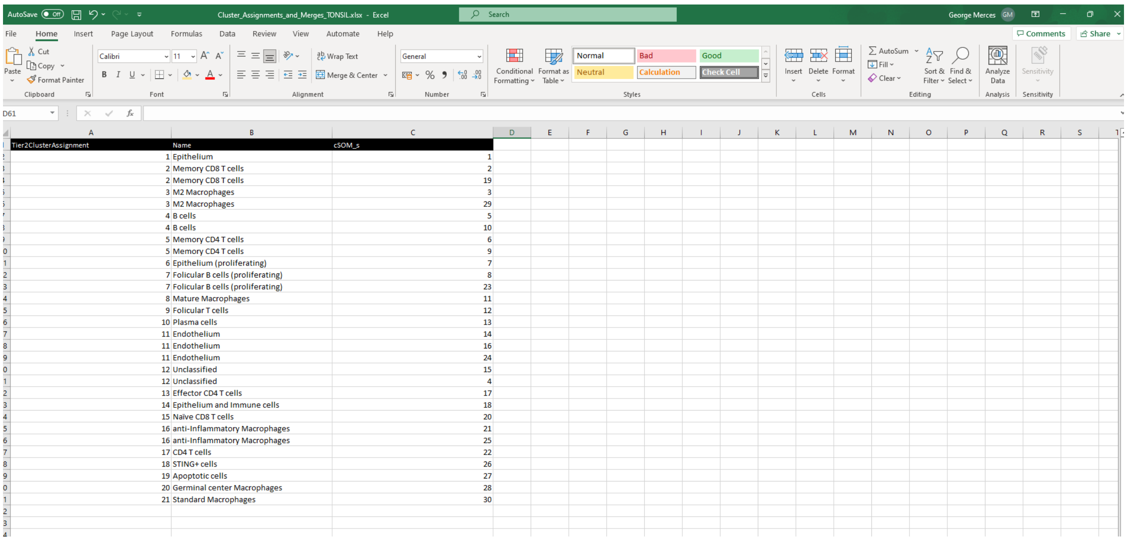

| Tier2ClusterAssignment | Name | cSOM_s |
|---|---|---|
| 1 | Epithelium | 1 |
| 2 | Memory CD8 T cells | 2 |
| 2 | Memory CD8 T cells | 19 |
| 3 | M2 Macrophages | 3 |
| 3 | M2 Macrophages | 29 |
| 4 | B cells | 5 |
| 4 | B cells | 10 |
| 5 | Memory CD4 T cells | 6 |
| 5 | Memory CD4 T cells | 9 |
| 6 | Epithelium (proliferating) | 7 |
| 7 | Folicular B cells (proliferating) | 8 |
| 7 | Folicular B cells (proliferating) | 23 |
| 8 | Mature Macrophages | 11 |
| 9 | Folicular T cells | 12 |
| 10 | Plasma cells | 13 |
| 11 | Endothelium | 14 |
| 11 | Endothelium | 16 |
| 11 | Endothelium | 24 |
| 12 | Unclassified | 15 |
| 12 | Unclassified | 4 |
| 13 | Effector CD4 T cells | 17 |
| 14 | Epithelium and Immune cells | 18 |
| 15 | Naïve CD8 T cells | 20 |
| 16 | anti-Inflammatory Macrophages | 21 |
| 16 | anti-Inflammatory Macrophages | 25 |
| 17 | CD4 T cells | 22 |
| 18 | STING+ cells | 26 |
| 19 | Apoptotic cells | 27 |
| 20 | Germinal center Macrophages | 28 |
| 21 | Standard Macrophages | 30 |

#### Matlab Macro II

1. Following cluster analysis, it's time to run the Matlab Macro "OPTIMAL_Pipeline_Part_2_Heatmaps.m". Update the name for the csv file containing details about each cell identified in the process, along with the folder path names for the Mask outputs from CellProfiler, along with where you would like output heatmaps to be saved.

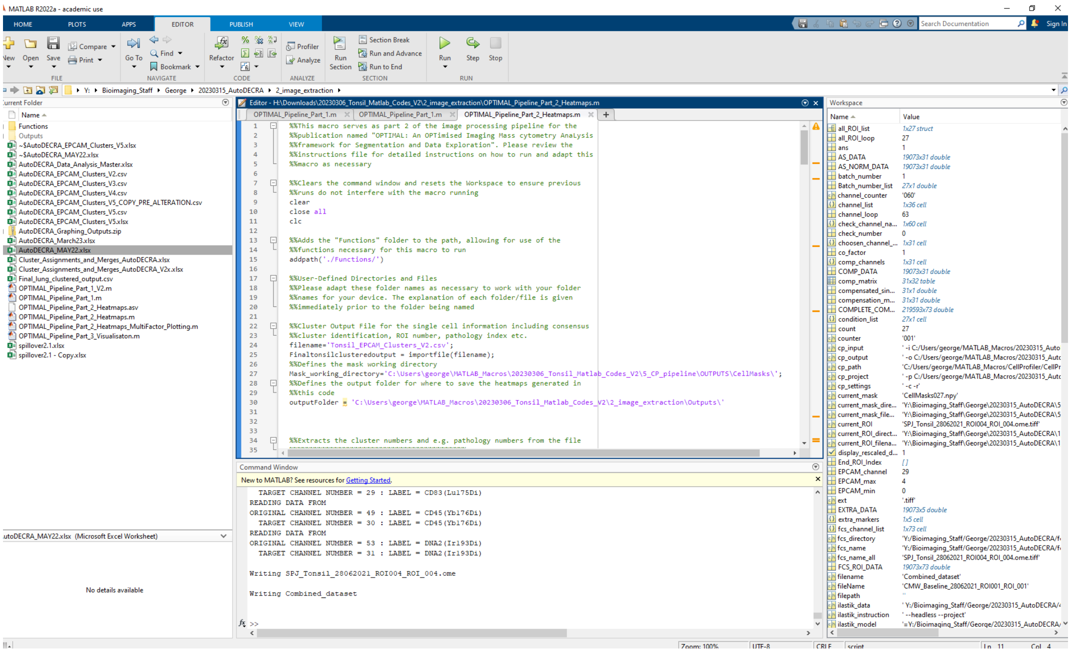

2. Update the name for the clustering information file that you have for your data, including the names of relevant sheets for specific clustering tiers if necessary. Make sure to adapt the "TIER_TO_USE" parameter to the relevant tier for your data

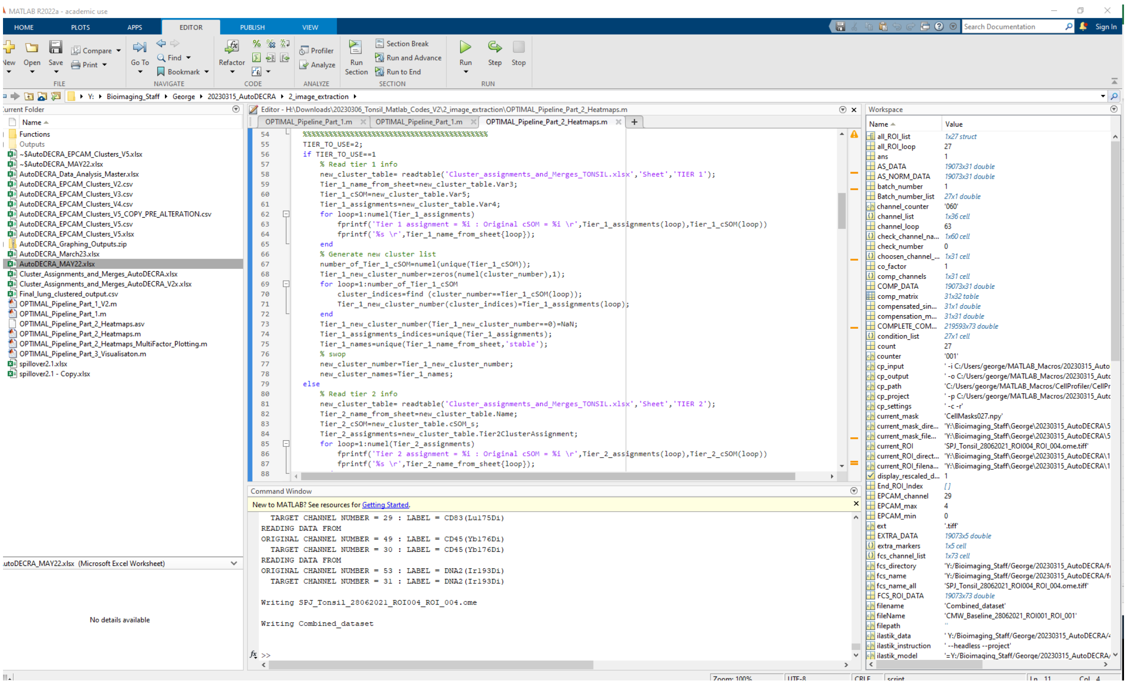

3. Adapt the settings for the heatmap generation in the section shown below. The key parameters are "pVal_sig" which sets the threshold for what proportion of the simulation images need to show positive, neutral, or negative interactions relative to the original dataset to be determined as a significant positive, neutral, or negative interaction.

“cut_off_percent” determines the proportion of images a cell type needs to be present in to be counted within the analysis. “permutations” refers to the number of simulation images generated. This is important in conjunction with the “pVal_sig” value, as too few permutations with a high significance value could lead to aberrant results. Finally “number_of_pixels” refers to the number of pixels a cell mask is expanded by to determine neighbouring cells, the smaller this number the smaller the region the macro will assess to identify neighbours. This number should be adjusted based on cell density and cell size.

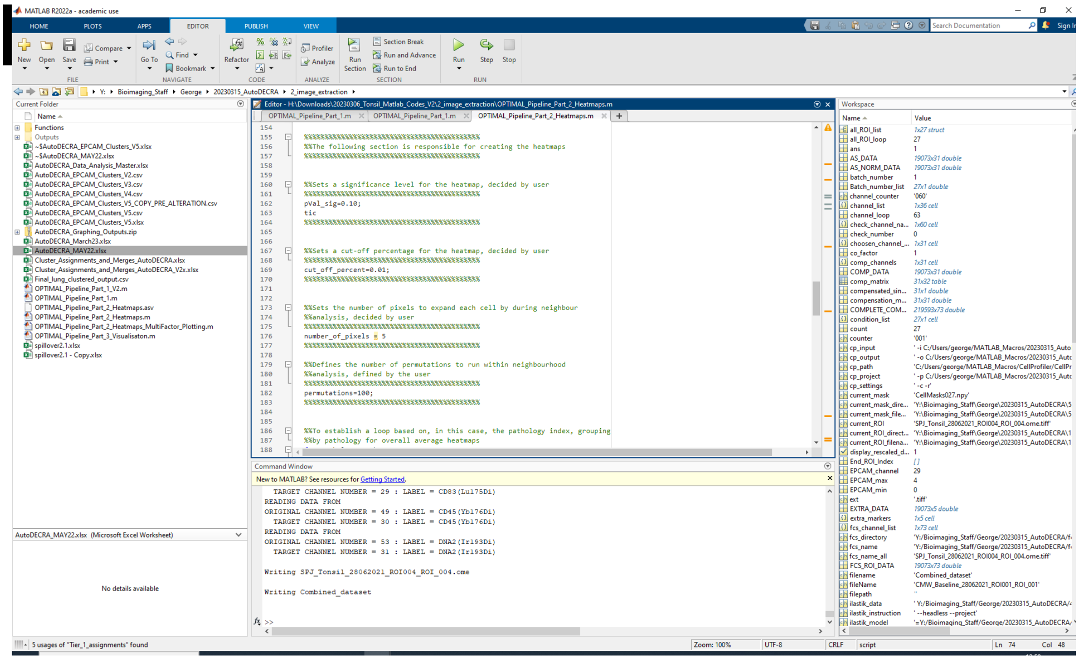

4. Create heatmaps based on clinical/metadata information by using “for” loops: If you have metadata information that you wish to group your data by for heatmap generation, e.g. for comparison between pathologies, you can use this data as long as it is present within your single cell csv file.

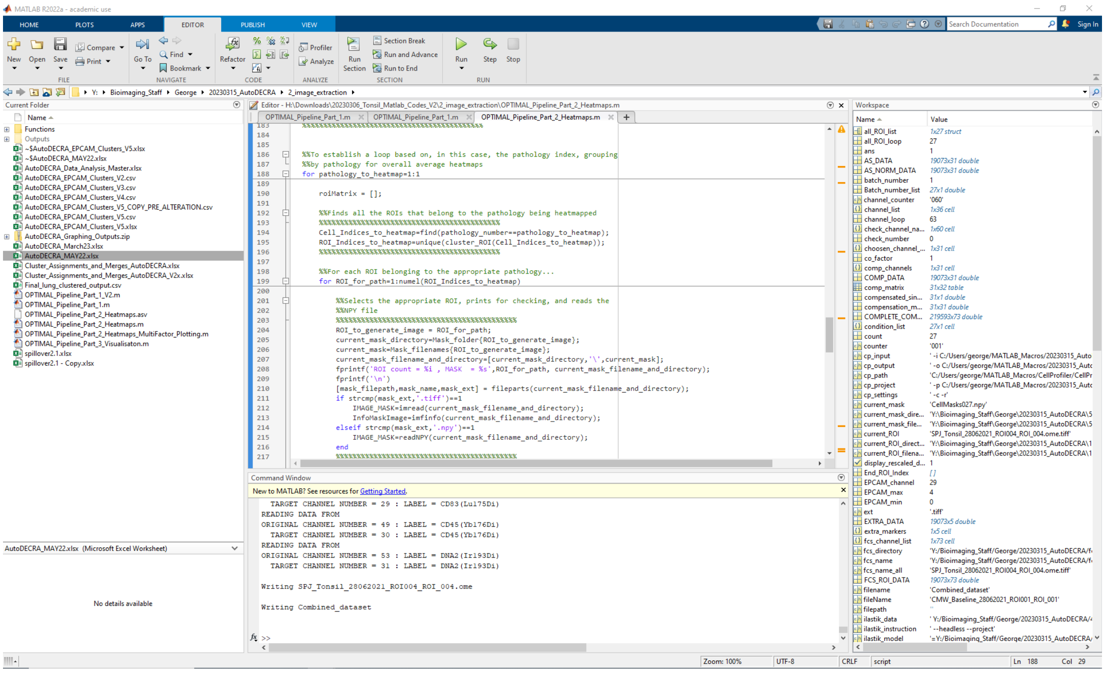

5. Set a size gate on the cells to be analysed by adapting the lower and higher area gate parameters

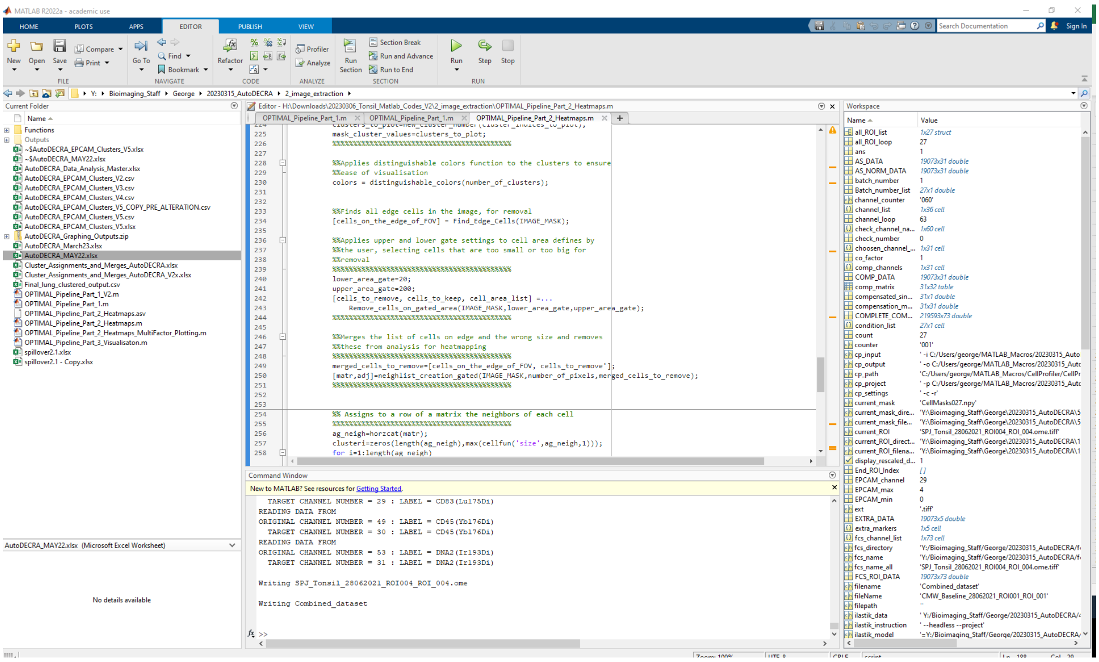

6. Adjust the save name for your heatmaps to reflect any specific alterations you are trialling or testing. For example, as default we have the heatmaps saved specifying the pathology and the pixel number for cell expansion, but you could incorporate any of the other parameters used within your heatmap generation.

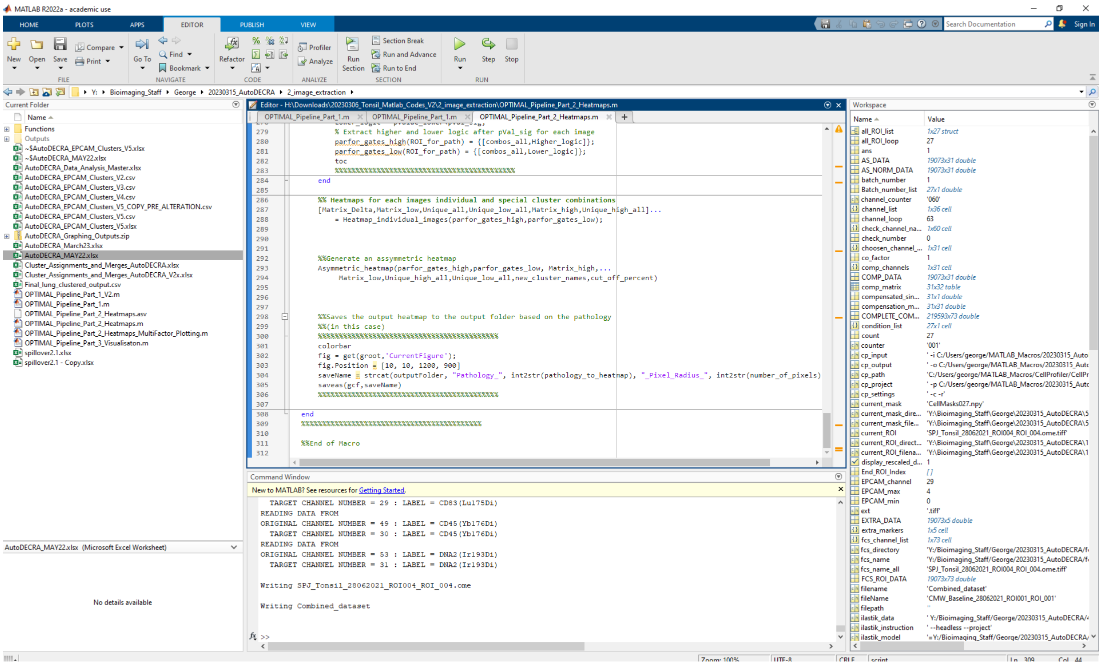

7. Run the script

#### Matlab Script III

1. To create colour-coded cell masks of your data based on cell type information, use the Matlab script “OPTIMAL_Pipeline_Part_3_Visualisaton.m”. First, adapt the folder paths for the output figure location, the csv file containing every cell with a specific cluster assignment, the directory containing the cell mask npy files from CellProfiler, and the raw image folder (for save name creation)

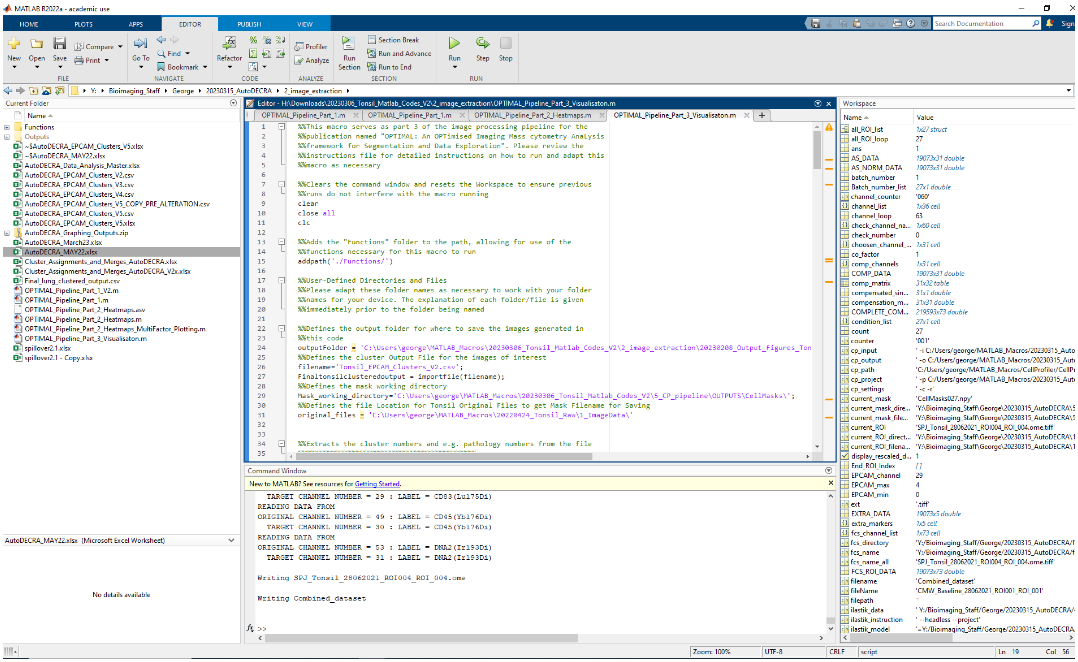

2. Update the name for the clustering information file that you have for your data, including the names of relevant sheets for specific clustering tiers if necessary. Make sure to adapt the “TIER_TO_USE” parameter to the relevant tier for your data

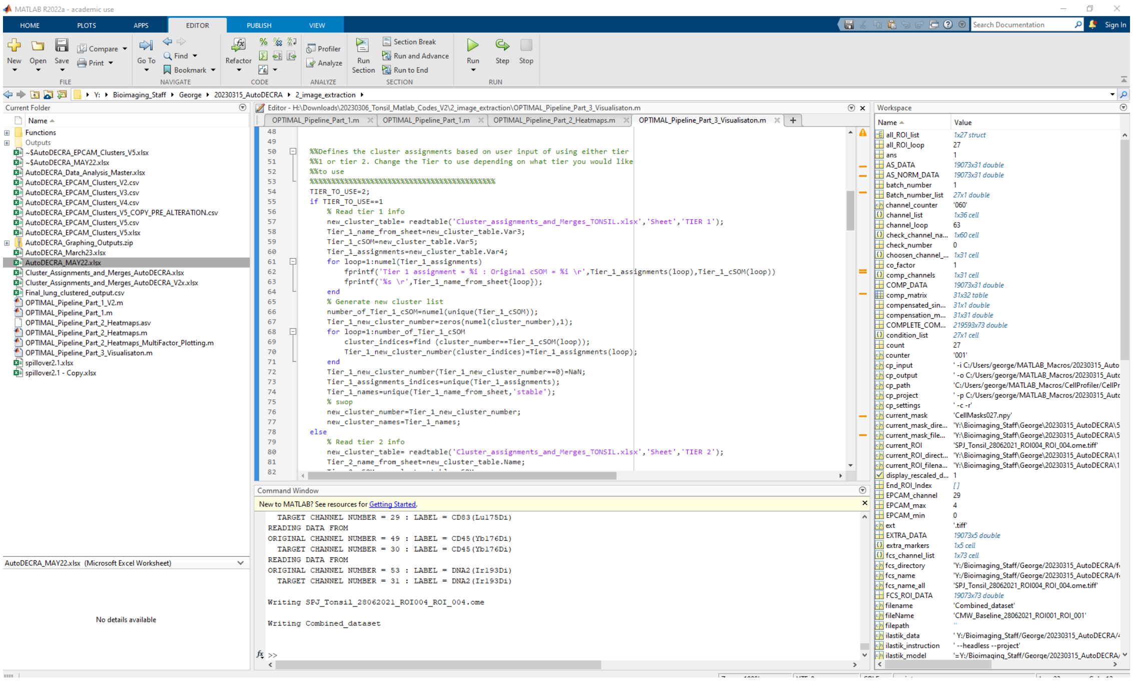

3. If you would like to plot only selected cell type clusters, adapt the list of clusters in the array “selected_clusters_to_plot” to only include clusters of interest

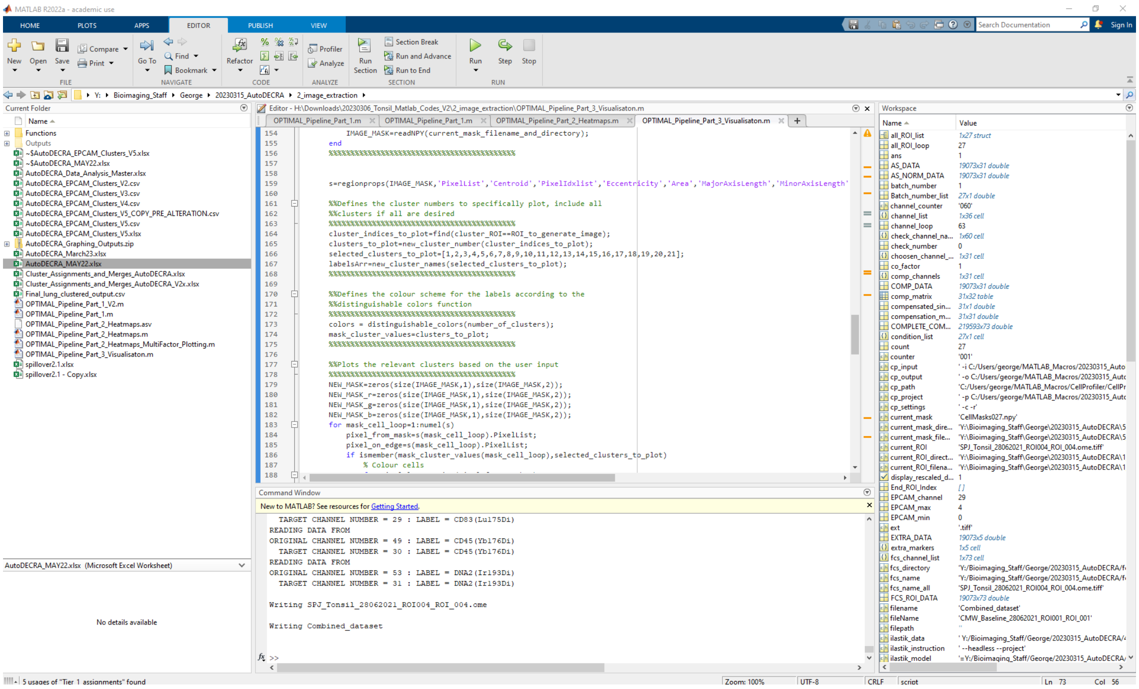

4. To incorporate additional information into the save name of the files, update the parameter “saveName” in the macro, by default we include the file name along with the clustering tier specified by the user

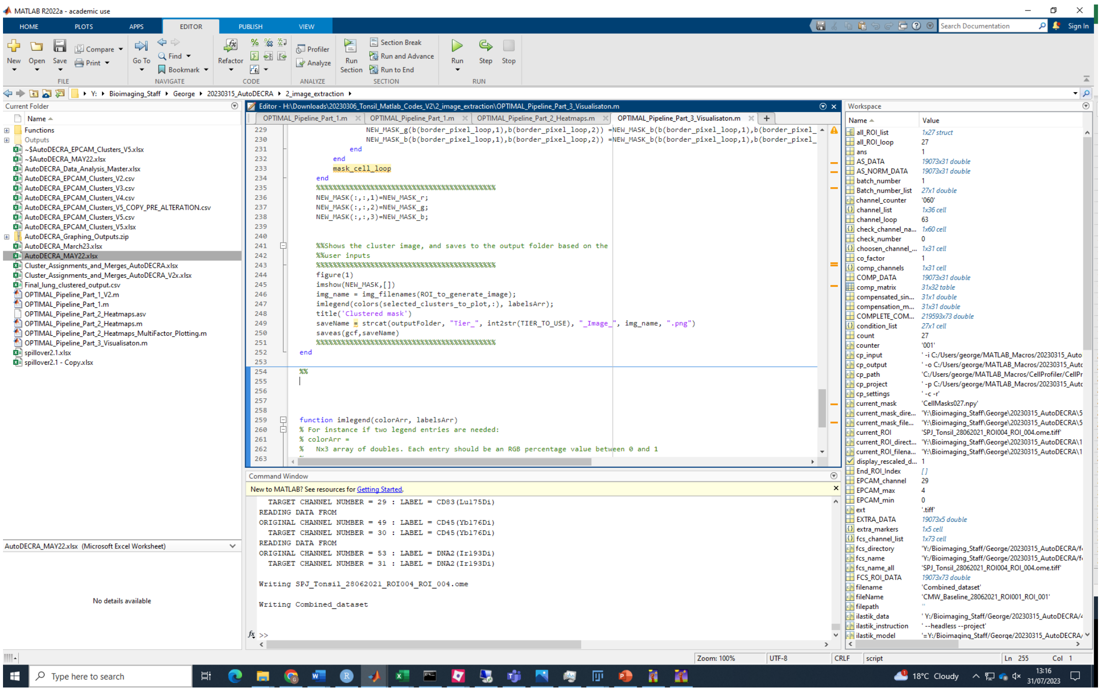

5. Run the script

**References within supplemental notes**


1. Mei HE, Leipold MD, Maecker HT. Platinum-conjugated antibodies for application in mass cytometry. Cytometry A 2016;89:292-300.
2. Chevrier S, Crowell HL, Zanotelli VRT, Engler S, Robinson MD, Bodenmiller B. Compensation of Signal Spillover in Suspension and Imaging Mass Cytometry. Cell Syst 2018;6:612-620 e5.
3. Berg S, Kutra D, Kroeger T, Straehle CN, Kausler BX, Haubold C, Schiegg M, Ales J, Beier T, Rudy M and others. ilastik: interactive machine learning for (bio)image analysis. Nat Methods 2019;16:1226-1232.
4. Carpenter AE, Jones TR, Lamprecht MR, Clarke C, Kang IH, Friman O, Guertin DA, Chang JH, Lindquist RA, Moffat J and others. CellProfiler: image analysis software for identifying and quantifying cell phenotypes. Genome Biol 2006;7:R100.
5. Stirling DR, Swain-Bowden MJ, Lucas AM, Carpenter AE, Cimini BA, Goodman A. CellProfiler 4: improvements in speed, utility and usability. BMC Bioinformatics 2021;22:433.
6. Burbidge JB, Magee L, Robb AL. Alternative Transformations to Handle Extreme Values of the Dependent Variable. Journal of the American Statistical Association 1988;83:123-127.
7. Lin TH, Li HT, Tsai KC. Implementing the Fisher's discriminant ratio in a k-means clustering algorithm for feature selection and data set trimming. J Chem Inf Comput Sci 2004;44:76-87.
8. Bruce Bagwell C. High-Dimensional Modeling for Cytometry: Building Rock Solid Models Using GemStone and Verity Cen-se' High-Definition t-SNE Mapping. Methods Mol Biol 2018;1678:11-36.
9. Wang Y. Understanding How Dimension Reduction Tools Work: An Empirical Approach to Deciphering t-SNE, UMAP, TriMap, and PaCMAP for Data Visualization Journal of Machine Learning Research 2021.
10. Van Gassen S, Callebaut B, Van Helden MJ, Lambrecht BN, Demeester P, Dhaene T, Saeys Y. FlowSOM: Using self-organizing maps for visualization and interpretation of cytometry data. Cytometry A 2015;87:636-45.
11. Levine JH, Simonds EF, Bendall SC, Davis KL, Amir EAD, Tadmor MD, Litvin O, Fienberg HG, Jager A, Zunder ER and others. Data-Driven Phenotypic Dissection of AML Reveals Progenitor-like Cells that Correlate with Prognosis. Cell 2015;162:184-197.
12. Schapiro D, Jackson HW, Raghuraman S, Fischer JR, Zanotelli VRT, Schulz D, Giesen C, Catena R, Varga Z, Bodenmiller B. histoCAT: analysis of cell phenotypes and interactions in multiplex image cytometry data. Nat Methods 2017;14:873-876.

### Table S1

| Antibody | Clone | Vendor | Cat. # | Metal Tag | Working Conc. (ug/mL) | Major target cell type/population | Main spatial location (Protein Atlas) |
|---|---|---|---|---|---|---|---|
| CD45RO | UCHL1 | Thermofisher | CAT#14-0457-82 | 115In | 4 | Activated/memory immune cells | Non-follicular, mantle zone |
| CD45RA | HI100 | Thermofisher | CAT#14-0458-82 | 141Pr | 4 | Naïve immune cells | Widespread locations |
| CD68 | KP1 | Biolegend | CAT#372902 | 142Nd | 5 | Macrophages/monocytes | Widespread locations |
| CD8a | C8/144B | Biolegend | CAT#372902 | 143Nd | 6 | CD8 T cells | Non-follicular, mantle zone |
| KI67 | Polyclonal | Novus | CAT#NB500-170 | 144Nd | 3 | Proliferating/mitotic cells | Follicles/GCs |
| Collagen I | 3D5E8 | Protein Tech | CAT#66761-1-Ig | 145Nd | 2.5 | Fibroblasts | Epithelium |
| CD138 | 4F3A8 | Protein Tech | CAT#67155-1-Ig | 146Nd | 3.5 | Plasma B cells | Non-follicular, mantle zone |
| CD163 | EDHu-1 | BioRad | CAT#MCA1853 | 147Sm | 10 | Perifollicular macrophages | Epithelium and mantle zone |
| MPO | 4C11F6 | Protein Tech | CAT#66177-1-Ig | 150Nd | 5 | Neutrophils | Non-follicular, mantle zone |
| CD56 | E7X9M | CST | CAT#99746BF | 153Eu | 12 | NK/NK T cells | Non-follicular, mantle zone |
| CD69 | 15B5G2 | Novus | CAT#NBP2-25236 | 155Gd | 0.25 | Memory T cells | Epithelium |
| EPCAM | Polyclonal | Abcam | CAT#ab71916 | 156Gd | 1 | Epithelial cells | Widespread locations |
| CD206 | 2A6A10 | Abcam | 60143-1-Ig | 157Gd | 0.5 | Tissue Macrophages and DCs | Epithelium and mantle zone |
| CD79a | EP3618 | Abcam | CAT#ab239891 | 158Gd | 7.5 | B cells (all) | Widespread locations |
| STING | D2P2F | CST | CAT#13647 | 159Tb | 5 | T and NK cell subsets | Widespread locations |
| CD1c | 2A7C11 | Novus | CAT#NBP2-61726 | 162Dy | 2.5 | Dendritic cells (DCs) | Epithelium |
| IFITM3 | Polyclonal | Protein Tech | CAT#11714-1-AP | 163Dy | 5 | Immune cells | Epithelium and mantle zone |
| CD57 | HNK-1 | Biolegend | CAT#359602 | 165Ho | 10 | GC-resident T/B cells | Follicles/GCs |
| IL6-R | Polyclonal | Thermofisher | CAT#PA5-100836 | 167Er | 10 | Immune cells | Widespread locations |
| Cleaved Caspase-3 | Asp175 | CST | CAT#9579S | 168Er | 5 | Apoptotic cells | Follicles/GCs |
| CD3 | Polyclonal | Fluidigm | CAT#3170019D | 170Er | 7.5 | T/NKT Cells | Widespread locations |
| CD31 | EPR3094 | Abcam | CAT#ab207090 | 172Yb | 2 | Endothelium | Epithelium and mantle zone |
| CD4 | EPR6855 | Abcam | CAT#ab181724 | 174Yb | 6 | CD4 T cells/Monocytes | Widespread locations |
| HLA-DR | LN3 | Thermofisher | CAT#14-9956-82 | 175Lu | 5 | Antigen presenting cells (MHCII) | Widespread locations |
| CD169 | SP213 | Abcam | CAT#ab245735 | 176Yb | 2.5 | Monocytes | Widespread locations |
| CD147 | E1S1V | CST | CAT#13287BF | 194Pt | 4 | Various cells | Widespread locations |
| Beta-2-M | EPR21752-214 | Abcam | CAT#ab237032 | 198Pt | 1.5 | All cells (MHC-I) | Widespread locations |

CST = Cell Signalling Technologies; GC – Germinal centres

### Figure S1

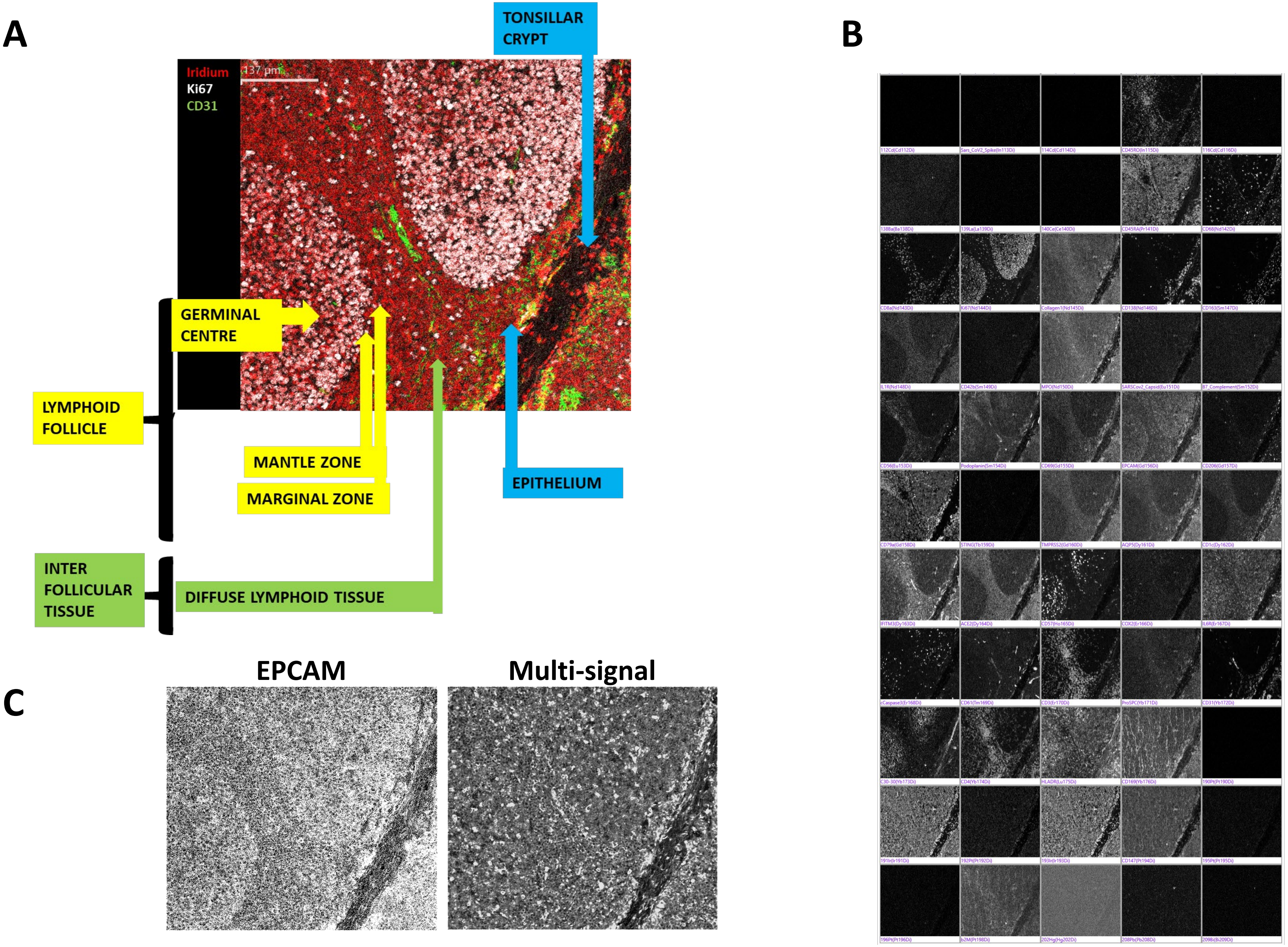
A
Iridium
Ki67
CD31
137 µm
TONSILLAR CRYPT
GERMINAL CENTRE
LYMPHOID FOLLICLE
MANTLE ZONE
MARGINAL ZONE
EPITHELIUM
INTER FOLLICULAR TISSUE
DIFFUSE LYMPHOID TISSUE
B
112Cd(Cd112Di)
Sars_CoV2_Spike(In113Di)
114Cd(Cd114Di)
CD45RO(In115Di)
116Cd(Cd116Di)
138Ba(Ba138Di)
139La(La139Di)
140Ce(Ce140Di)
CD45RA(Pr141Di)
CD68(Nd142Di)
CD8a(Nd143Di)
Ki67(Nd144Di)
Collagen1(Nd145Di)
CD138(Nd146Di)
CD163(Sm147Di)
IL1R(Nd148Di)
CD42b(Sm149Di)
MPO(Nd150Di)
SARSCov2_Capsid(Eu151Di)
B7_Complement(Sm152Di)
CD56(Eu153Di)
Podoplanin(Sm154Di)
CD69(Gd155Di)
EPCAM(Gd156Di)
CD206(Gd157Di)
CD79a(Gd158Di)
STING(Tb159Di)
TMPRSS2(Gd160Di)
AQP5(Dy161Di)
CD1c(Dy162Di)
IFITM3(Dy163Di)
ACE2(Dy164Di)
CD57(Ho165Di)
COX2(Er166Di)
IL6R(Er167Di)
cCaspase3(Er168Di)
CD61(Tm169Di)
CD3(Er170Di)
ProSPC(Yb171Di)
CD31(Yb172Di)
C30-30(Yb173Di)
CD4(Yb174Di)
HLADR(Lu175Di)
CD169(Yb176Di)
190Pt(Pt190Di)
191Ir(Ir191Di)
192Pt(Pt192Di)
193Ir(Ir193Di)
CD147(Pt194Di)
195Pt(Pt195Di)
196Pt(Pt196Di)
b2M(Pt198Di)
202Hg(Hg202Di)
208Pb(Pb208Di)
209Bi(Bi209Di)
C
EPCAM
Multi-signal

### Figure S2

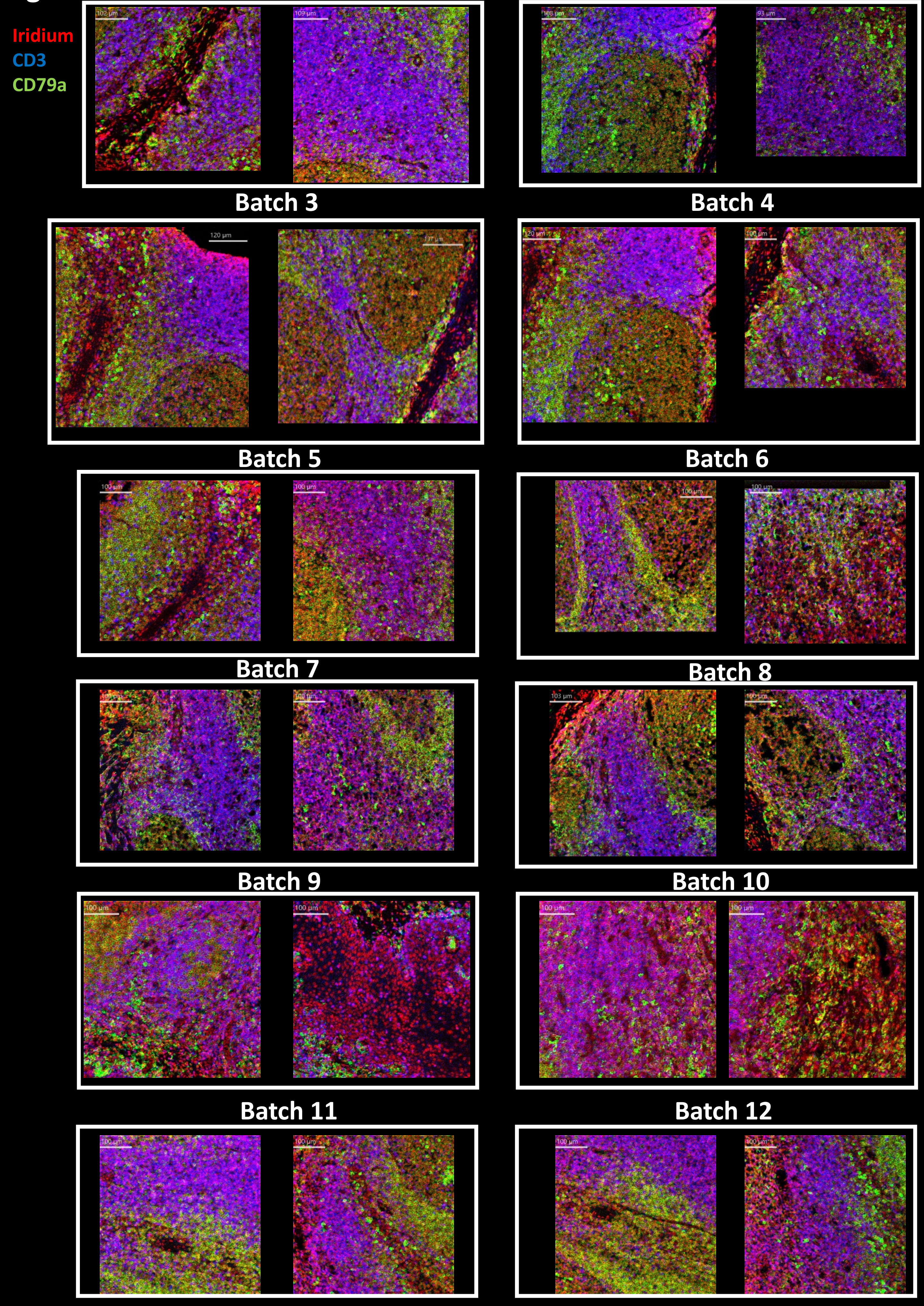
Iridium
CD3
CD79a
Batch 1
Batch 2
Batch 3
Batch 4
Batch 5
Batch 6
Batch 7
Batch 8
Batch 9
Batch 10
Batch 11
Batch 12

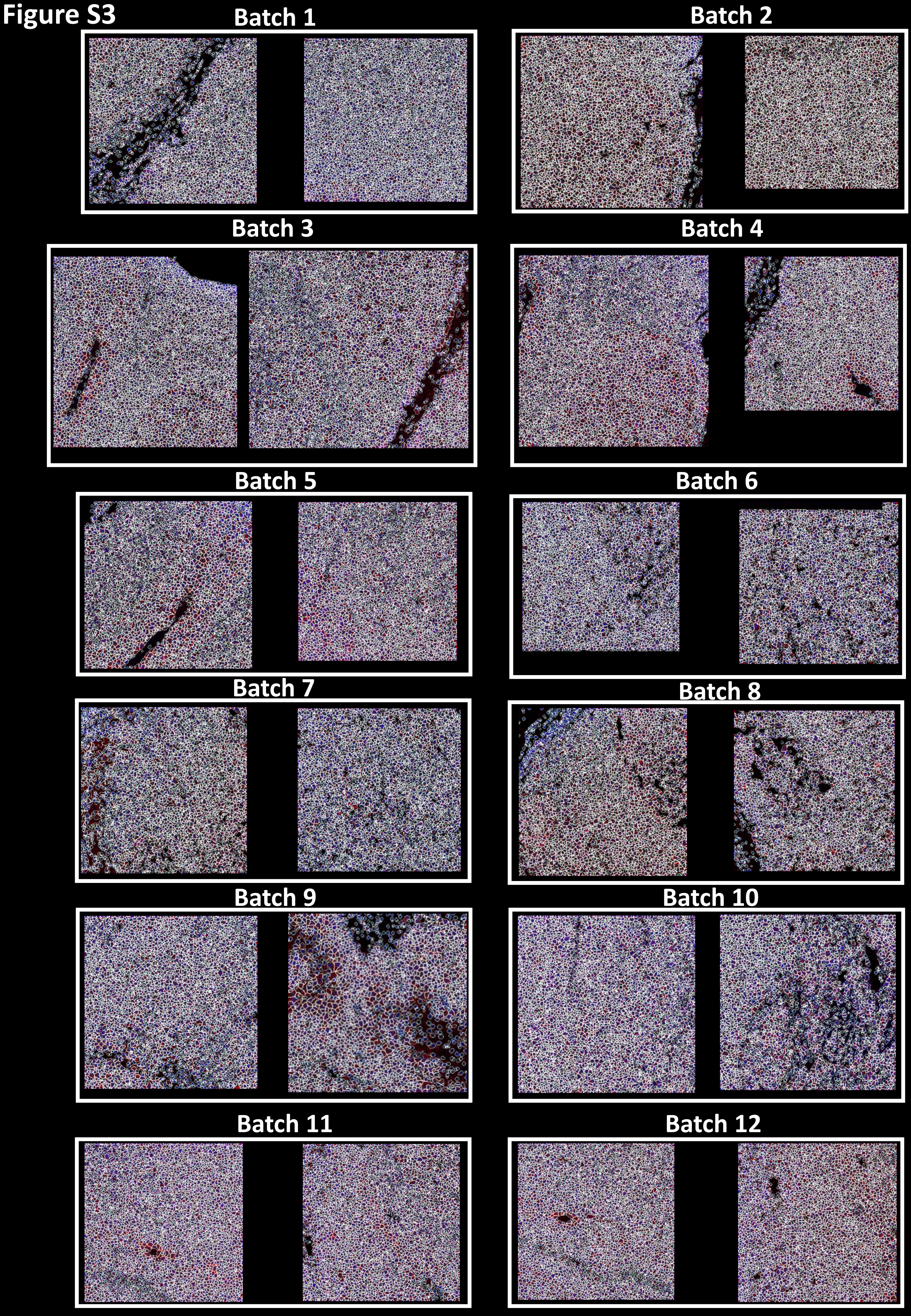
Figure S3
Batch 1
Batch 2
Batch 3
Batch 4
Batch 5
Batch 6
Batch 7
Batch 8
Batch 9
Batch 10
Batch 11
Batch 12

Figure S4

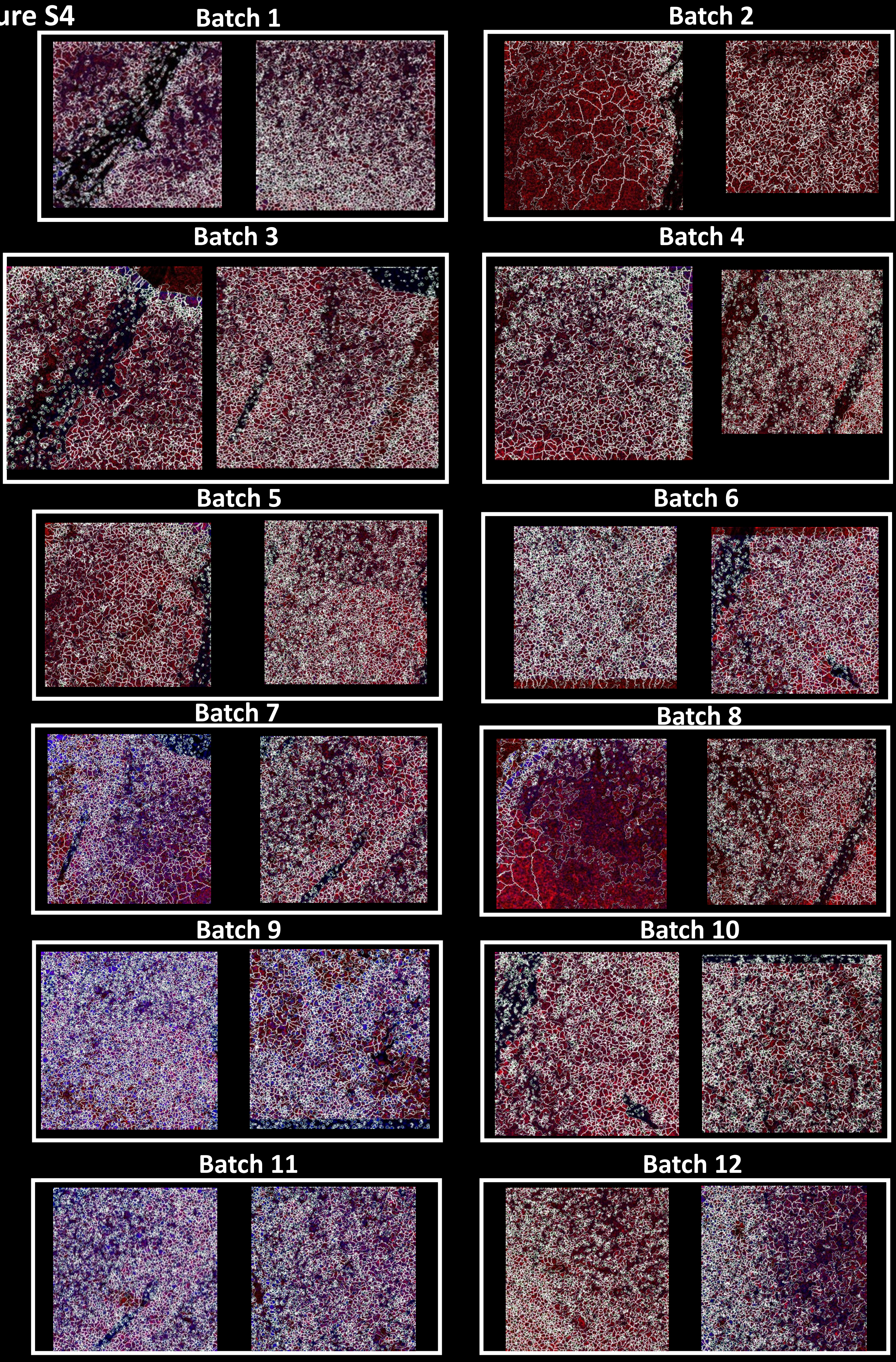

Batch 1
Batch 2
Batch 3
Batch 4
Batch 5
Batch 6
Batch 7
Batch 8
Batch 9
Batch 10
Batch 11
Batch 12

### Figure S5

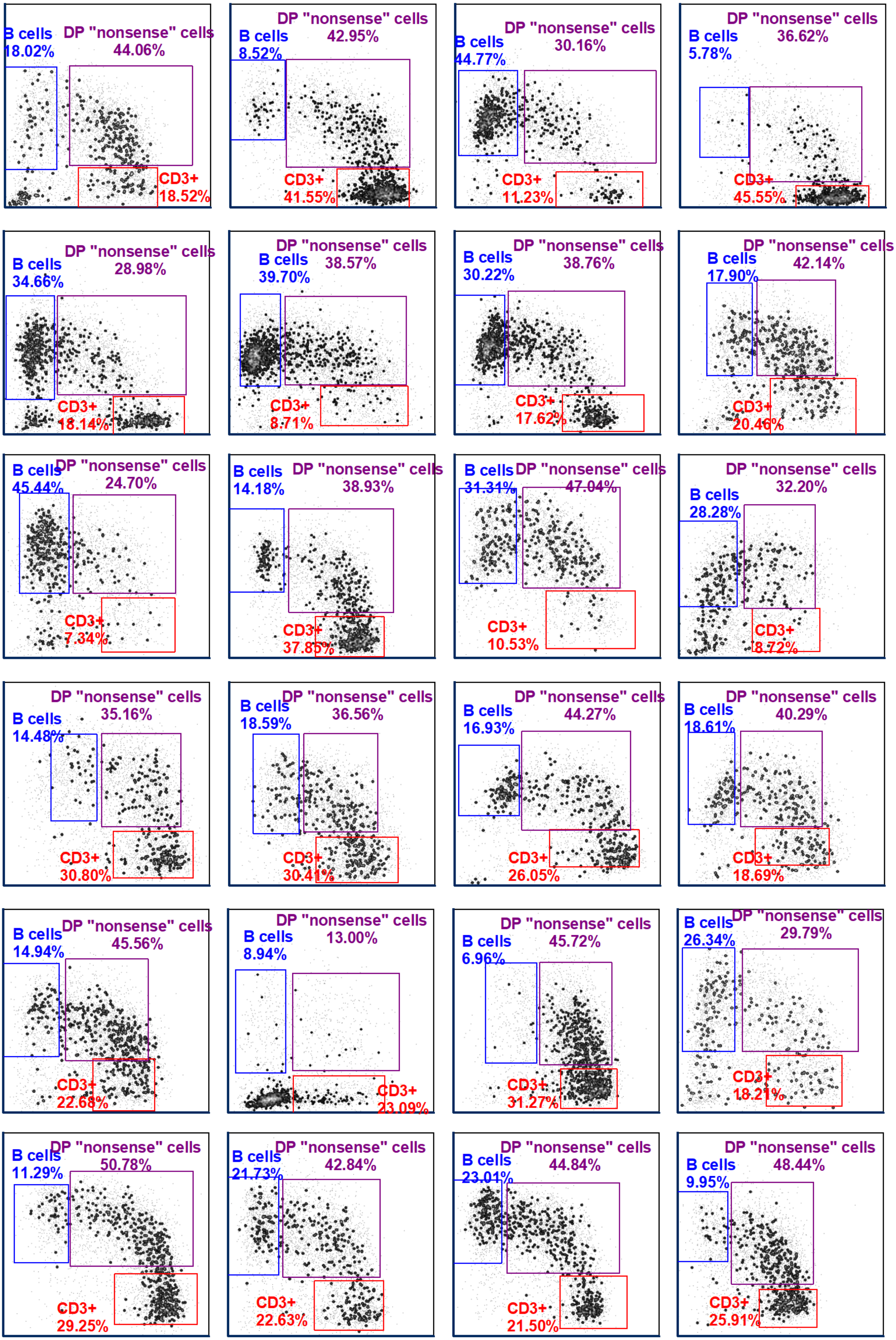

B cells 18.02%
DP "nonsense" cells 44.06%
CD3+ 18.52%
DP "nonsense" cells 42.95%
B cells 8.52%
CD3+ 41.55%
DP "nonsense" cells 30.16%
B cells 44.77%
CD3+ 11.23%
DP "nonsense" cells 36.62%
B cells 5.78%
CD3+ 45.55%
DP "nonsense" cells 28.98%
B cells 34.66%
CD3+ 18.14%
DP "nonsense" cells 38.57%
B cells 39.70%
CD3+ 8.71%
DP "nonsense" cells 38.76%
B cells 30.22%
CD3+ 17.62%
DP "nonsense" cells 42.14%
B cells 17.90%
CD3+ 20.46%
B cells 45.44%
DP "nonsense" cells 24.70%
CD3+ 7.34%
B cells 14.18%
DP "nonsense" cells 38.93%
CD3+ 37.85%
B cells 31.31%
DP "nonsense" cells 47.04%
CD3+ 10.53%
DP "nonsense" cells 32.20%
B cells 28.28%
CD3+ 8.72%
DP "nonsense" cells 35.16%
B cells 14.48%
CD3+ 30.80%
B cells 18.59%
DP "nonsense" cells 36.56%
CD3+ 30.41%
DP "nonsense" cells 44.27%
B cells 16.93%
CD3+ 26.05%
DP "nonsense" cells 40.29%
B cells 18.61%
CD3+ 18.69%
B cells 14.94%
DP "nonsense" cells 45.56%
CD3+ 22.68%
DP "nonsense" cells 13.00%
B cells 8.94%
CD3+ 23.09%
DP "nonsense" cells 45.72%
B cells 6.96%
CD3+ 31.27%
B cells 26.34%
DP "nonsense" cells 29.79%
CD3+ 18.21%
B cells 11.29%
DP "nonsense" cells 50.78%
CD3+ 29.25%
B cells 21.73%
DP "nonsense" cells 42.84%
CD3+ 22.63%
B cells 23.01%
DP "nonsense" cells 44.84%
CD3+ 21.50%
DP "nonsense" cells 48.44%
B cells 9.95%
CD3+ 25.91%

### Figure S6

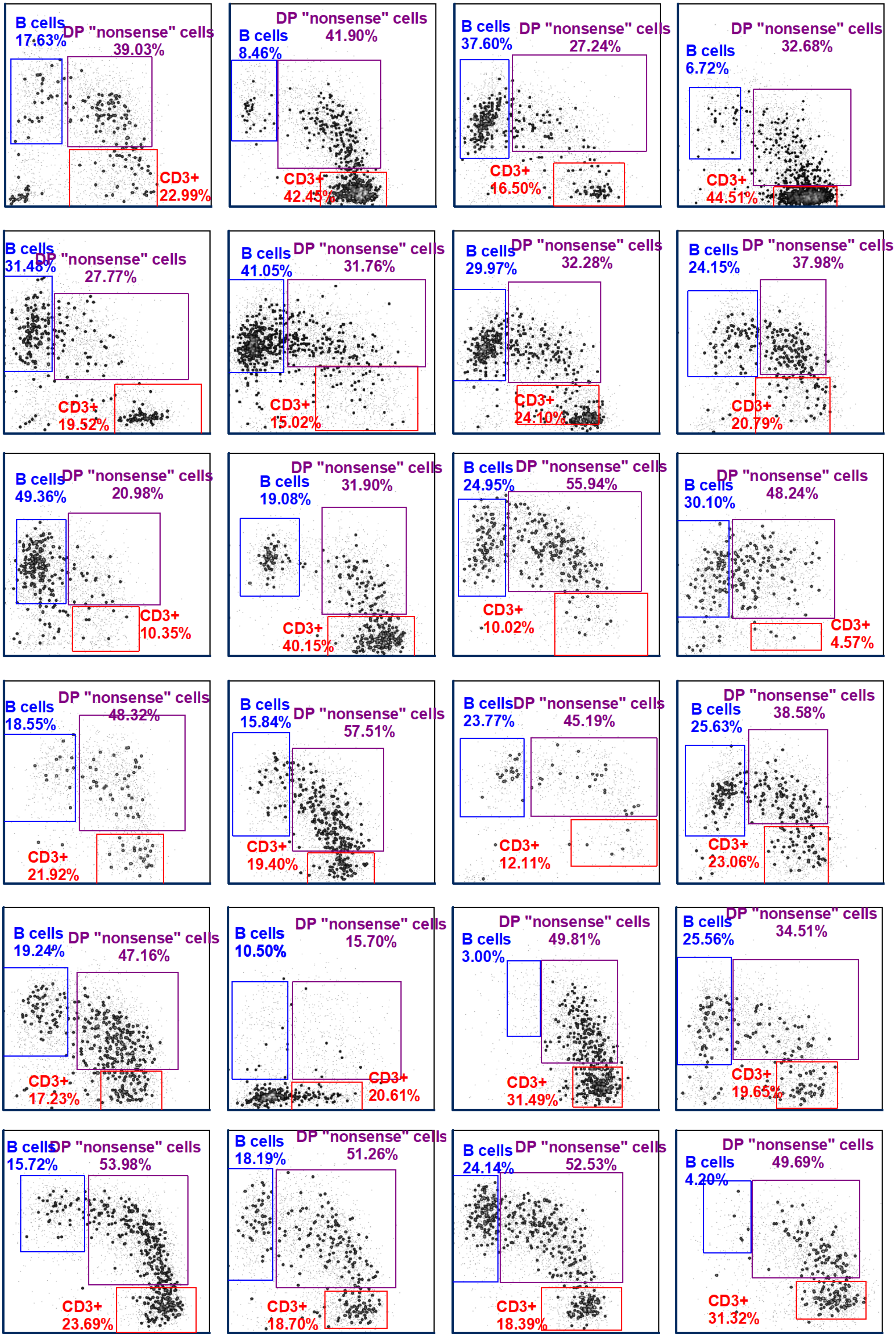
B cells 17.63%
DP "nonsense" cells 39.03%
CD3+ 22.99%
DP "nonsense" cells 41.90%
B cells 8.46%
CD3+ 42.45%
B cells 37.60%
DP "nonsense" cells 27.24%
CD3+ 16.50%
DP "nonsense" cells 32.68%
B cells 6.72%
CD3+ 44.51%
B cells 31.48%
DP "nonsense" cells 27.77%
CD3+ 19.52%
B cells 41.05%
DP "nonsense" cells 31.76%
CD3+ 15.02%
B cells 29.97%
DP "nonsense" cells 32.28%
CD3+ 24.10%
B cells 24.15%
DP "nonsense" cells 37.98%
CD3+ 20.79%
B cells 49.36%
DP "nonsense" cells 20.98%
CD3+ 10.35%
DP "nonsense" cells 31.90%
B cells 19.08%
CD3+ 40.15%
B cells 24.95%
DP "nonsense" cells 55.94%
CD3+ 10.02%
B cells 30.10%
DP "nonsense" cells 48.24%
CD3+ 4.57%
B cells 18.55%
DP "nonsense" cells 48.32%
CD3+ 21.92%
B cells 15.84%
DP "nonsense" cells 57.51%
CD3+ 19.40%
B cells 23.77%
DP "nonsense" cells 45.19%
CD3+ 12.11%
DP "nonsense" cells 38.58%
B cells 25.63%
CD3+ 23.06%
B cells 19.24%
DP "nonsense" cells 47.16%
CD3+ 17.23%
B cells 10.50%
DP "nonsense" cells 15.70%
CD3+ 20.61%
DP "nonsense" cells 49.81%
B cells 3.00%
CD3+ 31.49%
B cells 25.56%
DP "nonsense" cells 34.51%
CD3+ 19.65%
B cells 15.72%
DP "nonsense" cells 53.98%
CD3+ 23.69%
B cells 18.19%
DP "nonsense" cells 51.26%
CD3+ 18.70%
B cells 24.14%
DP "nonsense" cells 52.53%
CD3+ 18.39%
DP "nonsense" cells 49.69%
B cells 4.20%
CD3+ 31.32%

### Figure S7

| Arcsinh c.f.1 only | Arcsinh c.f.1 then 0_1 normalisation | Arcsinh c.f.1 then Z score normalisation |
|---|---|---|

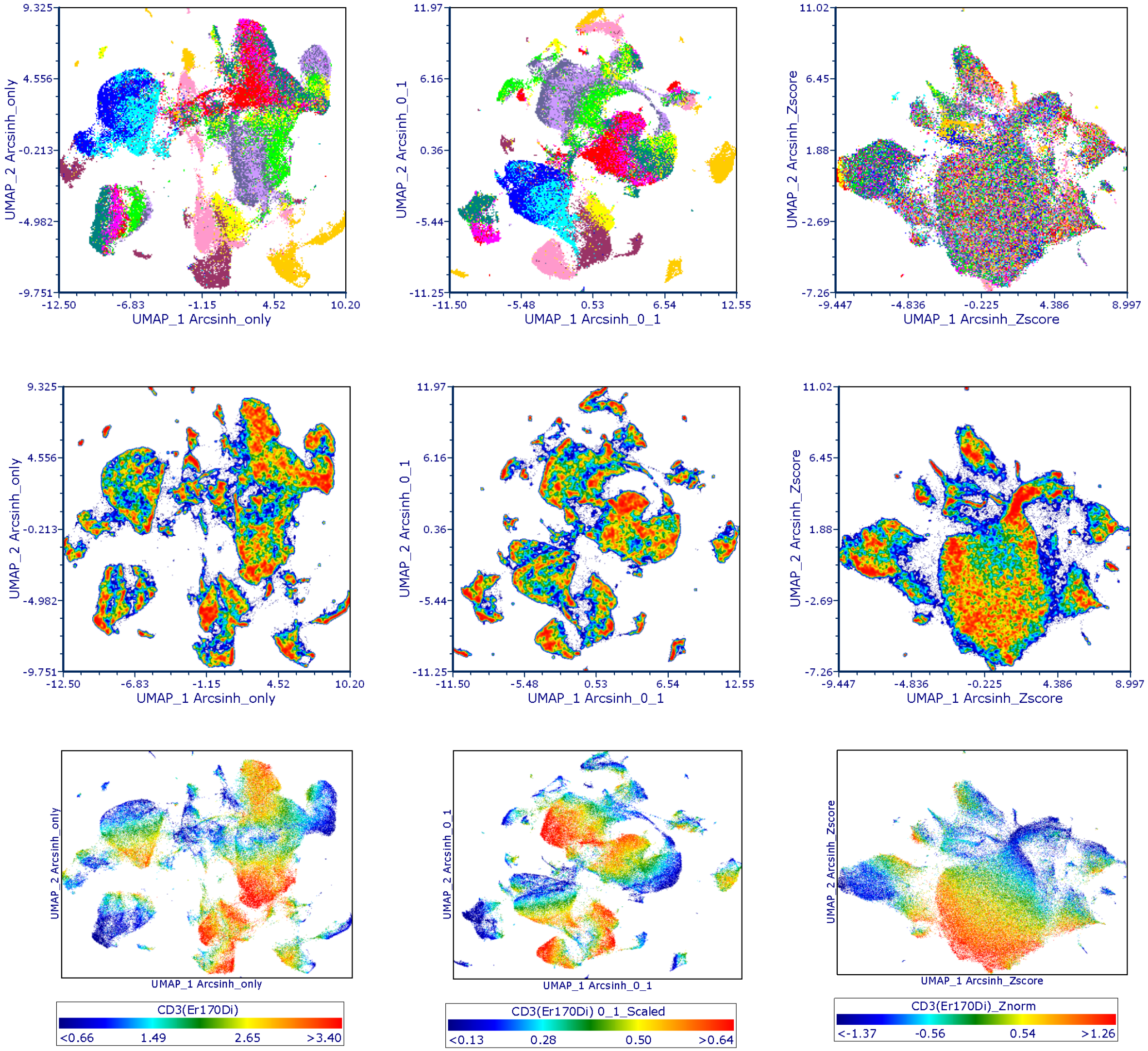


| Gate color | Gate | # of Events | % of all cells |
|---|---|---|---|
| n/a | None | 109535 | 100.00 |
| | Batch_1 | 9949 | 9.08 |
| | Batch_2 | 8505 | 7.76 |
| | Batch_3 | 12582 | 11.49 |
| | Batch_4 | 10515 | 9.60 |
| | Batch_5 | 8815 | 8.05 |
| | Batch_6 | 7830 | 7.15 |
| | Batch_7 | 8218 | 7.50 |
| | Batch_8 | 7953 | 7.26 |
| | Batch_9 | 7702 | 7.03 |
| | Batch_10 | 9095 | 8.30 |
| | Batch_11 | 9216 | 8.41 |
| | Batch_12 | 9150 | 8.35 |

Figure S8

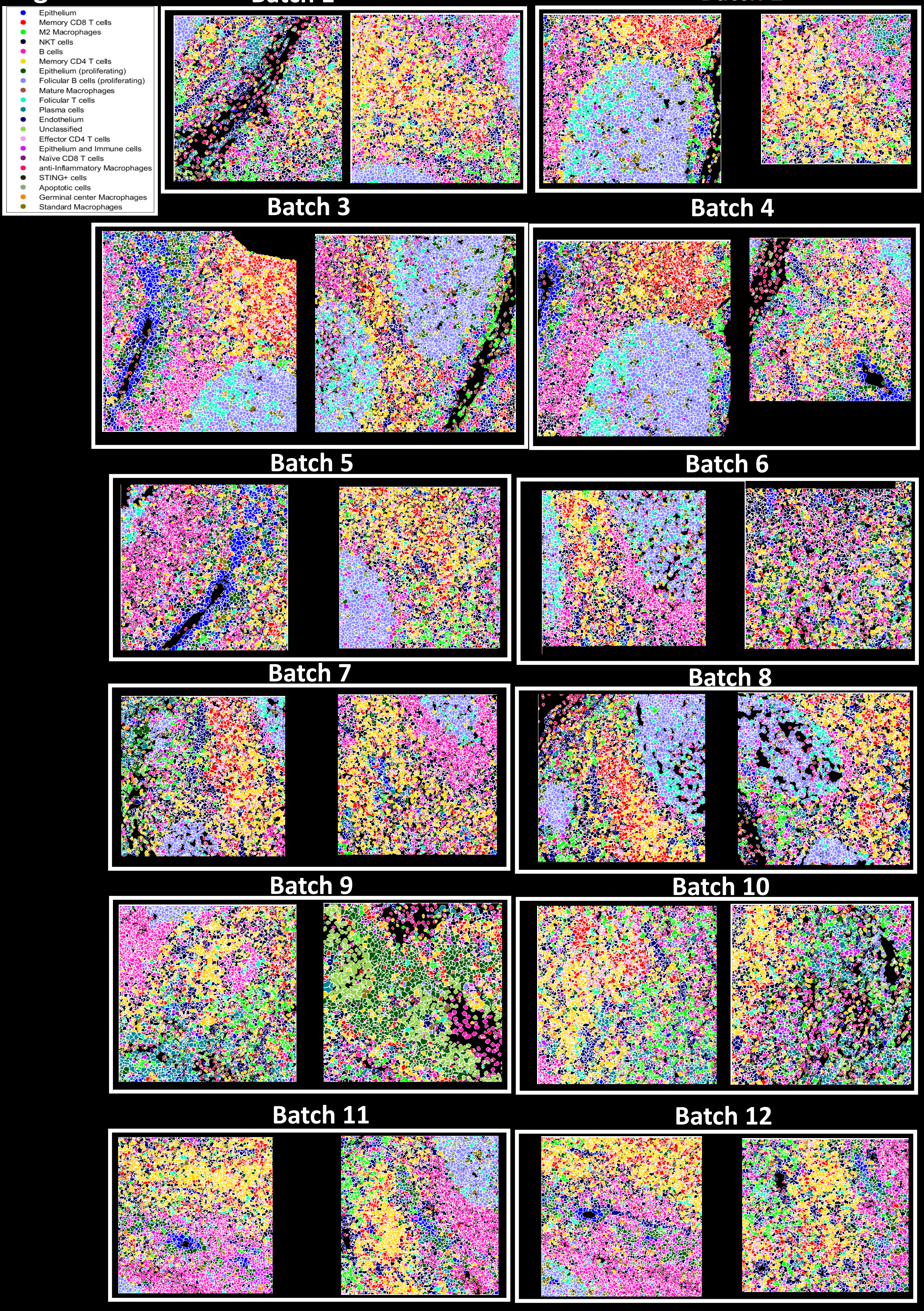
Epithelium
Memory CD8 T cells
M2 Macrophages
NKT cells
B cells
Memory CD4 T cells
Epithelium (proliferating)
Folicular B cells (proliferating)
Mature Macrophages
Folicular T cells
Plasma cells
Endothelium
Unclassified
Effector CD4 T cells
Epithelium and Immune cells
Naïve CD8 T cells
anti-Inflammatory Macrophages
STING+ cells
Apoptotic cells
Germinal center Macrophages
Standard Macrophages
Batch 1
Batch 2
Batch 3
Batch 4
Batch 5
Batch 6
Batch 7
Batch 8
Batch 9
Batch 10
Batch 11
Batch 12

### Figure S9

A

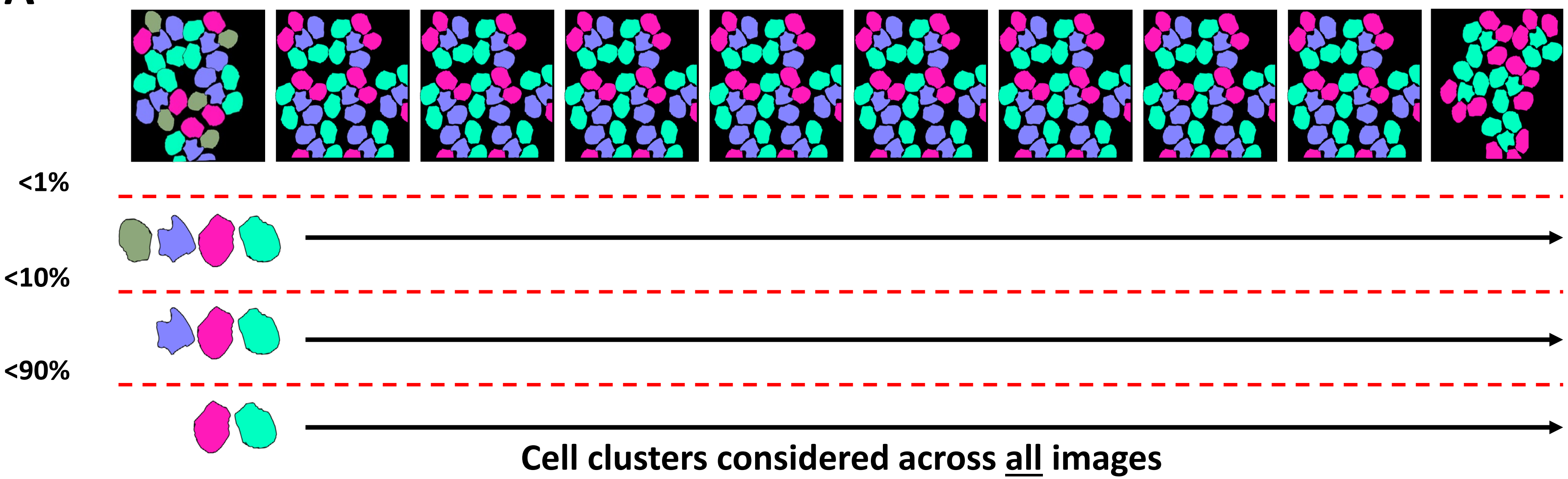


B

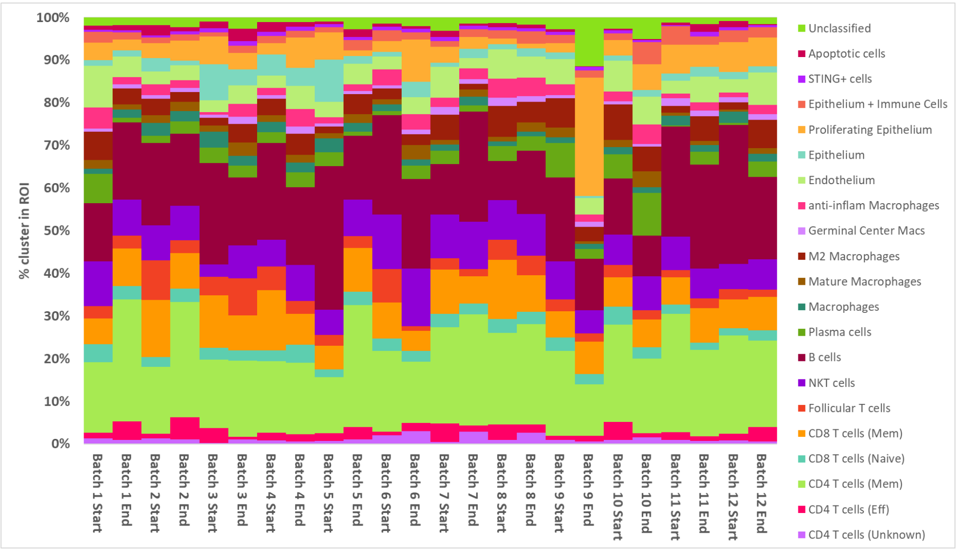


C

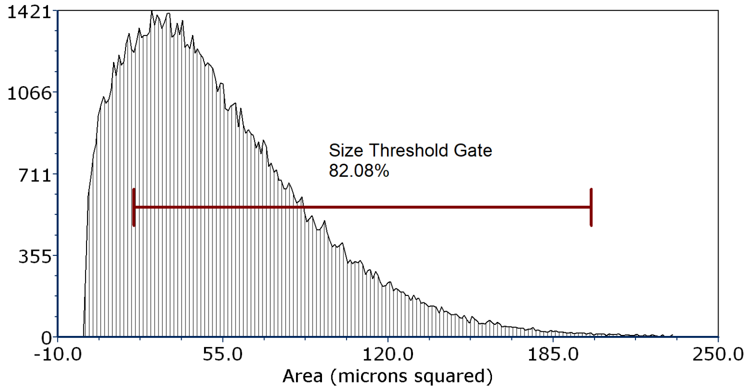

### Figure S10

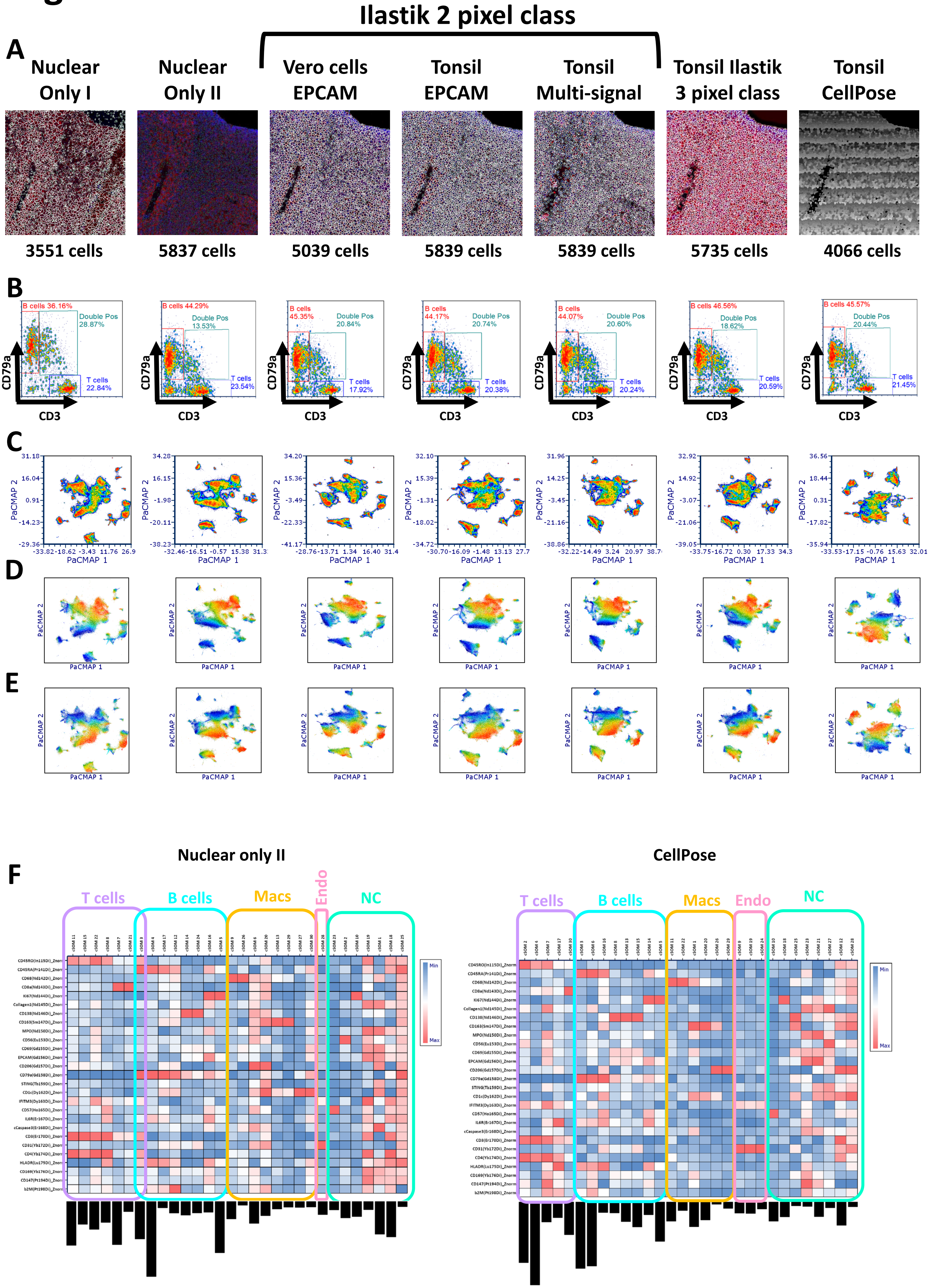
Ilastik 2 pixel class
A
Nuclear Only I
Nuclear Only II
Vero cells EPCAM
Tonsil EPCAM
Tonsil Multi-signal
Tonsil Ilastik 3 pixel class
Tonsil CellPose
3551 cells
5837 cells
5039 cells
5839 cells
5839 cells
5735 cells
4066 cells
B
B cells 36.16%
Double Pos 28.87%
T cells 22.84%
B cells 44.29%
Double Pos 13.53%
T cells 23.54%
B cells 45.35%
Double Pos 20.84%
T cells 17.92%
B cells 44.17%
Double Pos 20.74%
T cells 20.38%
B cells 44.07%
Double Pos 20.60%
T cells 20.24%
B cells 46.56%
Double Pos 18.62%
T cells 20.59%
B cells 45.57%
Double Pos 20.44%
T cells 21.45%
CD79a
CD3
C
PaCMAP 1
PaCMAP 2
D
E
F
Nuclear only II
CellPose
T cells
B cells
Macs
Endo
NC
Min
Max
